# Climbing invasions or climatic refugees: how many and to which extent non-native plant species could reach the top of the Pyrenees mountains under climate change?

**DOI:** 10.64898/2026.01.01.697276

**Authors:** Noèmie Collette, Sébastien Pinel, Anaïs Gibert, Valérie Delorme-Hinoux, Joris A.M. Bertrand

**Affiliations:** Laboratoire Génome et Développement des Plantes (LGDP), UMR 5096, Université de Perpignan Via Domitia (UPVD) - Centre National de la Recherche Scientifique (CNRS), Perpignan, France; Centre de Formation et de Recherche sur les Environnements Méditerranéens (CEFREM), UMR 5110, Université de Perpignan Via Domitia (UPVD) - Centre National de la Recherche Scientifique (CNRS), Perpignan, France

**Keywords:** Climate change, Invasion risk, Invasive species, Mountain, Range shifts, Species Distribution Models

## Abstract

Under climate change, the status of invasive species is becoming complex, as taxa considered invasive in one region may become threatened in another, experiencing simultaneous expansion and decline across their range. In this context, mountain systems may act both as climatic refuges and/or as out-of-reach areas for these species. The Pyrenees, a longitudinal barrier separating the Iberian Peninsula from the rest of Europe, are increasingly exposed to alien plant species spreading from lowlands. Several invasive species have already reached high elevations in the massif, raising concerns about how many more may follow and to what extent under future climate change.

This study assesses future invasion risks for 46 non-native plant species recognized as invasive on both sides of the Pyrenees. We project future dynamics of their bioclimatic niche suitability to identify which are likely to become widespread and where by 2100. Combining occurrence data with 19 bioclimatic variables at 1×1 km resolution, we model current bioclimatic niches and project its future distribution using an ensemble framework integrating five algorithms under four climate scenarios covering the 2021–2100 period.

Projections indicate that 57 % of invasive plant species in the Pyrenees would not be climatic winners under future warming. The centroid of suitable climatic conditions is projected to shift upslope by an average of 258 m across species. This upward movement would tend to stall around 2,000 m, as suggested by future hotspot patterns, effectively squeezing suitable areas and constraining future invasions within mountain zones, as climatic pressure increases at lower elevations. Such redistribution would not necessarily imply negative outcomes, and overlapping with endemic species would remain very limited.

**Synthesis:** Climate change may alter the ecological status of invasive species in mountain systems. In the Pyrenees, many invasive plants are projected to shift upslope while losing suitable lowland conditions, suggesting that future invasions may involve both redistribution and climatic decline rather than simple range expansion. These results raise a broader question: could today’s invasive species become tomorrow’s mountain flora?

## Introduction

Biological invasions are a defining feature of the Anthropocene (Essl *et al.*, 2011; Seebens *et al.*, 2017). The unprecedented acceleration of international trade (Hulme, 2021) and tourism (Anderson *et al.*, 2015), supported by expanding transportation networks (Hulme, 2009; Mcgrannachan *et al.*, 2021), has intensified species introductions worldwide, while climate change further facilitates the establishment and spread of non-native organisms (Hellmann *et al.*, 2008; Hulme, 2017; Stachowicz *et al.*, 2002; Walther *et al.*, 2009; Willis *et al.*, 2010). Among these, invasive species, defined as established non-native alien taxa that spread and form successive populations beyond their introduction points (Soto *et al.*, 2024) are reshaping ecological communities (Bellard *et al.*, 2016; PYSEK *et al.*, 2020b; VILA *et al.*, 2011). Global syntheses further show that they generate substantial economic costs across all continents (Ahmed *et al.*, 2023; Diagne *et al.*, 2021; Zenni *et al.*, 2021), with no sign of slowing down. However, these costs are unevenly distributed, as damages can outweigh management expenditures (Bradshaw *et al.*, 2024). Together, the growing concern surrounding invasive species and their increasing ecological and economic impacts raises practical questions about which species are most likely to expand and where management attention should be focused (Cuthbert *et al.*, 2022), as climate change may promote invasion in some cases while leading to contrasted outcomes in others (Bellard *et al.*, 2013; 2018; Cao PINNA *et al.*, 2024; Finch *et al.*, 2021; Lopes *et al.*, 2023).

In this global context, invasive vascular plants warrant attention, as they dominate recent introductions worldwide (Haubrock *et al.*, 2023; Seebens *et al.*, 2017). Experimental evidence shows that warming legacies enhance invasive plant growth (Ivison *et al.*, 2025; Lembrechts *et al.*, 2016) while it may simultaneously reduce enemy performance (Zhou *et al.*, 2025), creating synergistic advantages for colonization success. Mountain regions were long considered protected from plant invasions due to climatic constraints and limited propagule pressure (i.e. introduction frequency) (Alexander *et al.*, 2016; Zefferman *et al.*, 2015). However, these filters weaken under anthropogenic pressures (Kueffer *et al.*, 2013; Pauchard *et al.*, 2009; 2016), promoting invasive plant establishment at higher elevations (Larson *et al.*, 2021; Steinbauer *et al.*, 2018). In particular, tourism development and expanding infrastructure (e.g. roads, hiking trails and ropeway corridors), increase the frequency of introductions by improving human accessibility, thus intensifying propagule pressure and facilitating the upslope spread of non-native taxa into montane and subalpine environments (Barros & Pickering, 2014; Hemp, 2008; Koyama *et al.*, 2024; Liedtke *et al.*, 2020). Observations across mountain ranges worldwide document accelerating plant invasions at high elevations (Iseli *et al.*, 2023) accompanied by thermophilization patterns reflecting the progressive recruitment of a low-elevation pool of warm-adapted alien taxa (Gottfried *et al.*, 2012; Hamid *et al.*, 2020; Iseli *et al.*, 2025) as well as high rates of species turnover (Iseli *et al.*, 2023; Kalwij *et al.*, 2015; Zu *et al.*, 2022). However, despite growing evidence of invasive plants spread at high elevations, invasion risk is rarely assessed across entire mountain ranges and multiple species, leaving climate-driven redistributions along elevational gradients poorly understood.

As a major European mountain range, the Pyrenees provide a case study for assessing climate-driven invasion risks in mountain ecosystems. Spanning France, Andorra, and Spain between the Atlantic and Mediterranean regions, this range is located at the crossroads of several climatic and biogeographic influences. It is considered among the top biodiversity hotspots in Europe, with 3652 indigenous vascular plant species, of which about 5% are endemic (GÓMEZ *et al.*, 2017). The Pyrenees have acted both as refugia and dispersal corridors during past climatic fluctuations (GARCÍA *et al.*, 2022; Schmitt, 2009). However, under ongoing climate warming, it remains unclear whether they will primarily function as a biogeographic barrier constraining the further spread of invasive species, emerge as a climatic refuge for species displaced from warmer regions, or become newly colonized environments, fostering novel species assemblages involving native taxa. Most invasive plants in the Pyrenees occur at lower elevations, where human activities and associated anthropogenic corridors have facilitated their introduction and establishment, with valleys acting as the primary entry points into the range (Aymerich & Saez, 2019; Jantzi *et al.*, 2021; Martinez-Fuentes *et al.*, 2025; Ninot *et al.*, 2017). As a transboundary mountain range, it is interconnected by dense transport and tourism networks that facilitate species introductions from multiple biogeographic sources. Although few species are invasive at high elevations in the Pyrenees, factors such as second-home development, intense touristic and commercial connectivity, and ongoing climate warming may foster the future spread of some non-native species (Aymerich & SÁez, 2019; Soto *et al.*, 2025).

Although projections tend to predict an intensification of invasion impacts under future climates (Adhikari *et al.*, 2022; Carboni *et al.*, 2018; Petitpierre *et al.*, 2016), recent evidence suggests that non-native species trajectories may be more complex than uniformly expansive. Range dynamics involve both gains and losses, with some taxa thriving under new conditions (Paudel *et al.*, 2025), while others experience local declines or face extinction risks within their native range (Staude *et al.*, 2025). This challenges the traditional perception of invasive species as systematic “winners” of global change and highlights a dynamic continuum of biogeographic responses shaped by regional climates, human influence, and ecological constraints (Hong *et al.*, 2025). One quarter of naturalized plant species are threatened within their native range (Staude *et al.*, 2025), illustrating that invasion success in one region can coincide with conservation concerns in another (GarzÓn-Machado *et al.*, 2012; MarkovÁ *et al.*, 2020). This paradox, increasingly recognized across taxonomic groups (Robuchon *et al.*, 2025; Tedeschi *et al.*, 2025), raises fundamental questions about whether non-native populations should be conserved, and to what extent naturalization may act as inadvertent *ex-situ* refuges under accelerating global change.

In this context of contrasting invasion trajectories, a spatially explicit assessment of future climatic suitability is required to clarify patterns in mountain systems. Rather than assuming uniform expansion, we hypothesize that future climatic suitability in the Pyrenees will display species-specific and spatially heterogeneous dynamics, driven by interspecific differences in climatic niches and strong environmental gradients within this mountain chain. Consequently, invasion risk requires examining how suitable conditions may expand, persist, or contract across space and time, with direct implications for proactive management efforts and associated economic costs. We therefore used Species Distribution Models (SDM) to assess the dynamics of future bioclimatic suitability of 46 non-native plant species classified as invasive on both sides of the Pyrenean range, documenting species-specific potential shifts and identifying areas most likely to become vulnerable to invasion by 2100 under multiple climate scenarios. This study provides a rare, range-wide, spatially explicit assessment of climate-driven invasion risks across an entire mountain chain, revealing contrasting invasion trajectories and where invasion risk is likely to persist, shift, or emerge under future climates.

Spatially explicit tools, such as species distribution models (SDMs), provide a well-suited framework to address these objectives. SDMs are widely used in invasion ecology to project how climate change may expand suitable areas to non-native species (Barbet-Massin *et al.*, 2018; Renteria *et al.*, 2021; Srivastava *et al.*, 2019). Here, we employ an ensemble SDM framework to characterize large-scale patterns of climatic suitability and to identify where and for which species environmental conditions are becoming increasingly permissive to invasion under ongoing warming. By focusing on species-specific dynamics along broad climatic gradients, our analyses allow consistent comparisons of invasion risk across species and regions and provide an operational basis for anticipating future spread and guiding surveillance and preventive management efforts.

## Material and methods

### Study area, species occurrences and bioclimatic data

This study follows the mountain delimitation of Körner *et al*. (2011), updated by Snethlage *et al*. (2022). This framework defines mountain systems using topographic ruggedness and bioclimatic criteria, ensuring a standardized and ecologically meaningful delineation. Within this framework, the Pyrenees span ∼40,000 km² along the France-Spain-Andorra border (Figure 1), with over 150 peaks above 3000 m (GÓMEZ *et al.*, 2017). The range shows a marked asymmetry, with steep northern slopes and broader southern valleys, and lies at the interface of Atlantic, Mediterranean, and continental climatic influences, resulting in strong environmental heterogeneity. Mean annual temperatures range from >10 °C in lowlands to <4 °C at high elevations, while precipitation varies from >2,000 mm on Atlantic-facing slopes to ∼1,000 mm in more sheltered areas (Cuadrat *et al.*, 2024). The Pyrenees lie at the transition between the Eurosiberian and Mediterranean biogeographical regions, forming a major ecotonal zone where overlapping climatic influences generate strong environmental gradients and contribute to high biodiversity.

**Figure 1.**
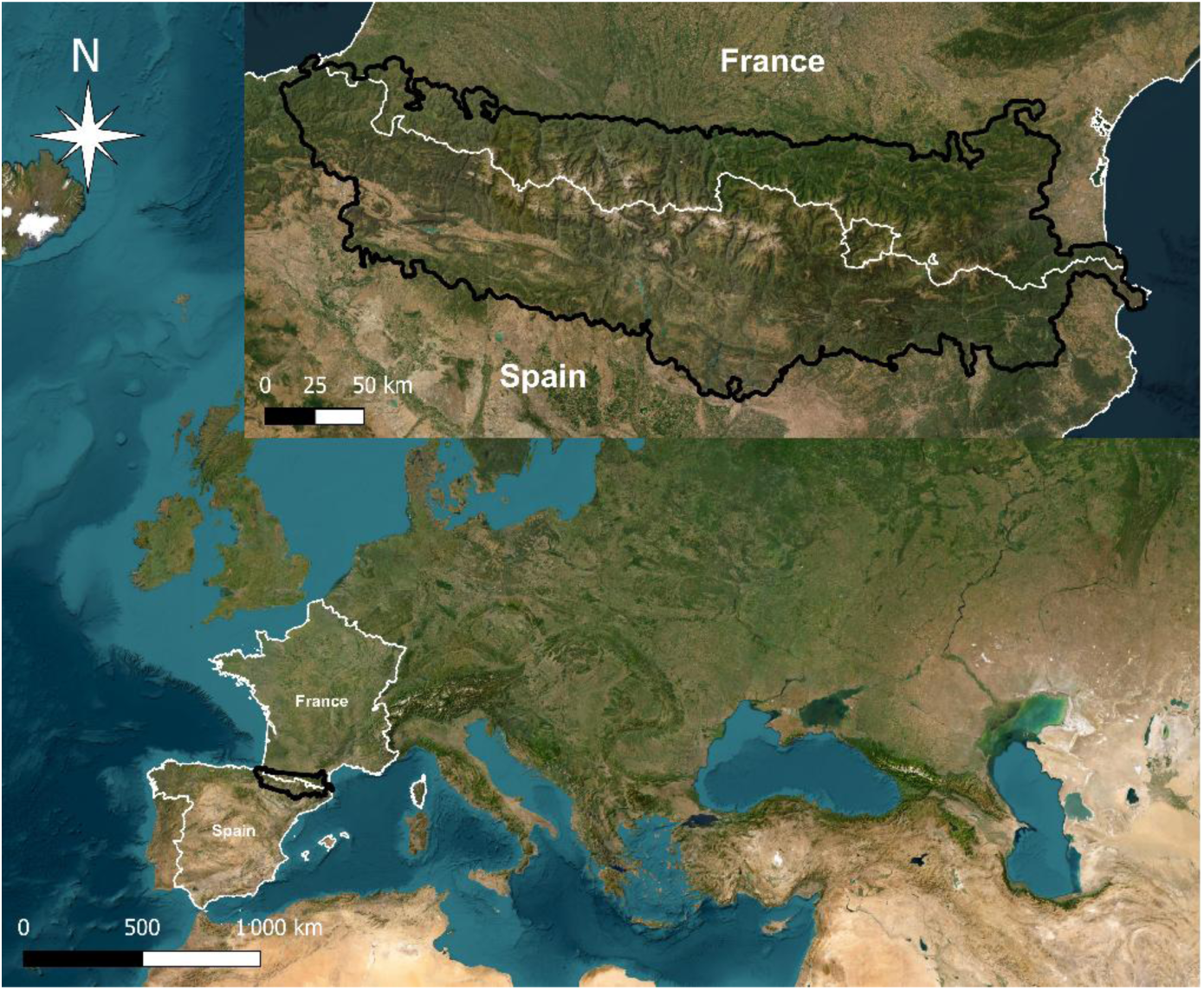
Geographic location of the Pyrenean mountain range in Europe. The Pyrenees are shown in their European context and in detail with the mountain boundary highlighted in black (top panel), following the Global Mountain Biodiversity Assessment definition (Körner *et al*., 2011, updated by Snethlage *et al*., 2022). National boundaries are indicated in white. Maps are displayed in WGS84 (EPSG:4326). Background imagery © Esri, *Esri World Imagery*.

We focused on plant species listed as invasive in official lists or authoritative catalogues, with a major impact on at least one regional list of areas adjacent to the Pyrenees (i.e. widely distributed taxa forming dense populations, according to French classifications, see Appendix S1-A in Supporting Information) All selected species occur in both France and Spain, ensuring their relevance across the Pyrenean range. The official list of invasive species in Andorra (Appendix S1-A), whose territory lies entirely within the Pyrenees, was used to confirm species presence and inform the discussion at the mountain-range scale.

For each species, post-1970 occurrence data were obtained from the Global Biodiversity Information Facility (doi: 10.5281/zenodo.18114751), including all available records worldwide. To limit spatial sampling bias, records with coordinate uncertainty > 1 km were removed, and spatial thinning was applied to retain a single occurrence per grid cell. Species lacking GBIF occurrence records in regions adjacent to the Pyrenees or species with insufficient occurrences to ensure representation across all spatial blocks required by the spatial cross-validation scheme were excluded. After filtering, 46 species remained out of the 50 initially identified (see Appendix S2).

Bioclimatic predictors were obtained from the WorldClim v2.1 database (Fick & Hijmans, 2017), including 19 temperature and precipitation-related variables for the baseline period (1970–2000) and four future periods (2021–2040, 2041–2060, 2061–2080, and 2081–2100, referred to as 2030, 2050, 2070, and 2090). Future projections were generated under four Shared Socioeconomic Pathways (SSPs, SSP126, SSP245, SSP370, and SSP585), covering a range of potential climate trajectories (O’NEILL *et al.*, 2017).

### Distribution of invasive species across the altitudinal belts of the Pyrenees and characteristics

Occurrence records were intersected with four elevation belts corresponding to the main vegetation zones of the Pyrenees: foothills and lower montane (0–900 m), montane (901–1 800 m), subalpine (1 801–2 300 m), and alpine zone above 2 300 m (Saule *et al.*, 2018). This analysis used the full set of occurrence records (i.e., non–spatially thinned data). For each belt, we quantified species richness, species composition and relative proportions, and summarized species composition by life form, using data from the FloraVeg.EU database (ChytrÝ *et al.*, 2024). Life forms were harmonized into four categories: trees, shrubs, herbs, and aquatic species (Appendix S2).

A database was compiled to characterize the invasion history, biogeographic origin and ecological attributes of invasive species. From Plants of the World Online (Powo, 2025) we extracted native and introduced distributions (WGSRPD level-3), biome affiliation and extinction risk (Bachman *et al.*, 2024). Native and introduced ranges were quantified as the percentage of WGSRPD regions occupied by each species and their sum was used as an estimate of total geographic coverage. Primary introduction pathways, introduction period in Europe and the earliest occurrence records in Europe, France and Spain were compiled from EASIN (European Commission, Joint Research Centre, 2025). Life span, life form, flowering duration, dispersal mode and distance were compiled from FloraVeg.EU database. Species presence in the central Pyrenees (Andorra) was additionally included to account for the ability of species to occur within the core of the mountain range. A full list of compiled invasion attributes is provided in Appendix S2. To relate species attributes to their projected bioclimatic trajectories, each taxon was classified according to whether it showed a consistent increase (“winner”) or display decrease (“loser”) in suitable bioclimatic areas by the end of the century across SSPs scenarios. Differences between winners and losers were assessed separately for numeric and categorical traits, with the null hypothesis that trait distributions did not differ between both groups. Numeric variables describing geographic coverage, flowering duration and introduction chronology were analyzed using Wilcoxon rank-sum tests, whereas categorical variables related to biome affiliation, life history, dispersal strategy, introduction pathways and biogeographic origin were analyzed using chi-square tests or Fisher’s exact tests when expected cell counts were < 5.

### Bioclimatic suitability modeling and spatial dynamics of invasion risk

#### Bioclimatic suitability modeling

We adopted the ensemble modeling framework described in Collette *et al.*, 2026. Because invasive plants are typically widespread, occurrence data are less imbalanced; model performance was therefore evaluated using the Area Under the Receiver Operating Characteristic Curve (AUC-ROC), rather than the Area Under the Precision–Recall Curve (AUC-PR) which is more appropriate for taxa with more imbalanced data. The spatial cross-validation scheme was adjusted by increasing the number of spatial blocks to five, with an adjusted block size. These study-specific methodological adaptations are detailed in the ODMAP protocol (Overview, Data, Model, Assessment and Prediction), a standardized framework for transparent SDMs reporting (Zurell *et al.*, 2020, see Appendix S1-B).

Model calibration was performed within the smallest bounding box maximizing GBIF data coverage while maintaining computational efficiency. This extent included regions spanning a wide range of environmental conditions (e.g., northern Europe and the Alps) thereby ensuring broad environmental representation. This approach is consistent with evidence that models calibrated on invaded ranges outperform those built across entire distributions (Barbet-Massin *et al.*, 2018). Future projections were restricted to a bounding box encompassing the Pyrenean range, and all suitability maps for analyses were subsequently cropped to the Pyrenees following the mountain delimitation proposed by KÖRNER *et al.* (2011) and updated by Snethlage *et al.* (2022) in WGS84 (EPSG:4326) (Figure 1). Maps are available in Appendix S1-C.

While these results illustrate the use of bioclimatic projections to explore potential climate-driven shifts in invasion risk, their interpretation requires careful consideration of their scope and limitations (Collette *et al.*, 2025; Fenollosa *et al.*, 2025; Gallien *et al.*, 2015; Srivastava *et al.*, 2019). Accordingly, projected changes in climatic suitability should be interpreted as indicators of potential establishment of non-native species rather than direct predictors of future abundance or realized spread and must be interpreted alongside broader ecological and socio-environmental processes.

#### Spatial and temporal dynamics of bioclimatic suitable area

To quantify changes in bioclimatic suitability, continuous suitability maps were binarized for each period and SSP using the maximum sum of sensitivity and specificity (maxSSS) threshold (Liu *et al.*, 2013; 2016). Suitable area was calculated for the current period and future period–SSP combination. Spatial overlap between current and future binary maps was computed to estimate the proportion of currently suitable areas projected to remain suitable under each scenario. Each taxon was classified according to whether it showed an increase (“winner”) or decrease (“loser”) in suitable bioclimatic areas by the end of the century across all SSP scenarios. Latitudinal, longitudinal and elevational shifts between the current period and 2081–2100 were quantified by overlaying binarized projections with elevation data from Cgiar-CSI (2018; ∼1 × 1 km resolution at the equator) and geographic coordinates. For each species, elevational, latitudinal and longitudinal centroids, defined as the mean elevation, were calculated for the current period and for each SSP. Only species retaining suitable areas under all SSPs were used to estimate overall centroid shifts between present and future conditions. Species-specific shifts under each SSP are reported in Appendix S1-D, supporting information.

*Mapping bioclimatic invasion hotspots and their overlap with endemic biodiversity*.

Continuous suitability maps for the current and future (2081–2100) periods were averaged across species to identify bioclimatic hotspots, defined as areas where the highest number of invasive taxa is projected to find suitable climatic conditions. To assess potential spatial conflicts between biological invasions and native biodiversity, invasive-species hotspots were compared with projected bioclimatic hotspots previously identified for 59 endemic plant species of the Pyrenees (Collette *et al.*, 2026). Both current and future hotspots were derived using a consistent modeling framework, allowing direct comparison of spatial overlap within each period. Hotspot maps of all SSPs were binarized using a threshold of 0.2, corresponding to the lowest non-zero class of the legend and retaining all areas showing any hotspot signal. Spatial overlap between invasive and endemic hotspots was quantified using the Jaccard similarity index (spatial overlap index; (Jaccard, 1901)). This index ranges from 0 (no overlap) to 1 (complete overlap), providing a standardized measure of spatial congruence between invasion risk hotspots and areas of high endemic suitability.

## Results

### A strong decline in invasive species richness with elevation

Intersecting occurrence records with the four elevational belts revealed that invasive were clustered along valleys and low elevations, especially in the northeastern Pyrenees (Figure 2a). The foothill and lower montane belt (0–900 m), which represents 45.8 % of the Pyrenees surface, hosted all the studied species, accounting for 81.5**%** of all occurrences, making it the richest zone (Figure 2b). Species richness then declined steadily with elevation: 35 species in the montane belt (901–1 800 m), 10 species in the subalpine belt (1 801–2 300 m), and 7 species in the upper alpine zone (> 2 300 m). After standardizing by belt area, species density followed the same decreasing trend with elevation (see Appendix S1-E). Seven species are recorded in the alpine zone, *Elaeagnus angustifolia, Fallopia baldschuanica, Helianthus tuberosus, Impatiens balfourii, Parthenocissus inserta, Prunus cerasifera, Senecio inaequidens.* Within the subalpine belt, most species were rare; nine of the 10 species occurred only marginally (0.11–2 % of their occurrences). Only *Elodea canadensis* showed higher representation at this elevation with 8.7 %, indicating greater capacity in colder, high-mountain environments. Overall, invasive plants in the Pyrenees are primarily concentrated at low elevations, although several taxa already show the ability to colonize high-mountain environments.

**Figure 2.**
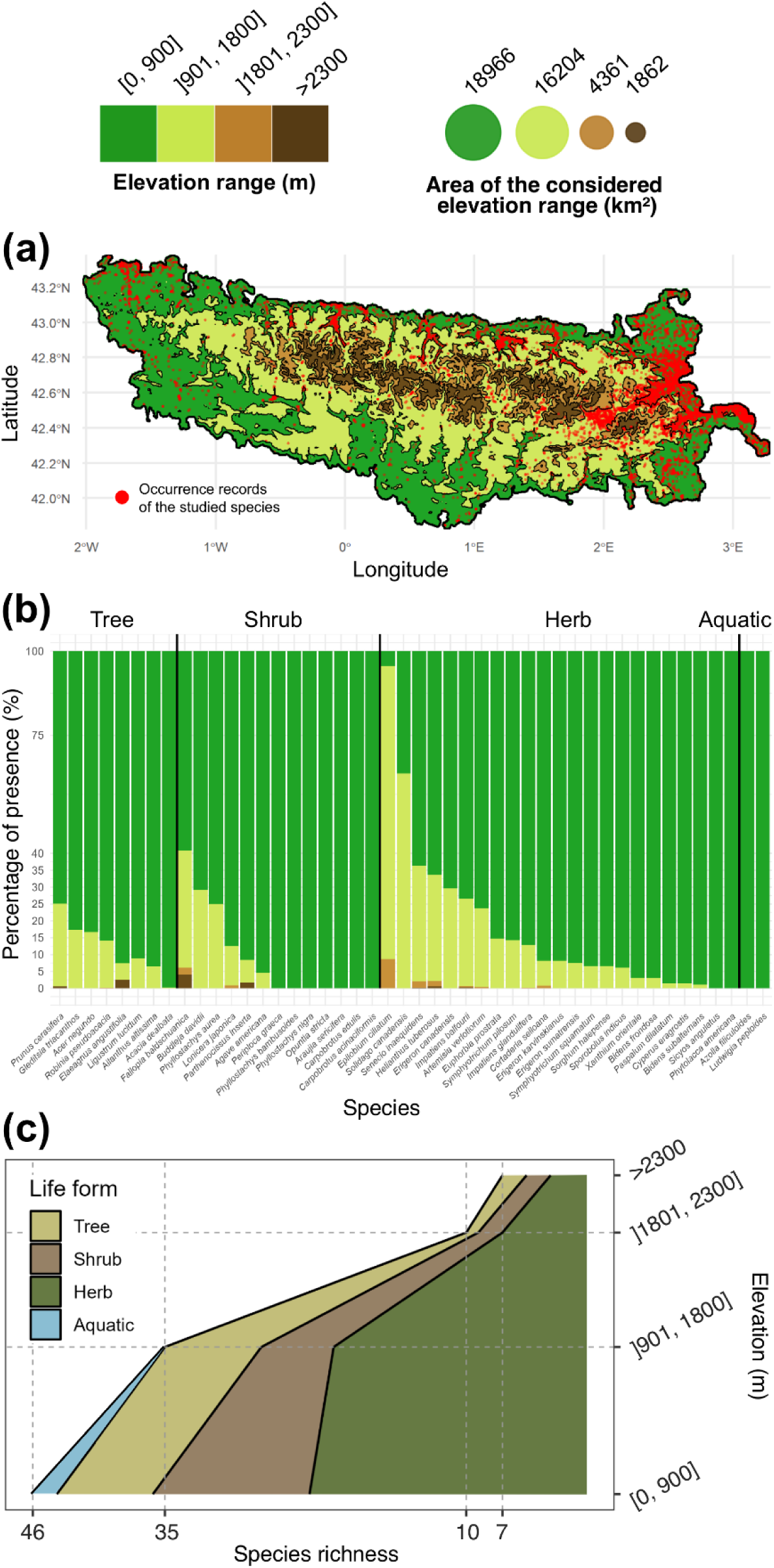
Spatial, elevational and life-form distribution of 46 invasive plant species in the Pyrenees. **(a)** Spatial distribution of all occurrence records across the main vegetation belts of the Pyrenees: foothills and lower montane (0– 900 m), montane (901–1 800 m), subalpine (1 801–2 300 m), and alpine (> 2 300 m). **(b)** Percentage of occurrences per species across the four vegetation belts and their life-form categories (tree, shrub, herb and aquatic). **(c)** Species richness across elevation for each life-form category.

Life-form composition also shifted with elevation (Figure 2c). Herbs dominated all elevational belts, reflecting their broad ecological tolerance. Shrubs and trees were concentrated in the lower and mid-montane belts, with richness declined sharply with elevation. Trees were absent from the upper alpine zone (> 2 300 m), indicating that woody invasive species are currently unable to establish in the most climatically constrained conditions. This consistent dominance of herbs and the limited altitudinal distribution of woody species highlight the strong climatic and habitat filtering along the Pyrenean gradient.

No invasion attribute differed significantly between climatic winners and losers (all p > 0.05; Table S3). However, introduction pathways (p = 0.055) and species presence in the central Pyrenees (p = 0.086) showed marginal associations with projected trajectories. Winners were more frequently associated with escape from confinement pathways, whereas losers were more often linked to release in nature pathways. In addition, species occurring in the central Pyrenees tended to be more frequently classified as losers.

### Future bioclimatic suitability declines for most invasive species, with 47% potential winners

Projections indicated variation in future bioclimatic suitability across the 46 invasive taxa (Figure 3). Overall, 57 % of species were projected to experience a reduction in suitable climatic areas by 2081–2100 under at least one SSP scenario. Losses were substantial for several species: four were projected to lose more than 75 % of their suitable area, seven to lose 50–75 %, one to lose 33–50 %, and fourteen to lose less than 33 %. Averaged across species, changes ranged from +18.7 % under the most optimistic scenario (SSP126; min: +222.6 %, max: −65.2 %) to −75.1 % under the most pessimistic one (SSP585; min: −3.7 %, max: −100 %). The mean positive change observed under optimistic scenarios was driven by several species that, although ultimately classified as “losers” under more pessimistic projections, still showed temporary increases in suitable areas under milder climate scenarios. This pattern was observed for fourteen species in at least one SSP. Two species (*Paspalum dilatatum* and *Elaeagnus angustifolia*) experienced a complete loss of suitable bioclimatic area in at least one scenario. In contrast, twenty species (43 %) were identified as consistent potential “winners” of climate change, with average gains of +388.4 % under SSP126 (min: +13.6 %, max: +1637.2 %) and up to +658.6 % under SSP585 (min: +22.6 %, max: +4178.5 %). For 11 species, projected range gains were greater under the most pessimistic scenario, suggesting a gradient in which suitability increases with the severity of climate change, whereas the opposite pattern was observed for the remaining species.

**Figure 3.**
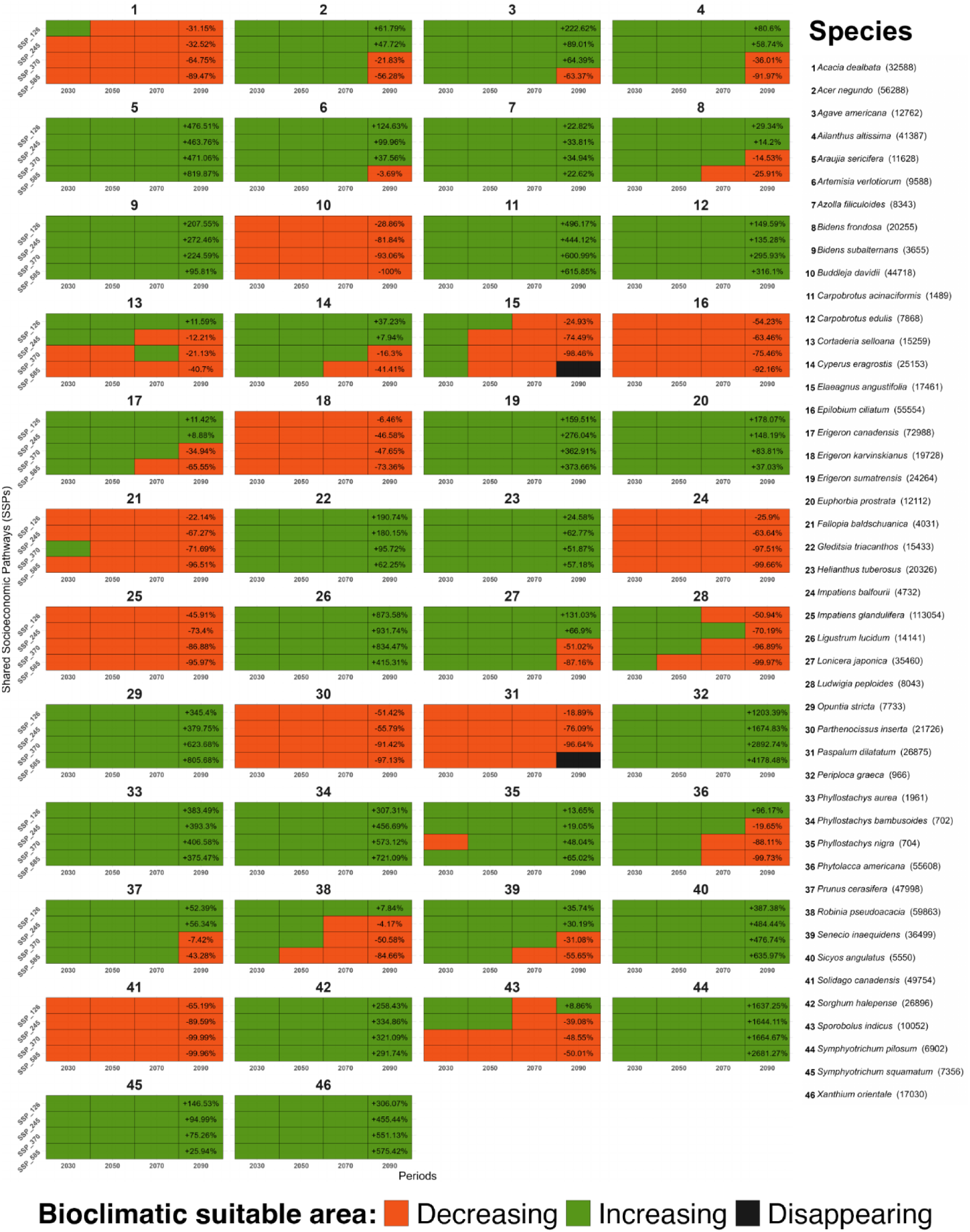
Projected extent changes in bioclimatic suitability for 46 invasive plant species in the Pyrenees. Extent changes in suitable areas over the 2021-2100 period under Shared Socioeconomic Pathways (SSP) SSP126, SSP245, SSP370, SSP585, relative to the baseline period (1970-2000) for 46 invasive plant species. Each box represents the extent change in suitable areas, with percentage values shown for the 2081-2100 period. Orange indicates a reduction in suitable areas, green expansion, and black a complete loss of suitability. 2021-2040, 2041-2060, 2061-2080, 2081-2100 periods designated as 2030, 2050, 2070, 2090.

Across all algorithms (GLM, GAM, GBM, RF, MaxEnt), cross-validated model performance was high (AUC 0.83 ± 0.07; Boyce index 0.83 ± 0.12; sensitivity 0.87 ± 0.05; see Appendix S1-F))

### Climate-driven upward, northward and westward shifts of climatically suitable areas with complete spatial reorganization

Across species retaining suitable conditions across SSPs during the 2081-2100 period, the centroid of climatically suitable areas shifted upward on average by 258 m (range −80 to +1072.7 m), with a mean northward shift of ∼13 km (range −33.2 to +58.9 km) and a western shift of ∼72 km (range −222.3 to +37.2 km) (Figure 4a). Overall, 43 species are projected to shift upslope (Figure 5), with the centroid of their climatically suitable areas moving upward by +2 % to +213.4 % (mean: + 71 %), whereas only one species is projected to shift downslope (*Buddleja davidii*, Figure 5), with centroid elevations downslope by −10.2. The strongest upward shifts were projected for *Epilobium ciliatum* (+ 1033.2 m; from 1164.7 to 2198.0 m), *Impatiens glandulifera* (+ 1072.7 m; from 1096.2 to 2168.9 m) and *Solidago canadensis* (+ 784.2 m; from 1214.1 to 1998.3 m), which were also the only species with current mean suitable elevations already exceeding 1000 m (Figure 5). Longitudinal shifts were predominantly westward (35 species, mean: −94.9 km; range: −222.3 to −0.1 km), whereas eastward shifts were less frequent (9 species; mean: +17.1km; range: +1.9 to +37.2 km). Latitudinal shifts were mainly northward (33 species; mean: +20.8 km; range: +2.4 to +58.9 km), whereas southward shifts were less frequent (11 species; mean: −10.2 km; range: −33.2 to 0 km) (Figure 4a).

**Figure 4.**
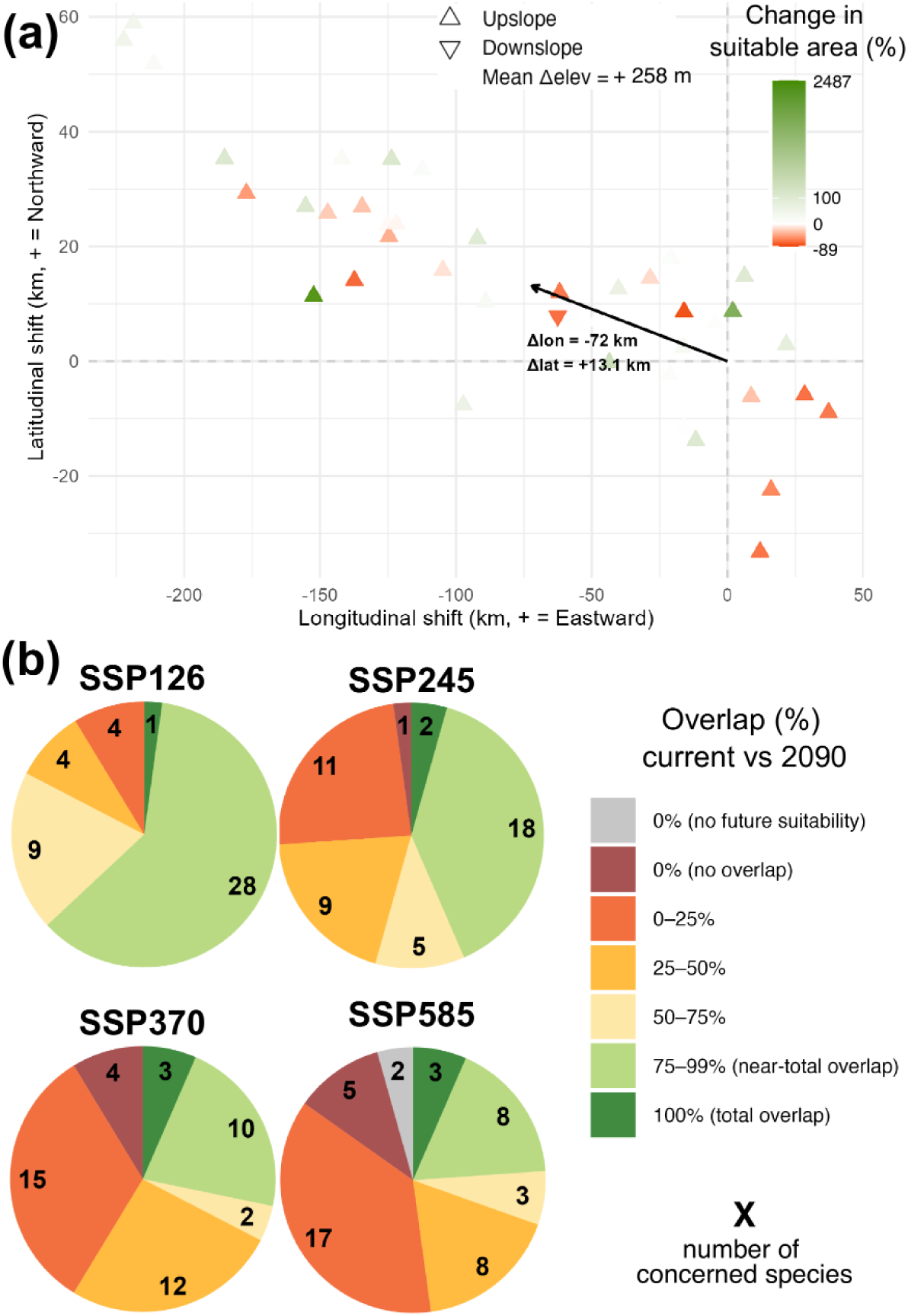
Projected distribution changes in bioclimatic suitability for invasive plant species between the current (1970-2000) and 2081-2100 periods in the Pyrenees. **(a)** Projected latitudinal, longitudinal and elevational shifts of the centroids of bioclimatic suitability for the 2081–2100 period, averaged across SSPs. Each triangle represents one of the 44 species retaining suitable areas under all SSPs. Triangle color reflects changes in suitable areas (green = gain; red = loss). Triangle orientation indicates elevational shifts: upward-pointing triangles denote upslope shifts, and downward-pointing triangles denote downslope shifts. The black arrow indicates the overall centroid mean displacement in latitude (△ lat) and longitude (△ lon) across species. **(b)** Spatial overlap (%) between current and future (2081–2100) bioclimatic suitable areas for invasive plant species under SSPs. Each pie chart displays the number of species considered and their distribution among overlap classes, ranging from total overlap (100%) to no overlap (0%). 2021-2040, 2041-2060, 2061-2080, 2081-2100 periods designated as 2030, 2050, 2070, 2090.

**Figure 5.**
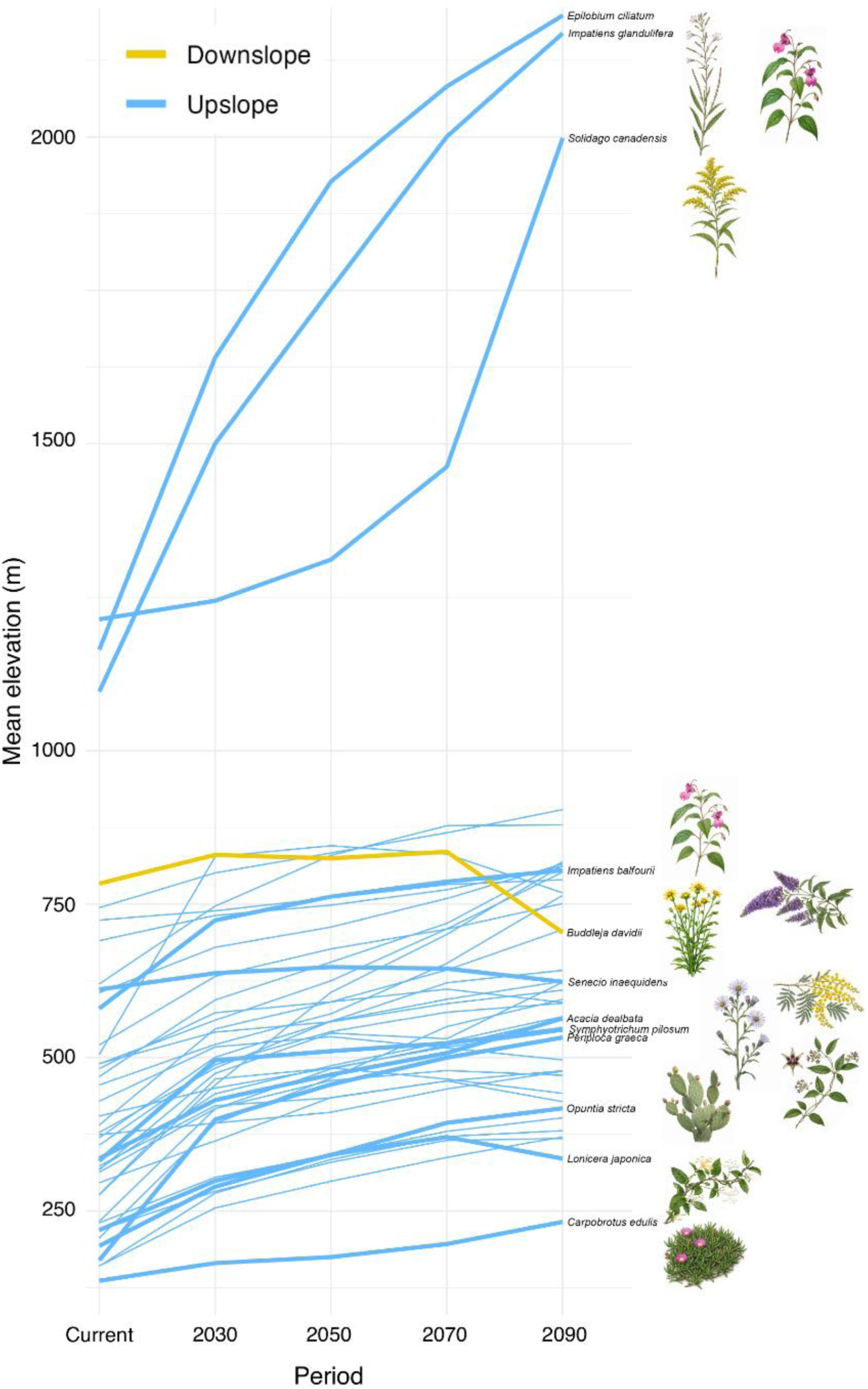
Projected elevational shifts in bioclimatic suitability for invasive plant species between the current (1970-2000) and 2081-2100 periods in the Pyrenees. Projected elevational shifts of the centroids of bioclimatic suitability of invasive plant species, averaged across SSPs. Yellow lines indicate downslope shifts and blue lines upslope shifts relative to the current period. Bolded traits are discussed in more detail in the Discussion section.

Spatial overlap between current and future suitable areas declined steadily as emission levels increased (Figure 4b). Under the most optimistic scenario (SSP126), twenty-nine species maintained high (>75 %) overlap with their current climatic range. This number decreased under intermediate scenarios (e.g., twenty species under SSP245 and thirteen under SSP370). Under the most pessimistic scenario (SSP585), overlap collapsed: seven species showed no overlap with their present climatic niche, seventeen retained 0–25 % overlap, and eleven maintained near (75 %) to complete overlap. Low overlap resulted either from the disappearance of present climatic conditions (SSP585: 2) or a strong spatial displacement of suitable areas relative to the current range (SSP245: 1 species; SSP370: 4; SSP585: 5).

### Rising invasion risk in montane belts with limited overlap with endemic hotspots

Invasive hotspot analysis, defined as areas where multiple species simultaneously retain suitable conditions, revealed marked changes (Figure 6). Maximum hotspot suitability declined from 0.8 under current conditions to 0.72 by the 2081–2100 period, accompanied by an increasingly pronounced upward displacement under more pessimistic climate scenarios. The most persistent hotspot was concentrated in the Atlantic sector of the Pyrenees, consistent with westward shifts in species centroids of bioclimatic suitability. No hotspot extended above 2 000 m in any scenario, while low-elevation plains south of the range became increasingly unsuitable, contributing to the overall decline in hotspot maximum suitability.

**Figure 6.**
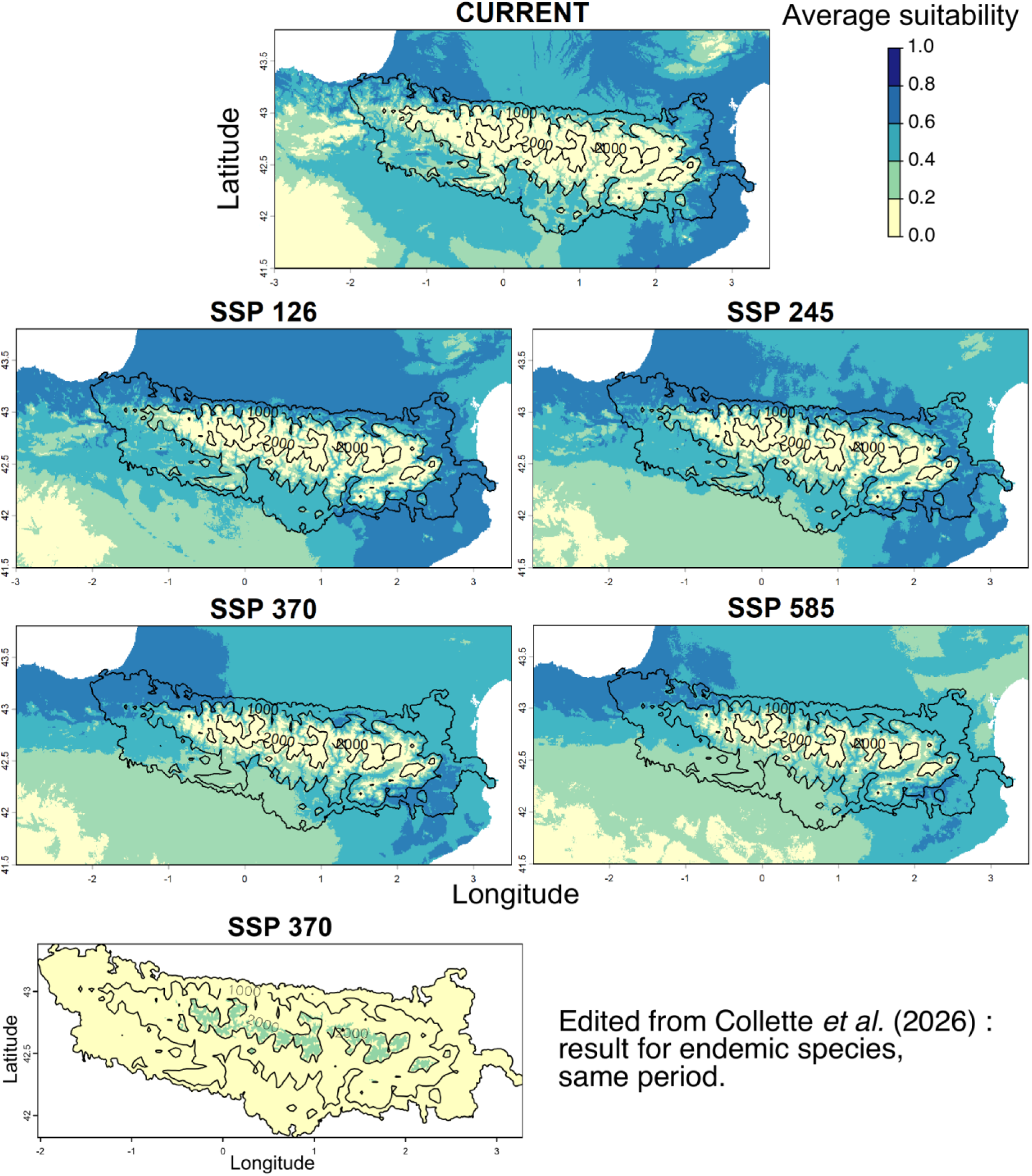
Current and future bioclimatic hotspots for invasive plant species in the Pyrenees. Current and projected (2081-2100) bioclimatic hotspots, i.e. areas where the highest number of species could find suitable bioclimatic conditions, under Shared Socioeconomic Pathways (SSPs). Darker colors indicate higher bioclimatic suitability. Contour lines mark the 1000- and 2000-meter elevation topography. The bottom inset reproduces part of a figure from Collette et al. (2026), presenting results obtained with the same methodology for 59 endemic Pyrenean species over the same period under SSP370, allowing direct comparison with the present study.

Spatial overlap between invasive and endemic bioclimatic hotspots in the Pyrenees was limited under current climatic conditions and declined sharply under future scenarios (Figure 7). Under current conditions, hotspot overlap covered 2 193 km², corresponding to 13.3 % of endemic hotspot area and 8.3 % of invasive hotspot area (Jaccard index = 0.05). By the 2081-2100 period, overlap declined across all scenarios, decreasing to 880 km² under SSP126 (8.4 % of endemic and 2.9 % of invasive hotspots; Jaccard index = 0.02), and becoming nearly negligible under SSP245 (< 3.8 km²; Jaccard index ≈ 0). No spatial overlap was detected under the most pessimistic scenarios (SSP370 and SSP585).

**Figure 7.**
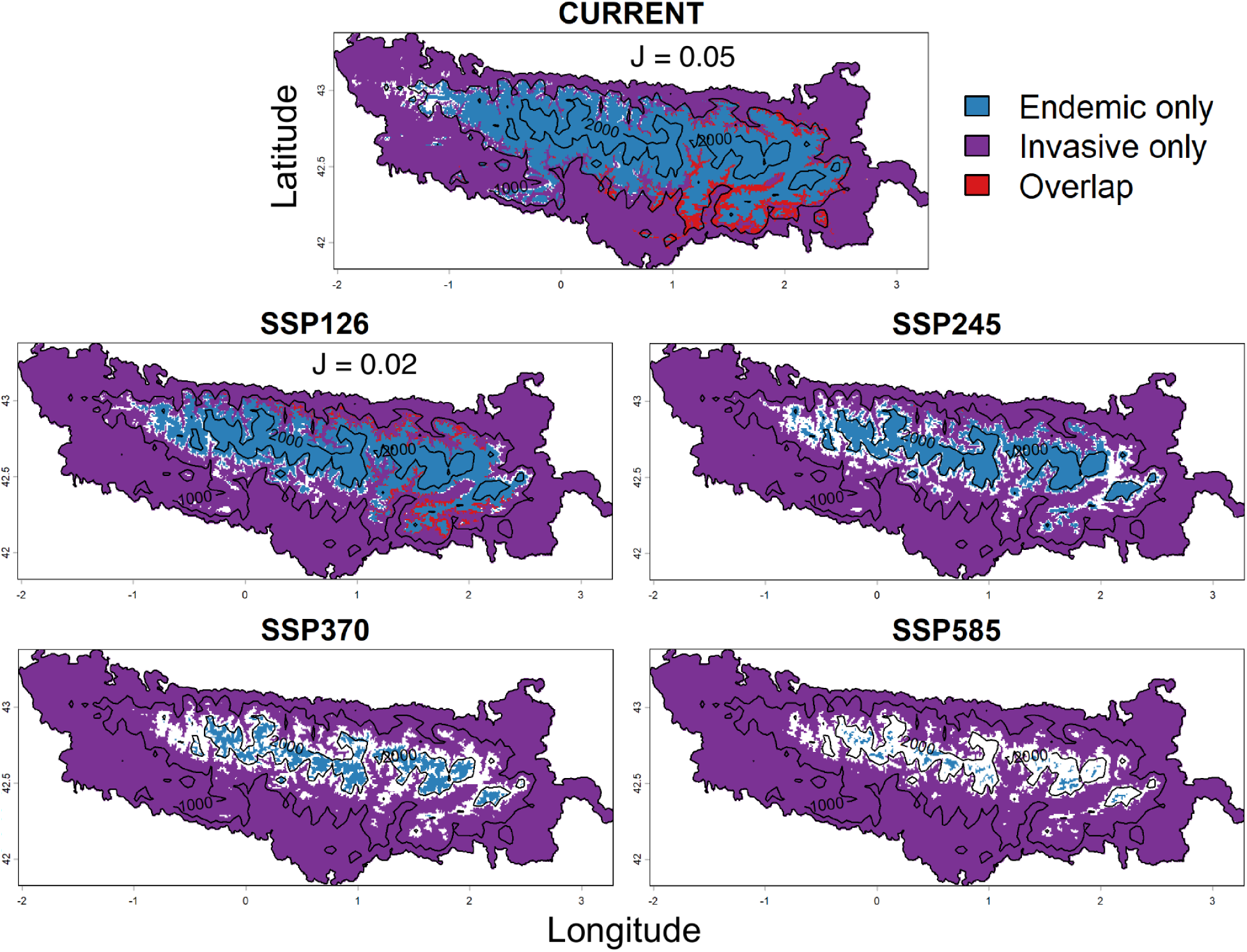
Spatial overlap between invasive and endemic bioclimatic hotspots in the Pyrenees. Current and projected (2081–2100) bioclimatic hotspots for invasive and endemic plant species under SSPs, binarized using a threshold of 0.2 to retain all areas showing any hotspot signal and facilitate the assessment of spatial overlap. Areas exclusive to endemic species (blue), invasive species (purple), and their overlap (red) are shown. Spatial overlap is quantified using the Jaccard index (J; shown when > 0). Black contours denote the 1,000 and 2,000 m elevation isolines.

## Discussion

Our study provides a range-wide, spatially explicit assessment of invasion risks in a major European mountain system and reveals that climate change will not systematically promote invasive plant expansion in mountain landscapes. Although invasive plant species are generally perceived, and often reported, as “winners” under climate change (Goury *et al.*, 2025), we found that 57% are more likely to be “losers”, with twenty species identified as putative winners. These results contrast with those of comparable studies and call for a re-evaluation of the status of invasive species and a better adjustment of control strategies.

### Climate change will promote invasive plant expansion in mountain landscapes mainly under optimistic climate scenarios

Our results challenge the expectation that invasive species generally benefit from global change. Most of the assessed taxa (57% of the 46 species) are projected to lose suitable bioclimatic areas by 2081–2100 period under at least one emission scenario, on average by +18.7 % under SSP126 to −75.1 % under SSP585. Importantly, among the 26 species projected to experience range losses, 14 could still become “winners” under at least one scenario, specifically the most optimistic ones, thereby contributing to the net gains observed under SSP126 for “losers” species. In contrast, all these species consistently remained “losers” under the SSP585 scenario, indicating that apparent climatic “winners” may quickly become “losers” once tolerance thresholds are exceeded. Two species even lose all suitable bioclimatic areas, *Paspalum dilatatum* and *Elaeagnus angustifolia*, under the most pessimistic scenario during 2081–2100 period. This demonstrates that even invasive species, often assumed to tolerate broad environmental conditions (Eyster & Wolkovich, 2021; Godoy *et al.*, 2011; Higgins & Richardson, 2014) and to gain a relative advantage over native species under climate change (Deslippe & Veenendaal, 2025; Liu *et al.*, 2017; Yang *et al.*, 2022), can experience sharp declines as climatic stress intensifies. Although part of the projected range contraction may reflect the decreasing surface available at higher elevations in mountain systems such as the Pyrenees (Elsen & Tingley, 2015), this intrinsic topographic constraint forces species tracking suitable climates upslope into progressively smaller areas, thereby increasing their vulnerability, ecological pressures, and potentially concentrating invasive species within restricted elevational bands. Conversely, a smaller subset of species showed expansion potential, with twenty species increasing their suitable bioclimatic area on average by +388.4% under SSP126 and up to +658.6% under SSP585. Among these, *Symphyotrichum pilosum* and *Periploca graeca* showed particularly pronounced gains, exceeding 2600% and 4100% relative to current conditions, respectively.

Together, these findings reinforce emerging global evidence that invasive species do not uniformly benefit from warming, a conclusion already suggested by early global assessments showing that climate change does not systematically advantage these species (Bellard *et al.*, 2013; Thuiller *et al.*, 2007). Moreover, the literature consistently shows that invasion success is context-specific, shaped by interactions between species characteristics (Buckley & CsergŐ, 2017) and recipient ecosystem properties (Catford *et al.*, 2022; Gioria *et al.*, 2023; Kaushik *et al.*, 2022; Novoa *et al.*, 2020; Pysek *et al.*, 2020a).

### Climate change reshapes invasion risk toward mountain belts

For invasive species, climate change is expected to drive spatial redistribution of suitable bioclimatic conditions, reshaping potential invasion pathways. Spatial overlap between current and future suitable areas declines with increasing emission scenarios, indicating a growing mismatch between present-day distributions and future climatic conditions. This reflects two complementary processes: the loss of currently suitable climates for some taxa, and the spatial displacement of suitability for others. Both point to pronounced distributional turnover and increasing constraints on species’ ability to track suitable conditions under high-emission trajectories (Chan *et al.*, 2024; Vitasse *et al.*, 2021), with implications for invasion risk assessment. Among taxa retaining suitable conditions across all climate scenarios by 2081-2100 period, the centroid of bioclimatic suitable areas is projected to shift upward by an average of +258 m, northward by 13.1 km and westward by 72 km. In the Pyrenees, these align with projected increases in mean annual temperature (+2.2 °C under SSP126 to +6.1 °C under SSP585) and declining precipitation (−3.6 mm under SSP126 to −132.4 mm under SSP585) (Collette *et al.*, 2026), indicating upslope tracking of suitable climates as foothill environments become increasingly restrictive. Westward and northward shifts further reflect climatic contrast across the range, with wetter Atlantic conditions in the western Pyrenees and warmer climates prevailing south of the range in Spain. These patterns are also consistent with the widespread altitudinal and latitudinal shifts in plant distributions documented in the literature over recent decades (Iseli *et al.*, 2023; Lenoir *et al.*, 2008; Rubenstein *et al.*, 2023). The upward recruitment is likely to be further reinforced by anthropogenic dispersal processes. Roads and valley corridors act as efficient conduits for upslope spread (Iseli *et al.*, 2023; Vorstenbosch *et al.*, 2020), as reflected by the clustering of current occurrence records along valleys, influenced by a large lowland urban center (>250 000 inhabitants) and the proximity of the Mediterranean coast. In addition, recreational and agro-pastoral activities can inadvertently transport propagules into otherwise isolated high-elevation habitats and facilitate their establishment by disturbing natural habitats (Barros *et al.*, 2025; Hemp, 2008; Koyama *et al.*, 2024; Liedtke *et al.*, 2020; Montagnani *et al.*, 2022). Together, these mechanisms highlight mountain valleys and adjacent elevational gradients as key areas for early detection, rapid response and coordinated transboundary management.

Despite these upslope shifts, no invasion hotspots, defined as areas where the highest number of species could simultaneously find suitable conditions, are projected to occur above 2 000 m, highlighting the persistent environmental constraints of alpine environments throughout the century. This pattern is consistent with the documented decline in species richness with elevation (Alexander *et al.*, 2016; Di MUSCIANO *et al.*, 2021), with few species occurring in the alpine zone (Becker *et al.*, 2005). However, the persistence and intensification of hotspots within the montane belt indicate increasing invasion pressure at higher elevations. In contrast, hotspots in the southwestern foothills and adjacent lowlands show marked losses of climatic suitability, suggesting a progressive retreat of invasive plants from areas expected to become excessively warm or dry. Lowland areas may become suitable for other non-native species adapted to warmer and drier conditions, potentially replacing those projected to decline. Together, these patterns suggest that the studied invasive species may increasingly concentrate at intermediate elevations, consistent with the recurrent mid-elevation peak in plant species richness along elevational gradients (Mccain & Grytnes, 2010), as limited suitability in lowland and alpine zones constrains further downslope or upslope expansion. In this context, mid-elevation slice of the Pyrenees may function as a climatic refugee but also as a climatic trap, temporarily retaining invasive populations within a narrowing elevational band.

At the regional scale, these projected elevational shifts in species distributions are supported by empirical evidence from a mountain system located south of the Pyrenees, where vegetation belts have already shifted upslope, with Mediterranean formations expanding and montane and subalpine communities regressing, in response to rising temperatures and land-use changes, particularly agro-pastoral abandonment (PEÑUELAS & Boada, 2003). In the Pyrenees themselves, invasion patterns already reflect strong biogeographical and anthropogenic structuring (MartÍnez-Fuentes *et al.*, 2025). On the Mediterranean side, coastal areas host the highest richness of established alien species, some of which have spread into the Pyrenean foothills and subalpine zones. In central areas such as Andorra, most introductions are recent, mainly after the 1980s, coinciding with the rise of mountain tourism and the ornamental plant trade (Soto *et al.*, 2025). In the southern Pyrenees, introduced species originate from diverse biogeographical regions, although many are associated with Mediterranean and temperate climatic conditions rather than boreal–alpine environments (Nualart *et al.*, 2026). Consequently, alien species are mainly concentrated at low to mid-elevations, where such habitats are available (Ninot *et al.*, 2007). In line with these observations, several species in our dataset already persist within the montane and, in some cases, alpine belts, suggesting that future climates could promote the expansion of these early colonizers and facilitate the establishment of additional taxa at higher elevations. Among the 46 studied species, 12 are already officially recognized as invasive in Andorra (Appendix S1-A), indicating that several invasive taxa are already established within the core of the Pyrenean range. However, projections suggest heterogeneous future responses among these species, with only two projected to increase their suitable area across all SSP scenarios, while up to six could be classified as climatic “winners” under the most optimistic scenario. These contrasted trajectories highlight substantial interspecific variation in climatic responses to future warming.

### Spotlights on trajectories of famous climatic “winners”, conditional “winners” and consistent “losers”

Species-specific climatic trajectories reveal heterogeneity in how invasive taxa may respond to future warming. To illustrate these contrasts, we focus on representative species from our dataset. Among the most consistent climatic “winners”, *Opuntia stricta* shows continuous increases in projected suitability across all scenarios. This perennial succulent is tolerant to heat, drought and climatic seasonality, with evidence of persistence beyond arid systems (Gaye *et al.*, 2025; Song *et al.*, 2025). It preferentially establishes in open, disturbed habitats associated with human activities (Misuri *et al.*, 2025; Shackleton *et al.*, 2017). Under future warming, its suitable bioclimatic conditions are projected to expand mainly at mid elevations, where reduced frost frequency may facilitate its upslope expansion into open grasslands and pastoral systems in the Pyrenees. Given its status as one of the world’s most problematic invasive species (Simberloff & Rejmanek, 2019) targeted monitoring is needed where future climatic suitability overlaps with human activities. Similar patterns in other mountain systems suggest that climatic winners under warming often combine thermophilic and ruderal strategies with high colonization capacity, traits overrepresented among invasive taxa (Goury *et al.*, 2025). *Opuntia stricta* exemplifies this syndrome and calls for heightened vigilance in the Pyrenees. Similar trajectories were projected for *Periploca graeca* and *Symphyotrichum pilosum*, two comparatively less documented neophytes associated with warm and disturbed habitats (TUNÇKOL *et al.*, 2017), suggesting that ongoing warming may facilitate the spread of localized thermophilic and ruderal species within mountain regions.

In contrast, *Senecio inaequidens,* an invasive grassland species, and *Lonicera japonica*, a disturbance-associated invasive climber, emerged as conditional “winners”. Both displayed increases in suitability under moderate warming (SSP126–245) followed by declines under high-emission scenarios (SSP370–585) after 2041–2060. Although *S. inaequidens* already recruits at high elevations in the Pyrenees and other mountain systems and has not fully occupied its current climatic niche (Vacchiano *et al.*, 2013), projected declines under severe warming suggest emerging climatic constraints. Its capacity for rapid local adaptation (Monty & Mahy, 2009), tolerance for cold winters (Lachmuth *et al.*, 2010), and early establishment in species-poor communities (Delory *et al.*, 2019) may nonetheless allow persistence despite declining suitability. *Lonicera japonica* displayed a comparable non-linear response, with gains under intermediate scenarios likely reflecting its broad ecological tolerance and affinity for disturbed habitats. Previous studies further suggest that reduced winter frost may facilitate its expansion toward higher elevations (Lemke *et al.*, 2011), although its preference for mesic environments could limit persistence under increasingly dry conditions (Schierenbeck, 2010). Together, these cases illustrate that early climatic gains do not necessarily translate into sustained expansion once key physiological thresholds are exceeded.

A third set of species, including *Buddleja davidii*, a fast-growing invasive shrub disrupting native pollination networks, and *Acacia dealbata*, a thermophilic invasive tree widespread in Mediterranean environments, consistently emerges as climatic “losers”, with projected contractions of suitable areas under all scenarios. In both cases, our hypothesis is that range losses are primarily driven by upward shifts of lower elevational limits rather than successful colonization at higher elevations, resulting in net range contraction (Zu *et al.*, 2021). This pattern is consistent with evidence from the Alps, where successful upslope colonization by invasive plants is largely restricted to cold-tolerant taxa (Petitpierre *et al.*, 2016), and with global analysis showing that local extinctions disproportionately affect populations at warm range margins as annual maximum temperatures increase (RomÁn-Palacios & Wiens, 2020). Despite the broad ecological and altitudinal tolerance of B. davidii in its native range, where it occurs up to 3500 m across temperate to subtropical climates (Tallent & Watt, 2009)), projected declines in the Pyrenees are best interpreted as an altitudinal squeeze. Increasing summer heat, moisture limitation and water stress reduce suitability at lower elevations, while upward expansion remains constrained by persistent cold conditions, temperature variability and minimum growing-season temperatures (Wittig, 2012). Reliance on phenotypic and ecological plasticity rather than strong local adaptation (EBELING *et al.*, 2011) may further limit the ability of this species to overcome upper-elevation climatic constraints. A comparable mechanism may explain the projected decline of *Acacia dealbata*, whose broad climatic tolerance across Mediterranean and subtropical regions contrasts with its sensitivity to prolonged frost, cool mountain conditions and reduced water availability, potentially limiting successful establishment at higher elevations while increasing drought stress contracts suitability at lower elevations (DessÌ *et al.*, 2021; Vieites-Blanco & GonzÁlez-prieto, 2020).

Together, these examples illustrate a continuum of invasion trajectories shaped by species-specific combinations of climatic tolerance, habitat dependence and sensitivity to climatic thresholds. While identifying traits associated with persistent expansion can inform risk assessment (but see EARLE *et al.*, 2023; MOLES *et al.*, 2012), invasion outcomes remain strongly context dependent. Our exploratory trait analyses further support this context dependency, as no invasion-history or ecological attribute significantly differentiated climatic winners from losers.

### Climate could constrain direct competition between invasive and endemic plants in the Pyrenees

A comparable analysis on Pyrenean endemic plant species (Collette *et al.,* 2026) indicates that suitability hotspots for endemics are projected to remain largely confined above 2 000 m throughout the century. In contrast, our results show that hotspots of invasive species occur below this elevation, despite a general upward shift under future warming scenarios. This consistent elevational segregation suggests that climate change may act as an environmental filter, allowing endemic taxa to persist in high-elevation climatic refugia while limiting the upward expansion of most invasive species.

Consistent with this pattern, our results indicate that climatic constraints may restrict the long-term co-occurrence of invasive and endemic plants. The spatial overlap between invasive and endemic plant hotspots is projected to strongly contract, decreasing from 13.3 % under current conditions to nearly zero in the future. The decline in invasive hotspot intensity, combined with their upward displacement potentially constrained below 2,000 m, suggests that ongoing warming will substantially reduce the area simultaneously suitable for both invasive and endemic taxa, thereby limiting the spatial scope for direct biotic interactions. Even where invasive and endemic species co-occur at high elevations, the impacts of non-native plants on alpine ecosystems are not systematic and have so far been reported as limited (Alexander *et al.*, 2016). Under our scenarios, invasive and endemic plants are projected to increasingly occupy distinct climatic niches, reducing the likelihood of widespread direct competition along the mountain chain, even if endemic species face substantial constraints in tracking suitable conditions. Nevertheless, such upward invasion pressure still raises management concerns, as alpine and subalpine communities could remain vulnerable to replacement by taxa that track warming climates more efficiently (Bradley *et al.*, 2024; Dainese *et al.*, 2017; Geppert *et al.*, 2023).

### Rethinking invasive risks under climate change: should invaders be “allowed” to persist for the sake of species conservation?

The divergent trajectories observed among taxa call for a reassessment of how invasion risk is defined in the context of rapid global change. Our results reveal contrasting situations in which the same mountain system may either facilitate or constrain the persistence of non-native species. *Carpobrotus edulis* illustrates this complexity in our analysis. In the Pyrenees, this species is a climatic winner, projected to expand its suitable area under future warming. Yet, model-based extinction-risk assessments suggest that it may become threatened in parts of its native range (Bachman *et al.*, 2024), despite its widespread distribution. Here, mountain systems may act as climatic refugia, providing a non-native taxa one of the last environments matching its fundamental climatic niche. Conversely, climatic losers such as *Acacia dealbata* illustrate the opposite trajectory. This species is currently considered an important invasive tree in southern Europe, particularly in disturbed and fire-prone environments where it can form dense stands and threaten native vegetation (Lorenzo *et al.*, 2010). Our projections suggest that its long-term persistence in the Pyrenees may become increasingly limited under future climatic conditions, potentially reducing its ecological impacts on the region. Here, mountain systems may instead act as climatic barriers, or even as traps, limiting future establishment and spread. Together, these contrasting outcomes raise a fundamental question: should global invasive species be managed uniformly when their climatic vulnerability and persistence differ across regions (Baquero *et al.*, 2023)? Beyond this dichotomy, our framework also allows for a third outcome, in which species may face extinction risks in both their native and invaded regions, as illustrated by *Impatiens balfourii* in our study. Such cases exemplify the conservation–invasion paradox, whereby invasive species may simultaneously pose ecological threats in some regions while raising conservation concerns in others (Hong *et al.*, 2025; Staude *et al.*, 2025), calling for more nuanced, context-dependent management strategies. In mountainous systems where most endemic species are threatened by climate change regardless of invasive species (Collette *et al.*, 2026), the future composition of the flora becomes a central issue. However the role of non-native and invasive species in this composition is unclear. In the Pyrenees, our results indicate only marginal spatial overlap between endemic and invasive plants, suggesting that invasive species may contribute to future floristic reorganization without constituting a primary threat to endemic persistence through direct spatial co-occurrence. Beyond species redistribution, further investigation is required into whether invasive species may also shape future flora through the generation of evolutionary novelty. Such novelty may arise through rapid adaptation (Clements & Jones, 2021; Moran & Alexander, 2014) or through hybridization with native taxa or among non-native lineages, potentially leading to the emergence of new evolutionary lineages (Ellstrand, 2009; Ellstrand & Schierenbeck, 2000) and thus generate novel evolutionary outcomes (Meyerson *et al.*, 2010; Mooney & Cleland, 2001) and diversity when colonizing new environments (Vellend *et al.*, 2007). However, although evolutionary change can occur over short timescales, its magnitude is frequently insufficient to keep pace with the rate of projected climate change (Jezkova & Wiens, 2016).

### Using bioclimatic niche projections to inform proactive invasion management

Although invasive species are increasingly recognized as an emerging threat in mountain systems, the Pyrenees still lack a harmonized transnational framework to identify priority areas for surveillance and to coordinate management across administrative boundaries. This gap is particularly critical in a transboundary mountain range where invasion pathways and suitable conditions extend beyond political borders. In this context, focusing monitoring efforts on areas projected to become climatically suitable could improve management efficiency, as early detection and rapid response are consistently more cost-effective and ecologically sustainable than reactive control following establishment (Ahmed *et al.*, 2022; Hanley & Roberts, 2019; Venette *et al.*, 2021). This is particularly relevant in mountain landscapes, where rugged and inaccessible terrain constrains mechanized control and increases reliance on labor-intensive interventions, thereby amplifying both financial and human costs (Mcdougall *et al.*, 2011). However, bioclimatic projections should not be interpreted as direct prescriptions for disengagement. While species projected to lose climatic suitability may justify a gradual reallocation of management effort, complete disengagement is rarely warranted. Taxa may persist long after climatic conditions become unsuitable, a process described as extinction debt (Kuussaari *et al.*, 2009). Even after population decline or local eradication, invasive species may continue to exert ecological and economic impacts through persistent legacies on soils, biotic interactions and ecosystem processes, affecting vulnerable communities well beyond their expansion phase (Ahmad *et al.*, 2021; Corbin & D’Antonio, 2012; Essl *et al.*, 2015; Prior *et al.*, 2018).

More broadly, our findings highlight a growing mismatch between climatic opportunities and conventional invasion risk assessments, calling for a reassessment of how non-native species are evaluated and prioritized under ongoing climate change. Climate change increasingly blurs the boundary between natural range shifts and biological invasions, challenging traditional invasion categories and complicating the terminology and frameworks traditionally used in invasion ecology (Fusco *et al.*, 2024; Walther *et al.*, 2009). These dynamics call for more flexible, context-dependent management frameworks, often referred to as climate-smart approaches (Colberg *et al.*, 2024), which highlight the importance of considering climate-driven shifts in species distributions and persistence, as well as their implications for management feasibility and effectiveness. Our results indicate that mountain systems such as the Pyrenees may simultaneously act as climatic barriers, refugia, or potential traps for non-native taxa. A central challenge lies in distinguishing species likely to lose climatic suitability and ecological relevance from those that may benefit from warming, in assessing the magnitude of their future impacts, and in anticipating which taxa could increasingly contribute to the future emblematic flora of the Pyrenees as suitable conditions disappear elsewhere. Integrating genetic, ecological and environmental data across invasion stages provides a robust framework for anticipating invasion dynamics under global change, and supports the development of flexible, climate-informed management strategies that explicitly account for evolutionary and demographic processes (SHERPA & DesprÉs, 2021).

## Data availability statement

The original data supporting the findings of this study are openly available from public domain resources, including GBIF, WorldClim. The code and data required to reproduce the results are accessible in the Zenodo repository at [DOI: 10.5281/zenodo.18114751]. Further supporting details are available in the Supplementary Information S1 and S2.

