## Supplemental data for "Climbing invasions or climatic refugees: how many and to which extent non-native plant species could reach the top of the Pyrenees mountains under climate change?"

**Table of Contents:**


| | |
|---|---|
| **Appendix A: Sources used to define the regional invasive species pool surrounding the Pyrenees** | Page 2 |
| **Appendix B: Detailed overview of SDMs framework according to the ODMAP protocol.** | Page 3 |
| **Appendix C: Current and future continuous bioclimatic niche suitability maps for invasive plants in the Pyrenees.** | Page 11 |
| **Appendix D: Longitudinal, latitudinal and elevational shifts in climatic suitability for invasive plant species in the Pyrenees under future climate scenarios for 2081-2100 period.** | Page 58 |
| **Appendix E: Species richness and density of invasive plant occurrence records across elevational belts in the Pyrenees.** | Page 63 |
| **Appendix F: Model performance evaluation** | Page 64 |
| **References for the Appendix** | Page 65 |

**Appendix A: Sources used to define the regional invasive species pool surrounding the Pyrenees**

This appendix documents the external reference lists used to identify non-native plant species recognized as invasive in regions adjacent to the Pyrenees. Species selection was based on official national and regional inventories from areas directly surrounding the Pyrenean massif, ensuring that the regional invasive species pool reflects invasion contexts relevant to the study area. The resulting species selection is documented at DOI: 10.5281/zenodo.18114751 (Appendix S2)

**Regions considered**

**France**

**Occitanie**: COTTAZ C., DAO J. & HAMON M., 2021. Liste de référence des plantes exotiques envahissantes de la région Occitanie. Synthèse, analyses de risque et catégorisation des taxons. Document technique des CBN d’Occitanie (CBNMed et CBNPMP). 46 p. + annexes

**Nouvelle-Aquitaine**: CAILLON A. (coord.), BONIFAIT S., CHABROL L., DAO J., LEBLOND N., RAGACHE Q., 2022 – Liste hiérarchisée des plantes exotiques envahissantes de Nouvelle-Aquitaine. – Conservatoire Botanique National Sud-Atlantique (coord.), Conservatoire Botanique National du Massif central et Conservatoire Botanique National des Pyrénées et de Midi-Pyrénées. 116 pages + annexes.

**Spain**

**Catalonia**: AYMERICH P. & SÁEZ L. 2019. Checklist of the vascular alien flora of Catalonia (northeastern Iberian Peninsula, Spain). Mediterranean Botany . 40(2): 215-242. DOI 10.5209/mbot.63608.

**Aragon**: *Gobierno de Aragón. Fichas de especies problemáticas para Aragón.* https://www.invasara.es/fichas-especies-problematicas-para-aragon/ (accessed 2025-11).

**Navarra**: GOBIERNO DE NAVARRA. 2015. ESPECIES DEL CATÁLOGO ESPAÑOL DE EXÓTICAS INVASORAS DETECTADAS EN NAVARRA.

**Andorra :** https://www.bopa.ad/bopa/034075/Pagines/GR20220617_11_55_53.aspx

**Appendix B: Detailed overview of SDMs framework according to the ODMAP protocol**

Overview, Data, Model, Assessment, and Prediction (ODMAP) protocol (ZURELL *ET AL.*, 2020) provides a standardized framework for transparently and reproducibly reporting species distribution models.

**Table B1: Overview, Data, Model, Assessment, and Prediction (ODMAP) protocol**. The following table details our study methodology according to the ODMAP structure. **Data and code availability links are provided in the Overview element, under the *Software, Codes, and Data* section**.

| ODMAP element | Contents |
|---|---|
| **OVERVIEW** | |
| *Authorship* | · **Authors:** |
| | · **Contact email:** |
| | · **Title:** Climbing invasions or climatic refugees: how many and to which extent non-native plant species could reach the Pyrenees mountains under climate change? |
| | · **DOI:** |
| *Model objective* | · **SDM purpose:** forecast / transfer |
| | · **Main target output:** continuous and binary bioclimatic suitability maps |
| *Taxon* | Plant species recognized as invasive on both sides of the Pyrenees |
| *Location* | The Pyrenees massif (Körner *et al.*, 2011, Snethlage *et al.*, 2022), southwestern Europe |
| *Scale of analysis* | · **Spatial Extent (Lon / Lat):** (xmin, xmax, ymin, ymax)<br><br>europe_extent (for calibration): -11, 19.5, 35.7, 63.5<br>pyrenees_extent : -3, 3.5, 41.5, 43.8 |
| | · **Spatial resolution:** 1kmx1km |
| | · **Temporal extent/time period:** 1970-2100 |
| | · **Temporal resolution:** 5 periods 1970-2000, 2021-2040, 2041-2060, 2061-2080, and 2081-2100 (designated as current, 2030, 2050, 2070, 2090 respectively) |
| | · **Type of extent boundary:** rectangular for modeling, and natural boundary of the Pyrenees for analyses |
| *Biodiversity data overview* | · **Observation type:** citizen science, field survey |
| | · **Response/data type:** presence-only |
| *Type of predictors* | Climatic |
| *Conceptual model* | · **Hypotheses about species-environment relationship:** species distributions are mainly driven by temperature and precipitation. Biotic interactions have a negligible impact. All populations of a given species respond uniformly to climatic variables over time, without accounting for local adaptations or genetic variability. |

| | |
|---|---|
| *Assumptions* | (1) Species are in equilibrium with the environment; (2) Biases in the modeling system are minimal; (3) Species niches are conserved over time; (4) All model variables are related to species occurrence. |
| *SDM algorithms* | · **Model algorithms:** Maximum entropy (Maxent), Generalized Linear Model (GLM), Gradient Boosting Model (GBM), Generalized Additive Model (GAM), Random Forest (RF) |
| | · **Model complexity:** we let the data determine model complexity. Model settings were automatically optimized to achieve the best performance. Tested settings were chosen to generate an intermediately complex response while preventing excessive overfitting yet still covering a sufficiently broad range. |
| | · **Model averaging:** Median of models over a maximum of 10 repetitions of each model algorithm (maximum 50 models). |
| *Model workflow* | **(1)** Species occurrence data downloading & cleaning.<br>**(2)** Processing climate data.<br>**(3)** Sampling of pseudo-absence points.<br>**(4)** Individual model fitting.<br>**(5)** Ensemble modeling generation.<br>**(6)** Analysis of bioclimatic niche suitability dynamics. |
| *Software, codes and data* | Computational environment:<br>Processor: Apple M2 Max<br>Total number of cores: 12<br>Total RAM memory: 64 GB<br><br>Analyses were conducted with R version 4.4.1 (R Core Team, 2024) in Rstudio environment (Posit team, 2024).<br><br>Key packages:<br>Retrieval occurrence data from GBIF – packages 'geodata' (Hijmans *et al.*, 2024) and 'rgbif' (Chamberlain *et al.*, 2025 ; Chamberlain & Boettiger, 2017).<br>Data cleaning – packages 'fuzzySim' (Barbosa, 2015) and 'sf' (Pebesma, 2018 ; Pebesma & Bivand, 2023)<br>Spatial thinning of occurrences, raster manipulation, and model projections – package 'terra' (Hijmans, 2024).<br>Generation of random pseudo-absences – package 'flexsdm' (Velazco *et al.*, 2022).<br>MaxEnt – package 'maxnet' (Phillips, 2021).<br>Generalized Additive Models (GAM) – package 'mgcv' (Wood, 2017)<br>Generalized Boosted Regression Models (GBM) – package 'gbm' (Ridgeway & Developers, 2024)<br>Random Forest – package 'randomForest' (Liaw & Wiener, 2001).<br>Block cross-validation – package 'blockCV' (Valavi *et al.*, 2019).<br>Computation of sensitivity, Boyce Index, AUC, and MaxSSS threshold – package 'modEvA' (Barbosa *et al.*, 2013).<br>Parallelization using mclapply – package 'parallel' (R Core Team, 2024) *(note: this approach does not work on Windows).* |
| | · **Data availability:** code available at [DOI: 10.5281/zenodo.18114751] |
| | · **Data availability:** GBIF Occurrences data: [DOI: 10.5281/zenodo.18114751] Environmental data: https://www.worldclim.org/ |

| DATA | |
|---|---|
| *Biodiversity data* | · **Taxon names:** *Acacia dealbata, Acer negundo, Agave americana, Ailanthus altissima, Araujia sericifera, Artemisia verlotiorum, Azolla filiculoides, Bidens frondosa, Bidens subalternans, Buddleja davidii, Carpobrotus acinaciformis, Carpobrotus edulis, Cortaderia selloana, Cyperus eragrostis, Elaeagnus angustifolia, Epilobium ciliatum, Erigeron canadensis, Erigeron karvinskianus, Erigeron sumatrensis, Euphorbia prostrata, Fallopia baldschuanica, Gleditsia triacanthos, Helianthus tuberosus, Impatiens balfourii, Impatiens glandulifera, Ligustrum lucidum, Lonicera japonica, Ludwigia peploides, Opuntia stricta, Parthenocissus inserta, Paspalum dilatatum, Periploca graeca, Phyllostachys aurea, Phyllostachys bambusoides, Phyllostachys nigra, Phytolacca americana, Prunus cerasifera, Robinia pseudoacacia, Senecio inaequidens, Sicyos angulatus, Solidago canadensis, Sorghum halepense, Sporobolus indicus, Symphyotrichum pilosum, Symphyotrichum squamatum, Xanthium orientale* |
| | · **Ecological level:** species |
| | · **Data source: GBIF:** [DOI: 10.5281/zenodo.18114751] |
| | · **Sampling design:** opportunistic & standardized monitoring sampling |
| | · **Sample size per taxon:** |

| **Species** | **Raw (study area, 1970+, uncertainty ≤ 1 km)** | **Gridded (after spatial thinning)** |
|---|---|---|
| *Acacia dealbata* | 52054 | 32588 |
| *Acer negundo* | 94680 | 56288 |
| *Agave americana* | 19868 | 12762 |
| *Ailanthus altissima* | 91630 | 41387 |
| *Araujia sericifera* | 22569 | 11628 |
| *Artemisia verlotiorum* | 15149 | 9588 |
| *Azolla filiculoides* | 13814 | 8343 |
| *Bidens frondosa* | 32284 | 20255 |
| *Bidens subalternans* | 4548 | 3655 |
| *Buddleja davidii* | 121516 | 44718 |
| *Carpobrotus acinaciformis* | 2798 | 1489 |
| *Carpobrotus edulis* | 23975 | 7868 |
| *Cortaderia selloana* | 26793 | 15259 |
| *Cyperus eragrostis* | 36844 | 25153 |
| *Elaeagnus angustifolia* | 36599 | 17461 |
| *Epilobium ciliatum* | 75721 | 55554 |
| *Erigeron canadensis* | 126382 | 72988 |
| *Erigeron karvinskianus* | 36807 | 19728 |
| *Erigeron sumatrensis* | 43285 | 24264 |
| *Euphorbia prostrata* | 16317 | 12112 |
| *Fallopia baldschuanica* | 5501 | 4031 |
| *Gleditsia triacanthos* | 25234 | 15433 |
| *Helianthus tuberosus* | 25119 | 20326 |
| *Impatiens balfourii* | 7251 | 4732 |
| *Impatiens glandulifera* | 329446 | 113054 |
| *Ligustrum lucidum* | 25459 | 14141 |
| *Lonicera japonica* | 58503 | 35460 |

| | | | |
|---|---|---|---|
| | *Ludwigia peploides* | 13143 | 8043 |
| | *Opuntia stricta* | 11902 | 7733 |
| | *Parthenocissus inserta* | 30311 | 21726 |
| | *Paspalum dilatatum* | 39009 | 26875 |
| | *Periploca graeca* | 1677 | 966 |
| | *Phyllostachys aurea* | 2438 | 1961 |
| | *Phyllostachys bambusoides* | 949 | 702 |
| | *Phyllostachys nigra* | 918 | 704 |
| | *Phytolacca americana* | 90921 | 55608 |
| | *Prunus cerasifera* | 65497 | 47998 |
| | *Robinia pseudoacacia* | 111481 | 59863 |
| | *Senecio inaequidens* | 242677 | 36499 |
| | *Sicyos angulatus* | 7485 | 5550 |
| | *Solidago canadensis* | 128689 | 49754 |
| | *Sorghum halepense* | 37415 | 26896 |
| | *Sporobolus indicus* | 14570 | 10052 |
| | *Symphyotrichum pilosum* | 8978 | 6902 |
| | *Symphyotrichum squamatum* | 9717 | 7356 |
| | *Xanthium orientale* | 22974 | 17030 |
| | · **Mask:** We clipped all data to europe_extent (for calibration): -11, 19.5, 35.7, 63.5 | | |
| | · **Details on scaling:** Spatial thinning, retaining one occurrence per grid cell. | | |
| | · **Details on data cleaning/filtering steps:** Occurrences since 1970 within the study area and ≤ 1 km coordinate uncertainty were retained, fitting the resolution of environmental variables. Species lacking GBIF occurrence records in regions adjacent to the Pyrenees were excluded, as species with insufficient occurrences to ensure representation across all spatial blocks required by the spatial cross-validation scheme.<br>Species eliminated are documented in "invasive species list.xlsx" sheet [DOI: 10.5281/zenodo.18114751] | | |
| | · **Details on background data derivation:** We adopted random (and not environmentally constrained due to computational time) pseudo-absences in equal number to presences. We generated 10 independent replicates (10 pseudo-absence sets) per species (Biomod2 team recommendation). | | |
| | · **Details on potential errors and biases in data:** The biases encountered are those related to citizen science (collected using different sampling efforts and methodologies, misidentification potential, collected in more accessible areas...). | | |
| *Data partitioning* | · **Selection of training data:** 5 block cross-validation, with blocks assigned to the training set in each iteration | | |
| | · **Selection of validation data:** Validation data were drawn from spatially independent blocks (determining through blockCV::cv_spatial_autocor function and divided by 1.5 to balance data sufficiency, resulting in a block size of [1353786/1.5] m) excluded from the training set during each cross-validation iteration. | | |
| *Predictor variables* | · **State predictor variables used:** list of 19 bioclimatic variables (11 related to temperature and 8 to precipitation) available on WorldClim 2.1 (Fick & Hijmans, 2017) | | |

| | |
|---|---|
| | · **Details on data sources:** WorldClim 2.1 https://www.worldclim.org/ [accessed 16 January 2025] |
| | · **Spatial resolution and spatial extent of raw data:** 1 km x 1 km, world |
| | · **Map projection (coordinate reference system):** EPSG:4326 |
| | · **Temporal resolution and temporal extent of raw data:** 1970-2000 |
| | · **Details on data processing and on spatial, temporal and thematic scaling:** We clipped current data (1970-2000) to europe_extent (for calibration), and future data (2021-2100) to the pyrenees_extent |
| | · **Details on measurement errors and bias, when known:** See Fick & Hijmans, 2017. Potential biases may arise from interpolation errors, particularly in our study area with sparse meteorological stations or complex topography. |
| *Transfer data for projection* | · **Details on data sources, spatial extent and resolution:** same as *Predictor variables* |
| | · **Temporal extent/time period:** 2021-2100 |
| | · **Temporal resolution:** 4 periods 2021-2040, 2041-2060, 2061-2080, and 2081-2100 (designated as 2030, 2050, 2070, 2090 respectively) |
| | · **Models and scenarios used:** All General Circulation Models available on WorldClim 2.1 for Shared Socioeconomic Pathways (SSPs) SSP126, SSP245, SSP370 & SSP585: ACCESS-CM2, CMCC-ESM2, EC-Earth3-Veg, UKESM1-0-LL, GISS-E2-1-G, INM-CM5-0, IPSL-CM6A-LR, MIROC6, MPI-ESM1-2-HR, MRI-ESM2-0, BCC-CSM2-MR. |
| | · **Details on data processing and scaling:** We clipped all data to the pyrenees_extent. The General Circulation Models were aggregated by taking the median for each period and SSP. |
| | · **Quantification of novel environmental conditions and novel environmental combinations:** NA |
| MODEL | |
| *Variable pre-selection* | The initial variables were selected based on their availability for future periods. |
| *Multicollinearity* | To reduce redundancy in models, correlated variables (Pearson $r > 0.8$) at presence locations for each species were grouped into clusters. Negatively correlated variables can capture complementary environmental gradients (e.g. warm summers vs. cold winters) and were therefore kept in the analysis. From each cluster, the highest-ranked variable was retained based on a predefined ecological priority list : BIO1 & BIO12 > BIO11 > BIO6 > BIO17 > BIO14 > BIO10 > BIO5 > BIO4 > BIO15 > BIO16 > BIO13 > BIO7 > BIO18 > BIO9 > BIO8 > BIO19 > BIO3 > BIO2 |
| *Model settings* | · **Models settings:**<br>For model tuning:<br>GBM:<br>n.trees_vals <- c(100, 200, 500)<br>interaction.depth_vals <- c(1, 2, 3)<br>shrinkage_vals <- c(0.01, 0.1, 0.05) |

| | |
|---|---|
| | n.minobsinnode_vals <- c(5, 10, 15)<br>bag.fraction_vals = c(0.4, 0.5, 0.6)<br>MXT:<br>regmult_vals <- c(0.5, 1, 2, 3)<br>classes_vals <- c("l", "q", "lq", "lqh", "lqhp", "lqhpt")<br>RF:<br>ntree_vals <- c(500, 1000)<br>mtry_vals <- c(floor(sqrt(length(vars_sel))), floor(length(vars_sel) / 2), length(vars_sel))<br>nodesize_vals <- c(5, 10, 20)<br>GAM:<br>sp_vals <- c(0.1, 0.5, 1, 2, 5)<br>GLM:<br>glm_form <- as.formula(paste("presence ~", paste("poly(", vars_sel, ", 2)", collapse = " + "), "+", paste(vars_sel, collapse = ":")))<br>glm_mod <- glm(formula = glm_form, family = binomial(link = "logit"), data = dat_random)<br>glm_mod_opt <- stepAIC(glm_mod, direction = "both")<br><br>Default setting for the other parameters.<br><br>For modeling:<br>GBM:<br>gbm_mod <- gbm::gbm(formula = gbm_form, data = dat_env_const, shrinkage = best_shrinkage, interaction.depth = best_interaction.depth, n.trees = best_n.trees, n.minobsinnode = best_n.minobsinnode, bag.fraction = best_bag.fraction, distribution = "bernoulli")<br>MXT:<br>mxt_mod <- maxnet::maxnet(p = dat_random$presence, data = dat_random[, vars_sel], regmult = best_regmult, maxnet.formula(p = dat_random$presence, data = dat_random[, vars_sel], classes = best_classes))<br>RF:<br>rf_form <- reformulate(termlabels = vars_sel, response = "presence_fact")<br>rf_mod <- randomForest::randomForest(formula = rf_form, data = dat_env_const, na.action = na.exclude, keep.forest = TRUE, importance = TRUE, nodesize=best_nodesize, mtry=best_mtry, ntree = best_ntree)<br>GAM:<br>gam_form <- reformulate(termlabels = supply(vars_sel, function(v) paste0("s(", v, ", sp = ", best_sp, ")")), response = "presence")<br>gam_mod <- mgcv::gam(formula = gam_form, family = binomial, data = dat_random, weights = weights_GAM, method = "REML")<br>GLM:<br>glm_mod <- glm(formula = glm_form, family = binomial(link = "logit"), data = dat_random, weights = weights_GLM)<br><br>GLM, GAM, GBM and RF prevalence (presence/absence ratio) was removed to convert presence/absence model into a favorability model with fuzzySim::Fav function. |
| *Model estimates* | · **Assessment of model coefficients:** Not analyzed. |
| | · **Assessment of variable importance:** Variable importance was assessed by quantifying how much deviance each variable explains. Two summary tables of variable importance are generated upon code execution, one presenting the mean scores per predictor and algorithm across all runs, and another providing details for each individual run. |

| | |
|---|---|
| *Model selection / Model averaging / Ensembles* | · **Model selection strategy:** Models were selected based on cross-validation performance using three metrics: Boyce Index, Sensitivity, and AUC ROC. For each algorithm and iteration, the set of parameters yielding the highest performance score (calculated as a weighted average of 0.5 Boyce Index, 0.25 AUC, and 0.25 Sensitivity) was retained. Only models with a Boyce Index > 0.5 were included in the final ensemble model to ensure robustness and reliability. |
| | · **Model averaging method / ensemble method:** Ensemble modeling was applied at two distinct stages: (i) for current conditions, based on model runs calibrated and validated (Boyce Index > 0.5, maximum 5 algorithms x 10 pseudo absence set = 50 models) under present-day climate; and (ii) for future projections, by projecting these same validated models to every combination of future periods and scenarios, and then creating an ensemble for each period–scenario combination. Consensus ensemble maps were computed as pixel-wise median across all algorithms and replicates. |
| *Non-independence correction/analyses* | · **Method for addressing spatial autocorrelation in residuals:** Spatial autocorrelation was addressed by analyzing a variogram of the environmental predictors to determine an appropriate block size for spatial cross-validation (determining through blockCV::cv_spatial_autocor function and divided by 1.5 to balance data sufficiency, resulting in a block size of [1353786/1.5] m). Residuals were not explicitly analyzed for spatial autocorrelation after model fitting. |
| *Threshold selection* | · **Details on threshold selection:** Continuous maps were converted into binary 'suitable–unsuitable' maps using a threshold based on maximizing the sum of sensitivity (True Positive Rate) and specificity (True Negative Rate) (LIU *ET AL.*, 2013 ; 2016). |
| ASSESSMENT | |
| *Performance statistics* | · **Performance statistics estimated validation data:** Performance metrics are computed for all models across all parameters sets. After selecting the best models (one per algorithm for each run, based on a weighted score calculated as a weighted average of 0.5 Boyce Index, 0.25 AUC, and 0.25 Sensitivity), the values of each metric are averaged to provide an overall assessment per algorithm (with standard deviation).<br>The same approach is applied to the models selected for the ensemble model and averaged across algorithms to be used as ensemble model performance. Algorithm performances were compared using Kruskal-Wallis, with Dunn's post hoc test (Bonferroni correction) identifying pairwise differences between algorithms for each evaluation metric.<br>In the code provided, it is also possible to evaluate the ensemble model with cross-validation, but this is not executed here, as it may lead to the loss of additional species. This is because changes in the pseudo-absence/presence ratio (as all pseudo-absence data from individual models are merged) can cause previously valid blocks (containing presence records) to become empty. |
| *Plausibility check* | · **Response plots:** Response plots were generated for all 10 models per algorithm and saved in each species' folder upon code execution. No plausibility checks were conducted. |
| | · **Expert judgments:** This study builds upon a previously published framework (COLLETTE *ET AL.*, 2026), which provides the methodological basis for the analysis. The continuous output maps are available in Appendix S1-C. |
| PREDICTION | |

| | |
|---|---|
| *Prediction output* | · **Prediction units:** Bioclimatic niche suitability and hotspots maps are expressed on a continuous scale (0–1). Gain and loss of habitat were estimated from binary predictions (suitable (1)/unsuitable (0)). |
| *Uncertainty quantification* | · **Algorithmic uncertainty:** A 5% divergence threshold was applied to identify pixels showing significant disagreement among the predictions of the algorithms retained for the median ensemble model. This calculation provides an indication of where predictions are less consistent between algorithms; these pixels were masked on the current distribution map to focus on more reliable areas. |
| | · **Uncertainty in scenarios:** No quantification of scenario uncertainty was performed. |
| | · **Visualization/treatment of novel environments:** NA |

**Appendix C: Current and future continuous bioclimatic niche suitability maps for invasive plants in the Pyrenees.**

This appendix provides continuous maps of bioclimatic niche suitability for 46 plant species recognized as invasive on both sides of the Pyrenees, modeled under current and future climatic conditions. Suitability is represented as a gradient from blue (low bioclimatic suitability) to yellow (high bioclimatic suitability). The maps are displayed in alphabetical order by species name.

List of species with displayed maps:

*Acacia dealbata, Acer negundo, Agave americana, Ailanthus altissima, Araujia sericifera, Artemisia verlotiorum, Azolla filiculoides, Bidens frondosa, Bidens subalternans, Buddleja davidii, Carpobrotus acinaciformis, Carpobrotus edulis, Cortaderia selloana, Cyperus eragrostis, Elaeagnus angustifolia, Epilobium ciliatum, Erigeron canadensis, Erigeron karvinskianus, Erigeron sumatrensis, Euphorbia prostrata, Fallopia baldschuanica, Gleditsia triacanthos, Helianthus tuberosus, Impatiens balfourii, Impatiens glandulifera, Ligustrum lucidum, Lonicera japonica, Ludwigia peploides, Opuntia stricta, Parthenocissus inserta, Paspalum dilatatum, Periploca graeca, Phyllostachys aurea, Phyllostachys bambusoides, Phyllostachys nigra, Phytolacca americana, Prunus cerasifera, Robinia pseudoacacia, Senecio inaequidens, Sicyos angulatus, Solidago canadensis, Sorghum halepense, Sporobolus indicus, Symphyotrichum pilosum, Symphyotrichum squamatum, Xanthium orientale*

For each species, we provide:

- Current bioclimatic niche suitability based on current climatic conditions (1970-2000), with areas of uncertainty (standard deviation > 5% between models) masked in black. Maps are cropped to the Pyrenees, while model calibration was performed over a broader European extent (xmin = −11, xmax = 19.5, ymin = 35.7, ymax = 63.5), provided upon request.
- Future projections for four periods (2021-2040, 2041-2060, 2061-2080, 2081-2100 designated as 2030, 2050, 2070, 2090), under four Shared Socioeconomic Pathways (SSP126, SSP245, SSP370, SSP585, best to worst-case climate scenarios, respectively).

Current bioclimatic suitability in the Pyrenees for *Acacia dealbata*

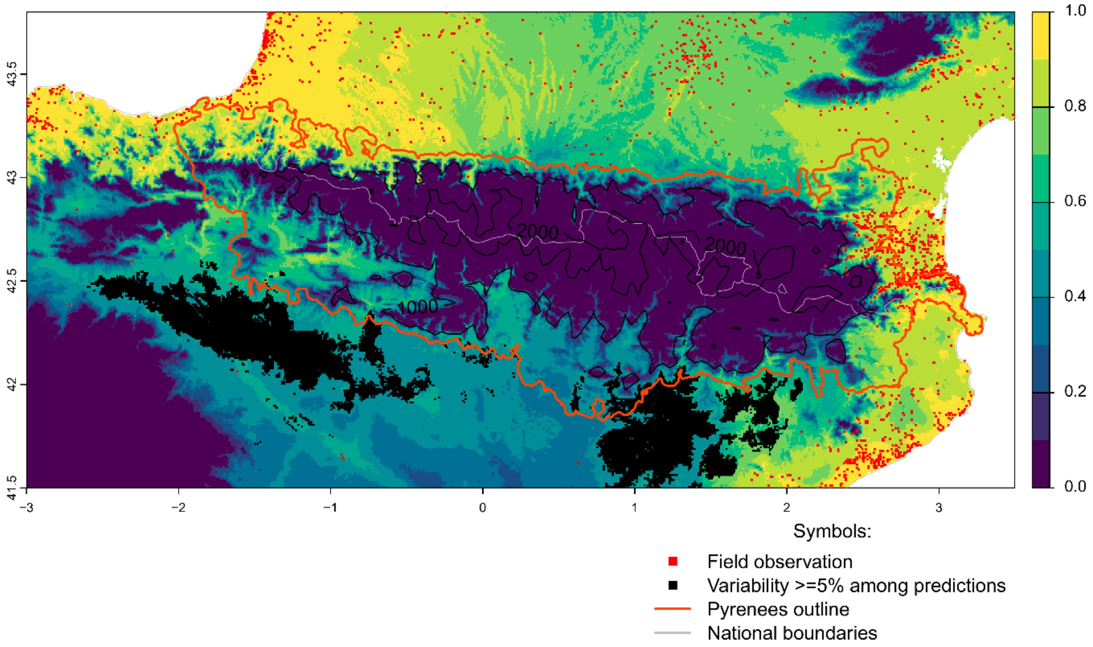


Future bioclimatic suitability in the Pyrenees by 2090 for *Acacia dealbata*

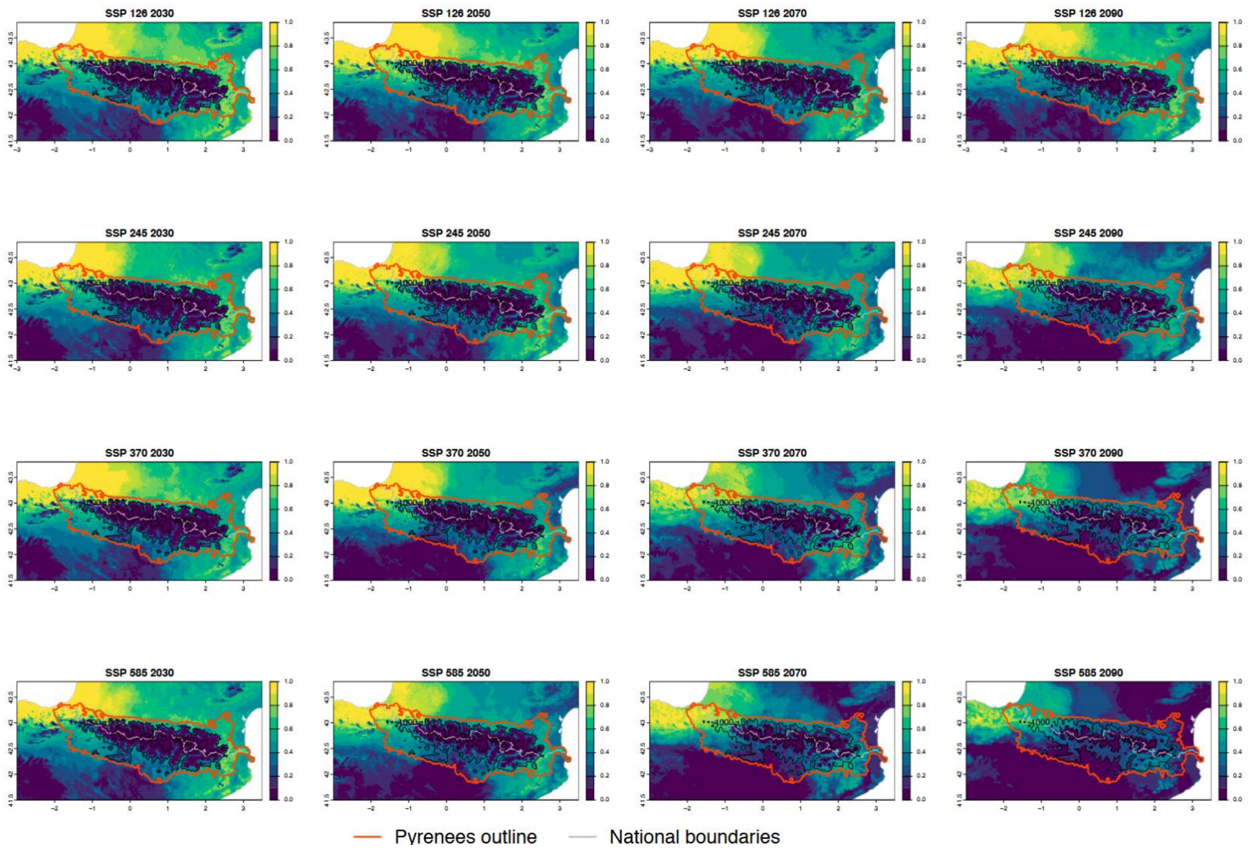

Current bioclimatic suitability in the Pyrenees for *Acer negundo*

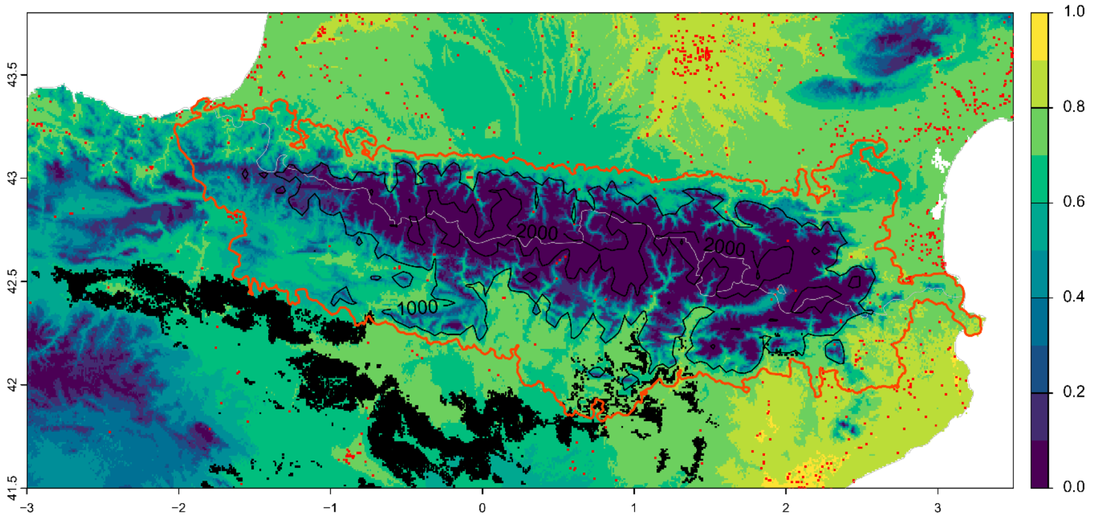


Symbols:

- Field observation
- Variability >=5% among predictions
- Pyrenees outline
- National boundaries

Future bioclimatic suitability in the Pyrenees by 2090 for *Acer negundo*

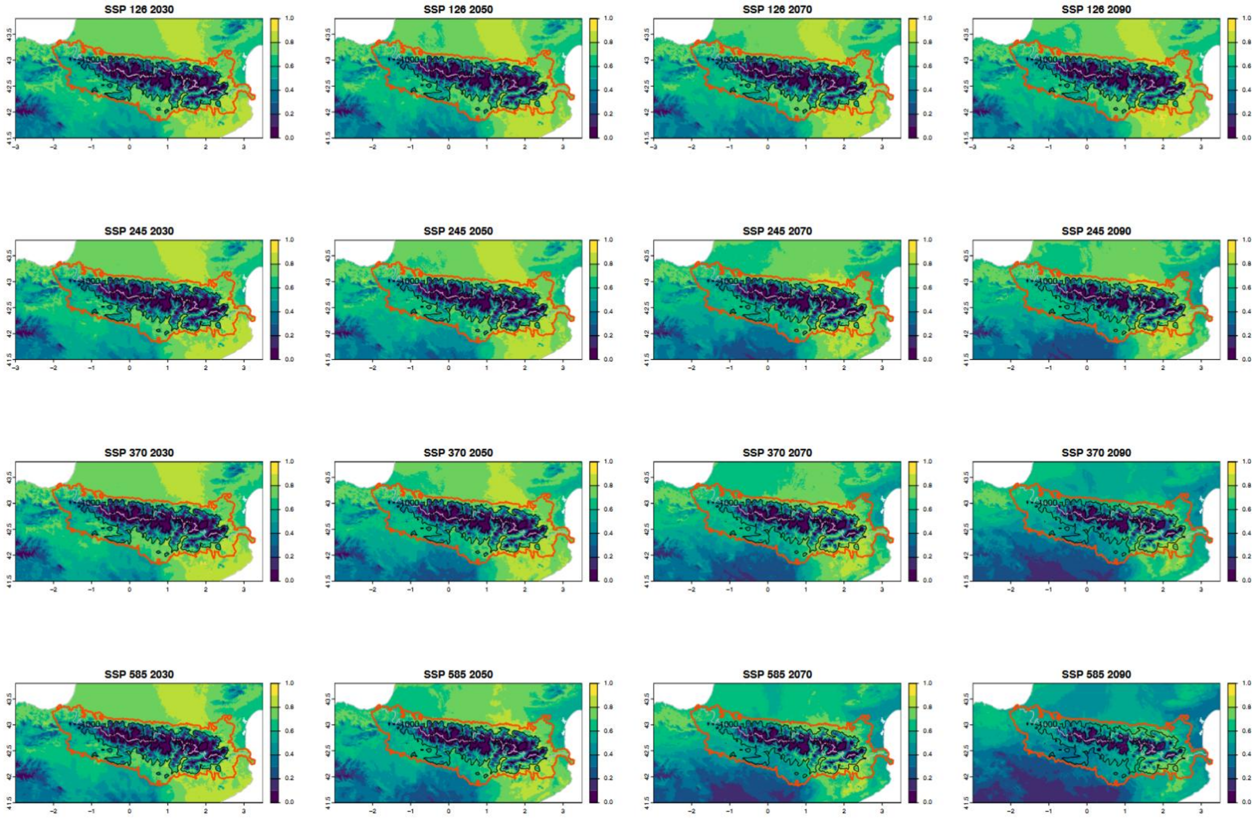


Pyrenees outline National boundaries

Current bioclimatic suitability in the Pyrenees for *Agave americana*

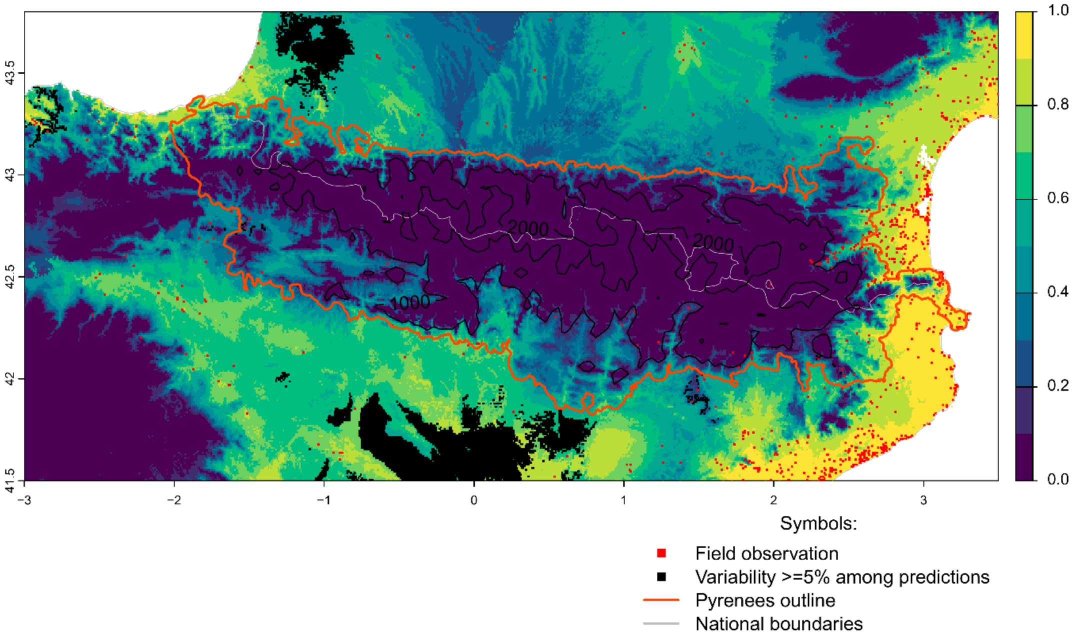


Future bioclimatic suitability in the Pyrenees by 2090 for *Agave americana*

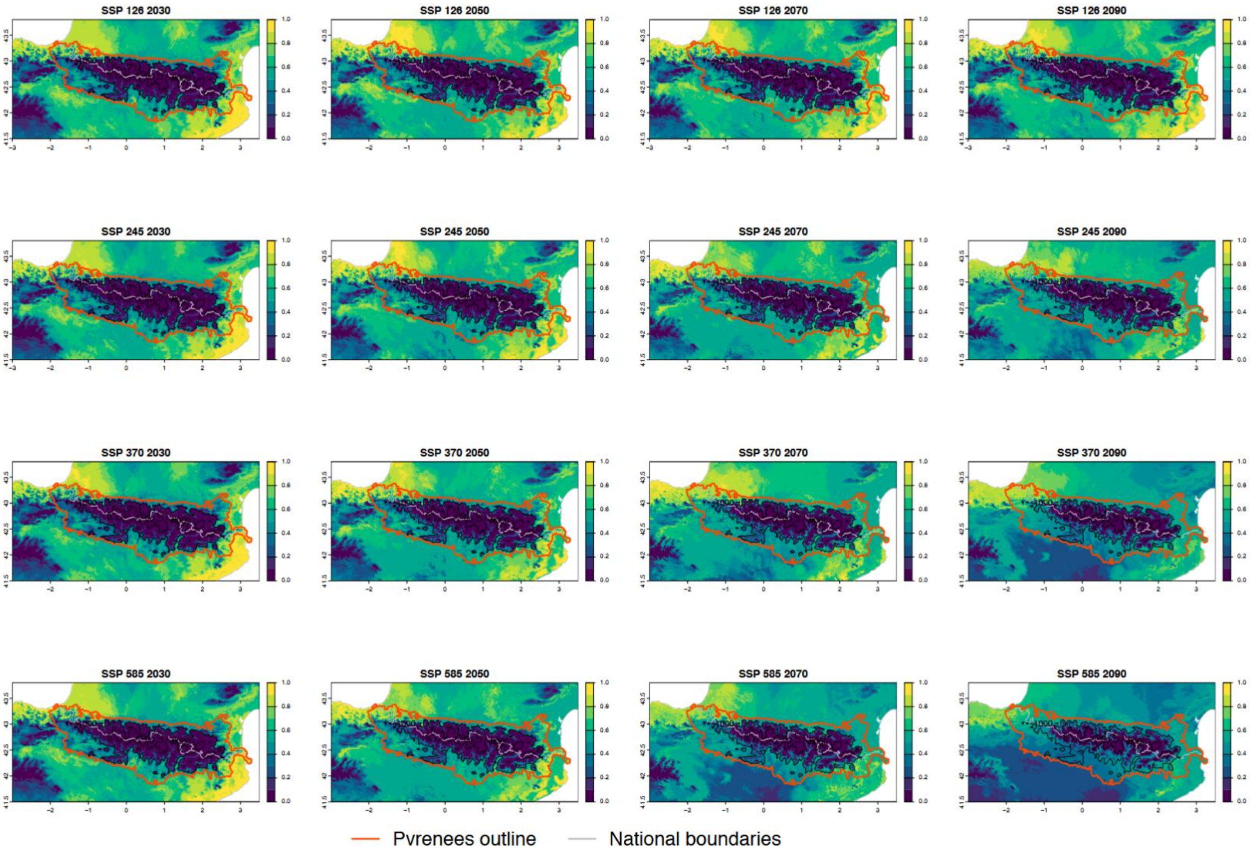

Current bioclimatic suitability in the Pyrenees for *Ailanthus altissima*

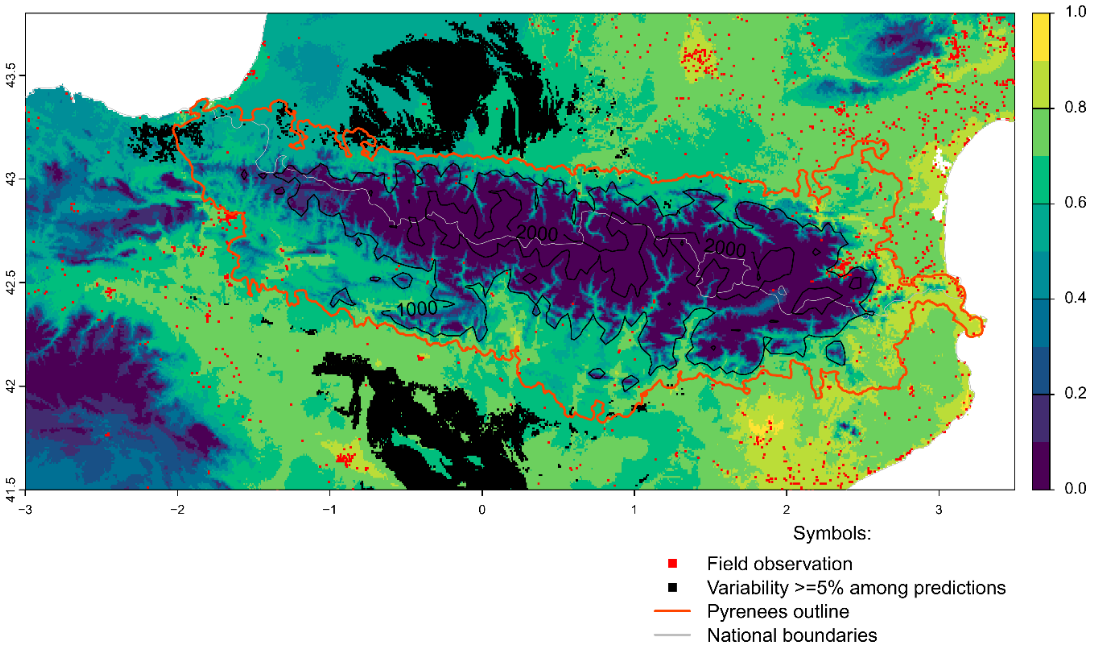


Future bioclimatic suitability in the Pyrenees by 2090 for *Ailanthus altissima*

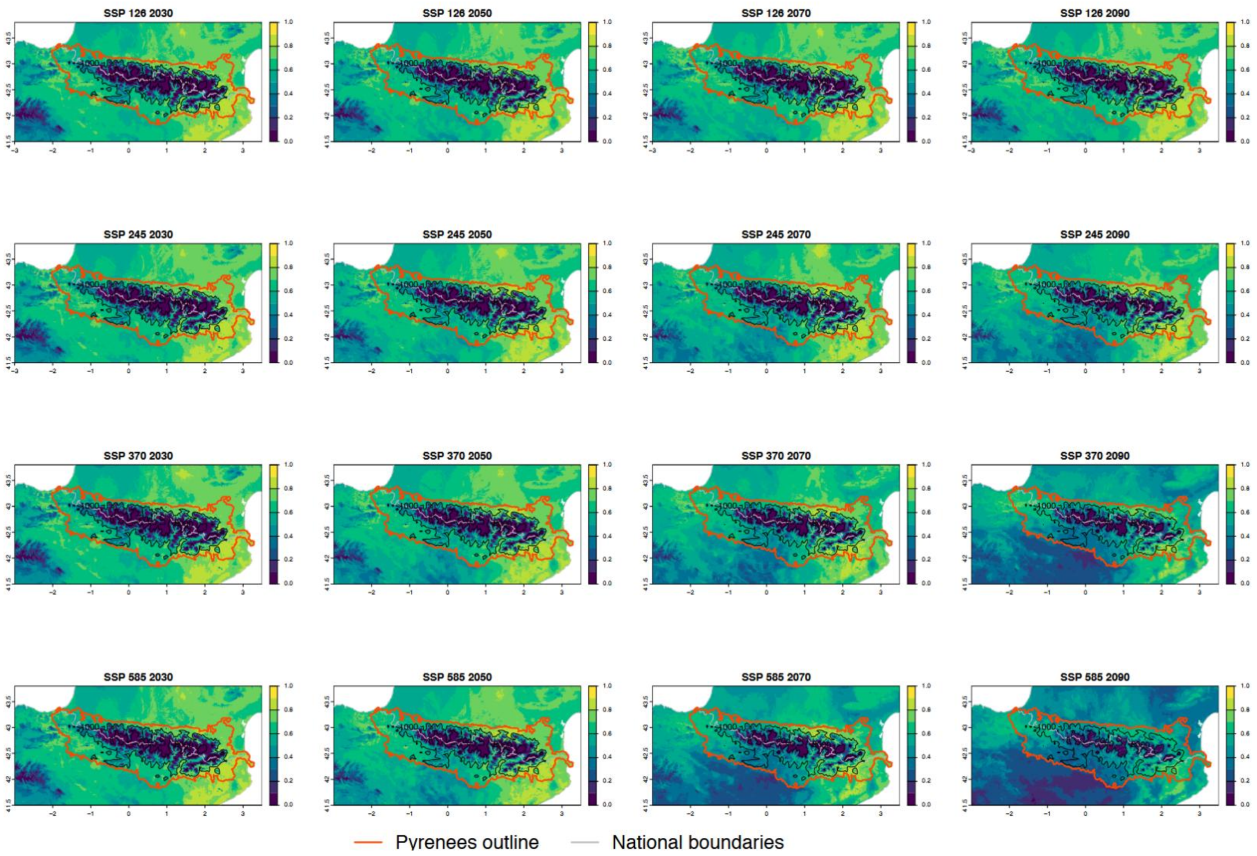

Current bioclimatic suitability in the Pyrenees for *Araujia sericifera*

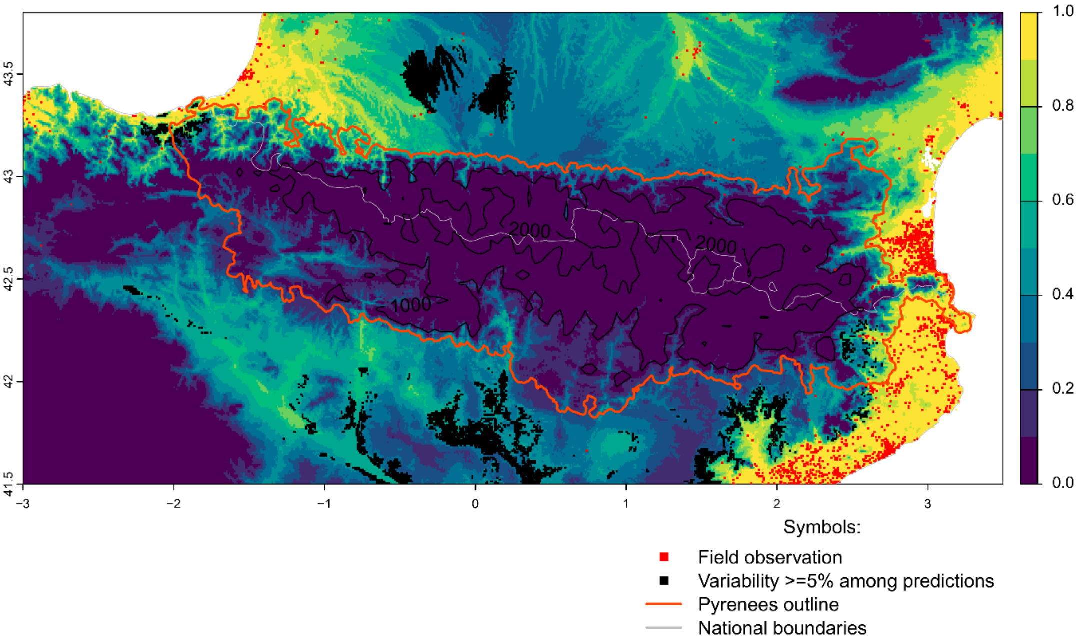


Future bioclimatic suitability in the Pyrenees by 2090 for *Araujia sericifera*

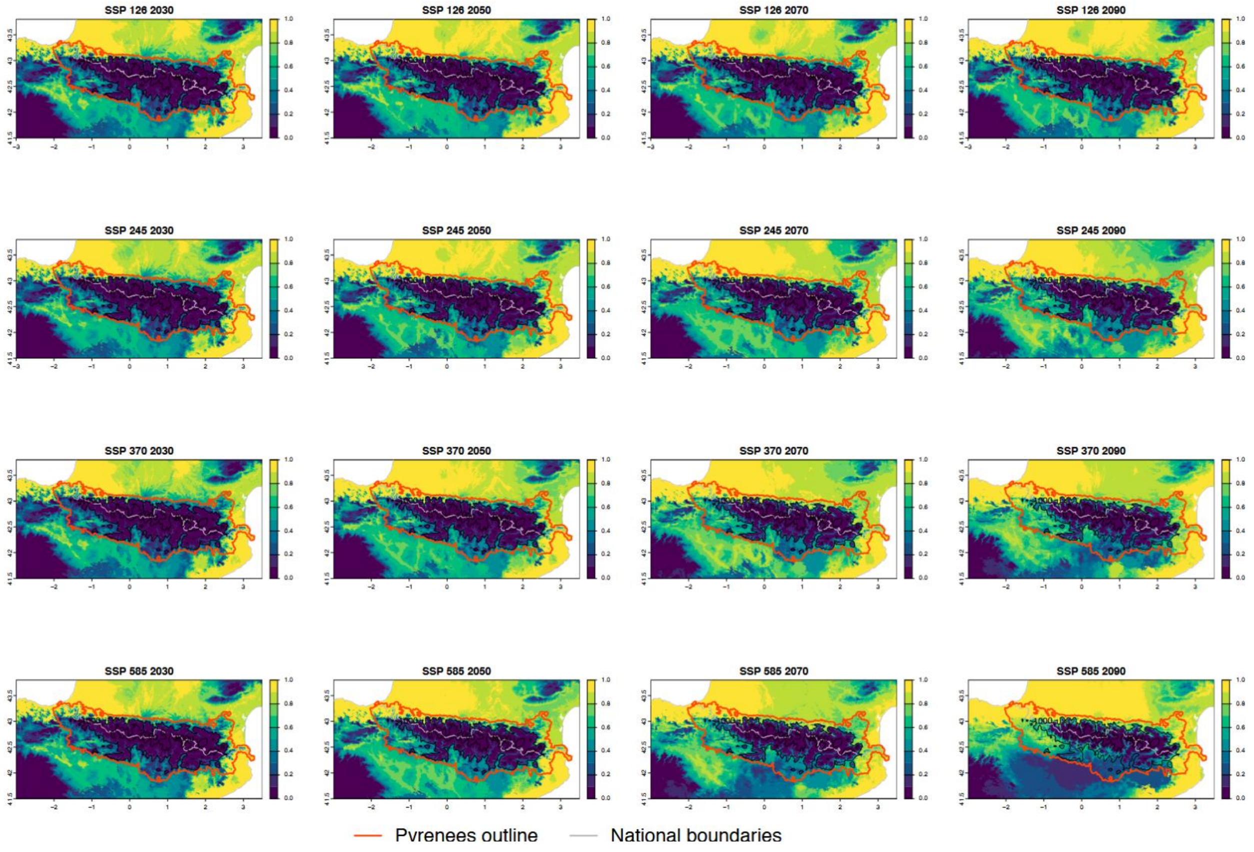

Current bioclimatic suitability in the Pyrenees for *Artemisia verlotiorum*

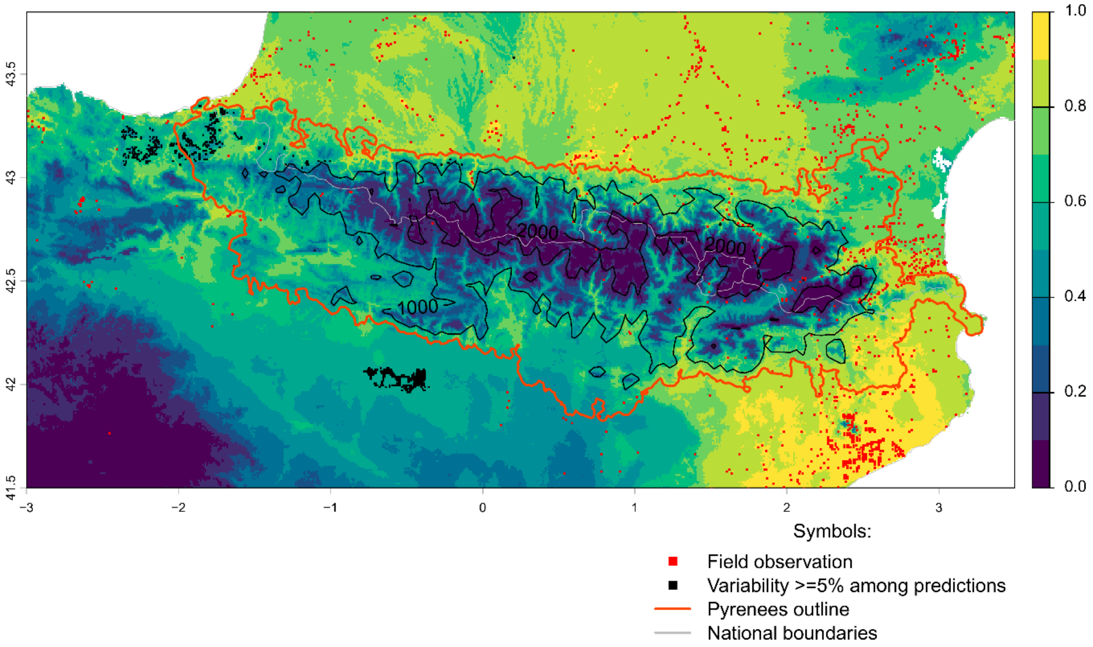


Future bioclimatic suitability in the Pyrenees by 2090 for *Artemisia verlotiorum*

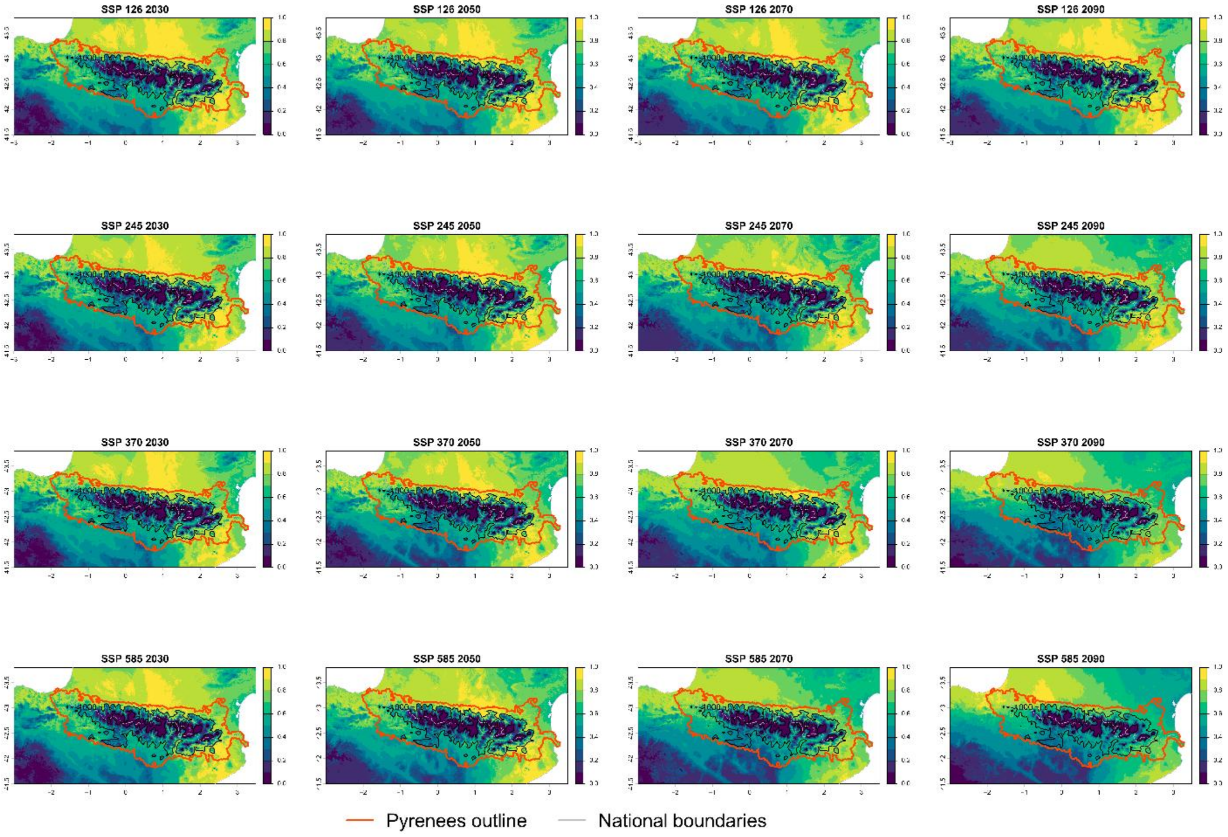

Current bioclimatic suitability in the Pyrenees for *Azolla filiculoides*

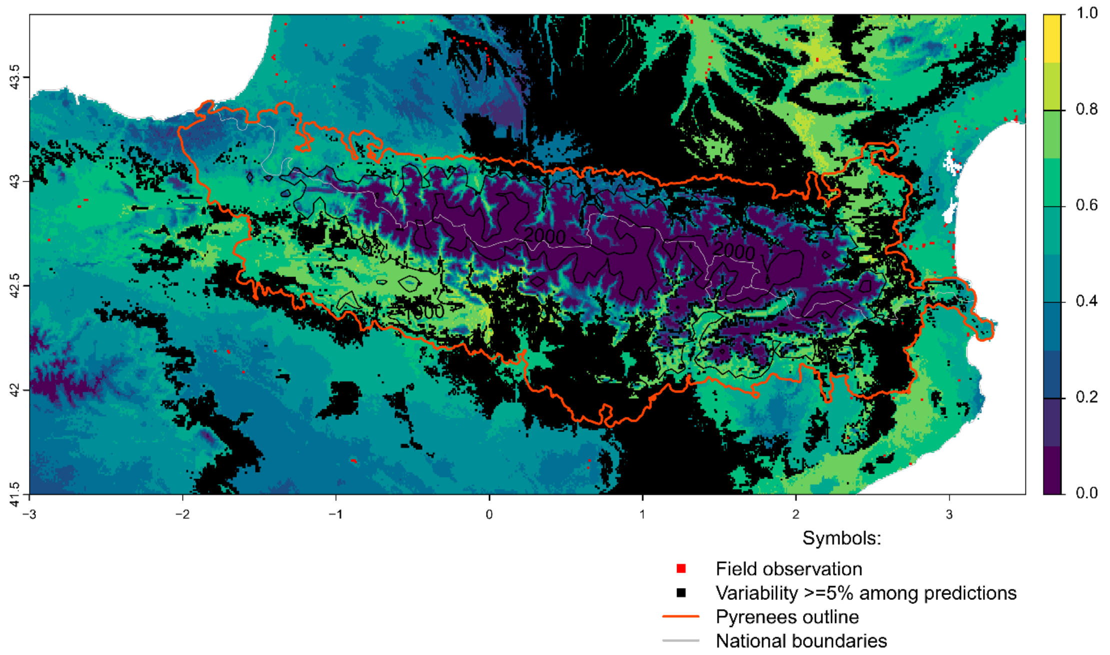


Future bioclimatic suitability in the Pyrenees by 2090 for *Azolla filiculoides*

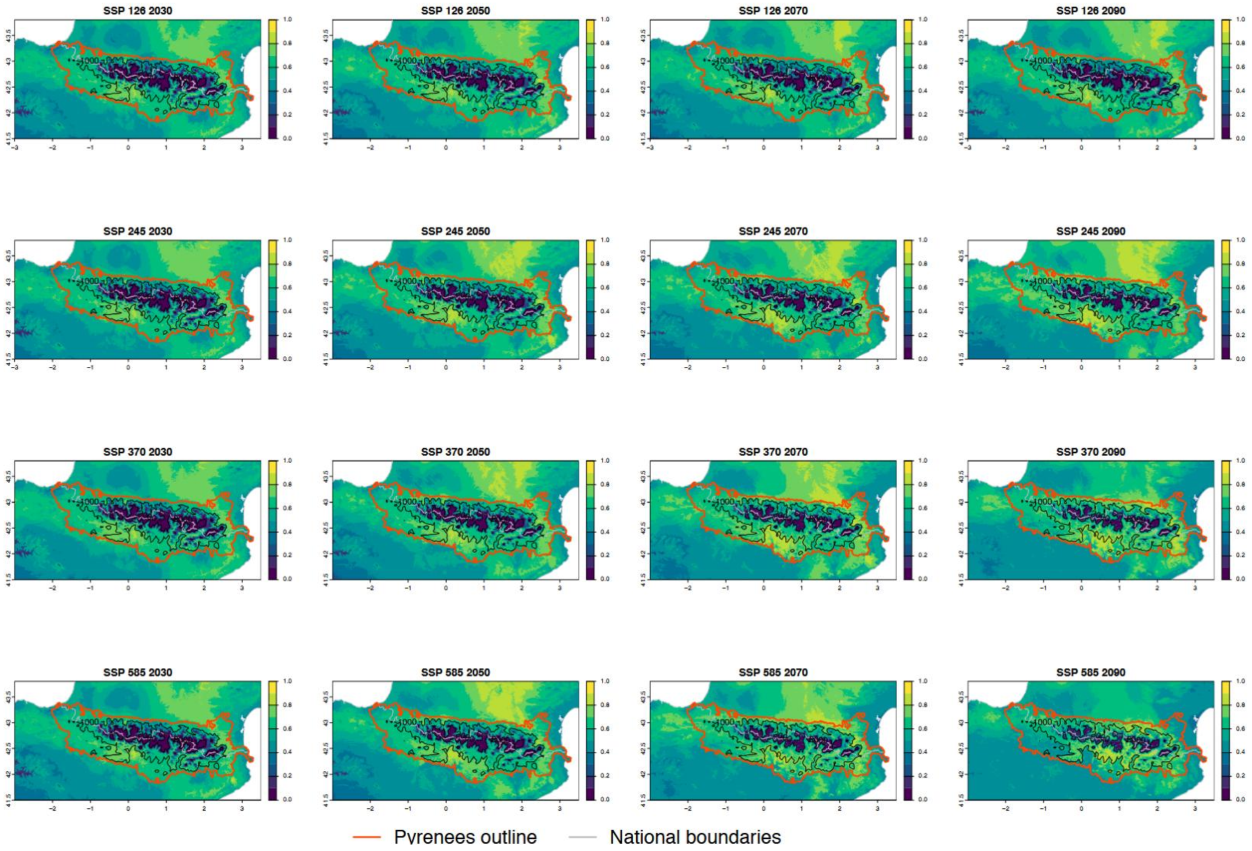

Current bioclimatic suitability in the Pyrenees for *Bidens frondosa*

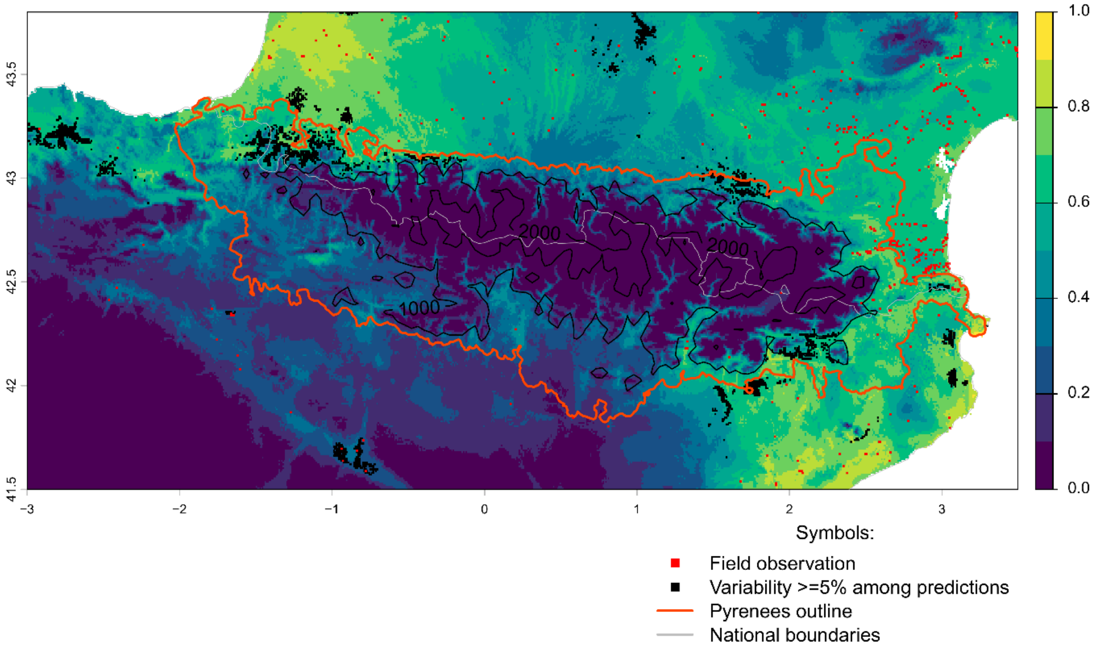


Future bioclimatic suitability in the Pyrenees by 2090 for *Bidens frondosa*

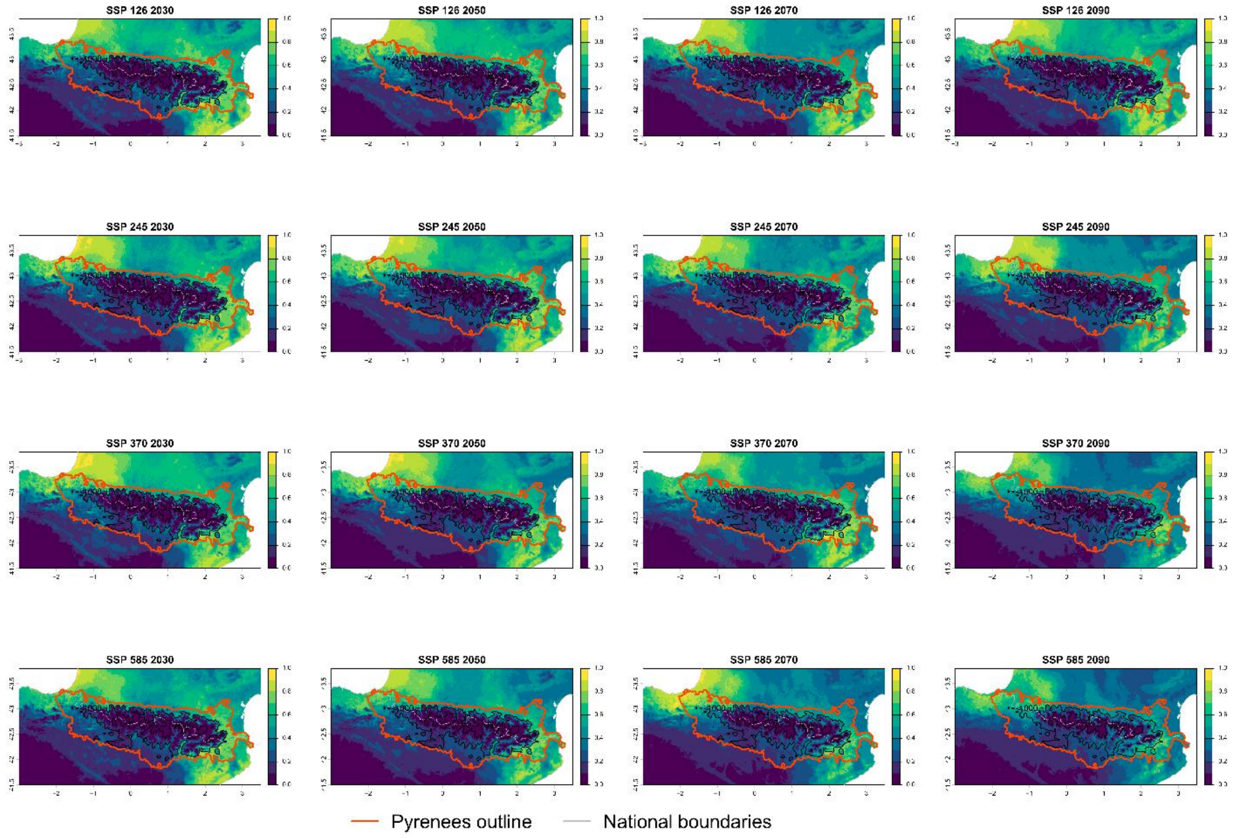

Current bioclimatic suitability in the Pyrenees for *Bidens subalternans*

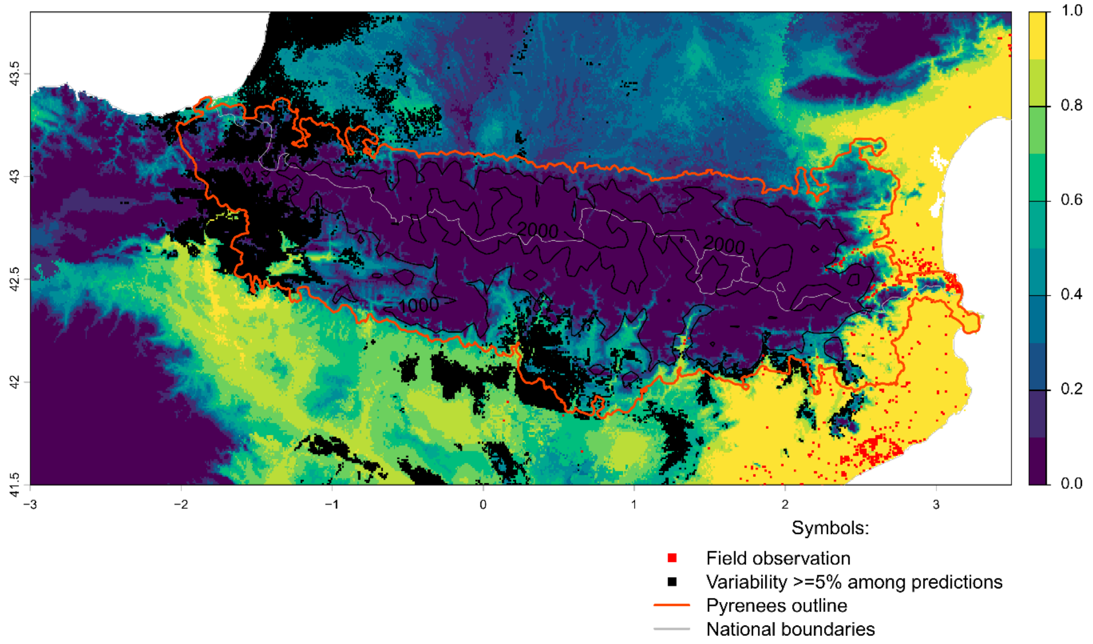


Future bioclimatic suitability in the Pyrenees by 2090 for *Bidens subalternans*

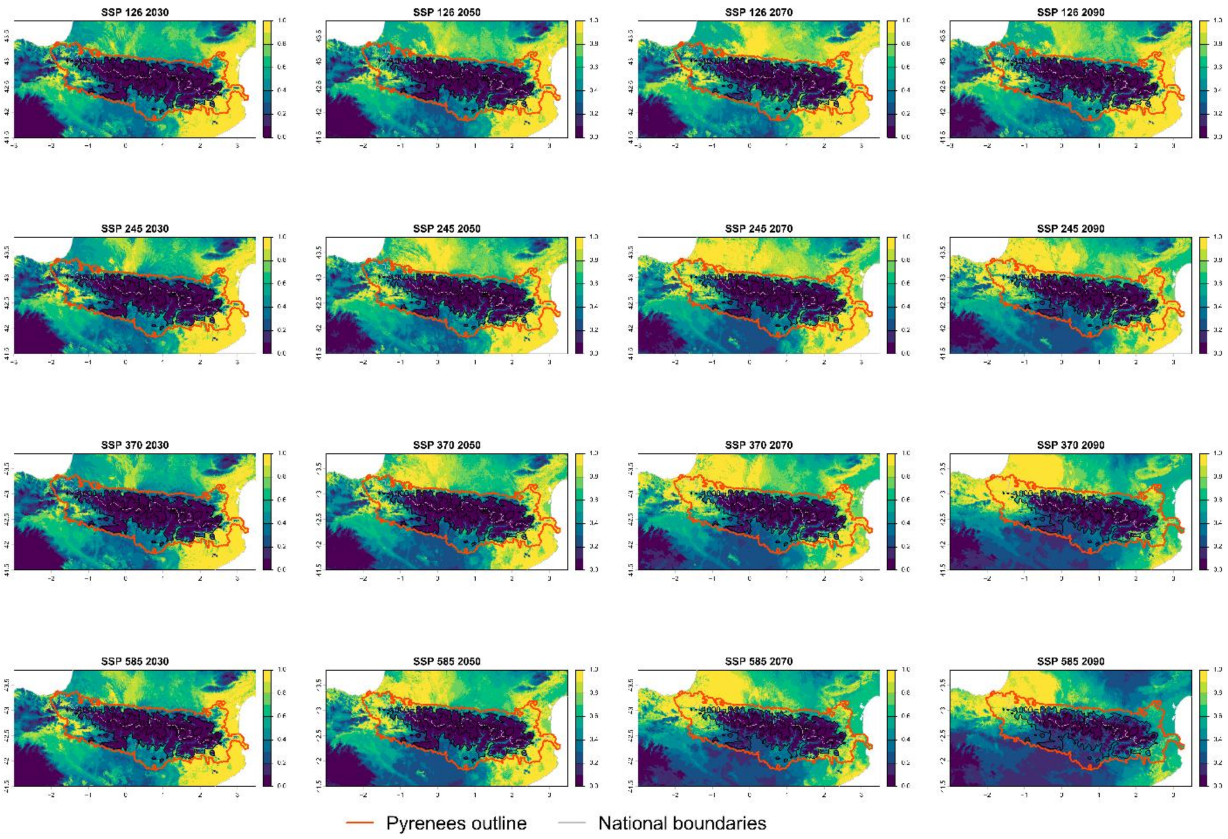

Current bioclimatic suitability in the Pyrenees for *Buddleja davidii*

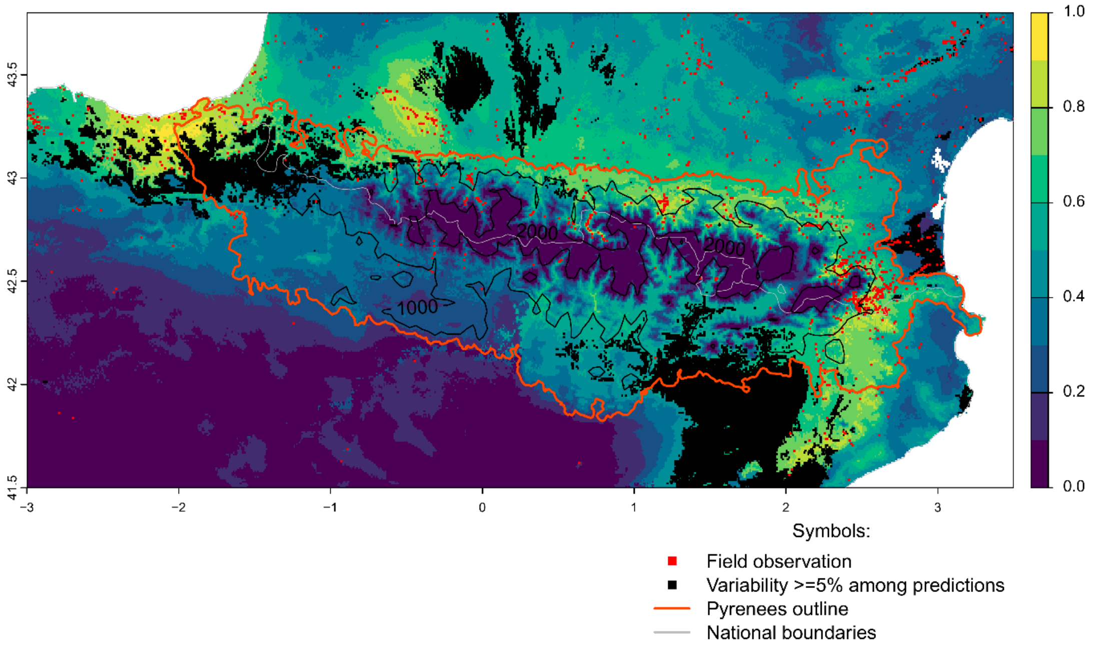


Future bioclimatic suitability in the Pyrenees by 2090 for *Buddleja davidii*

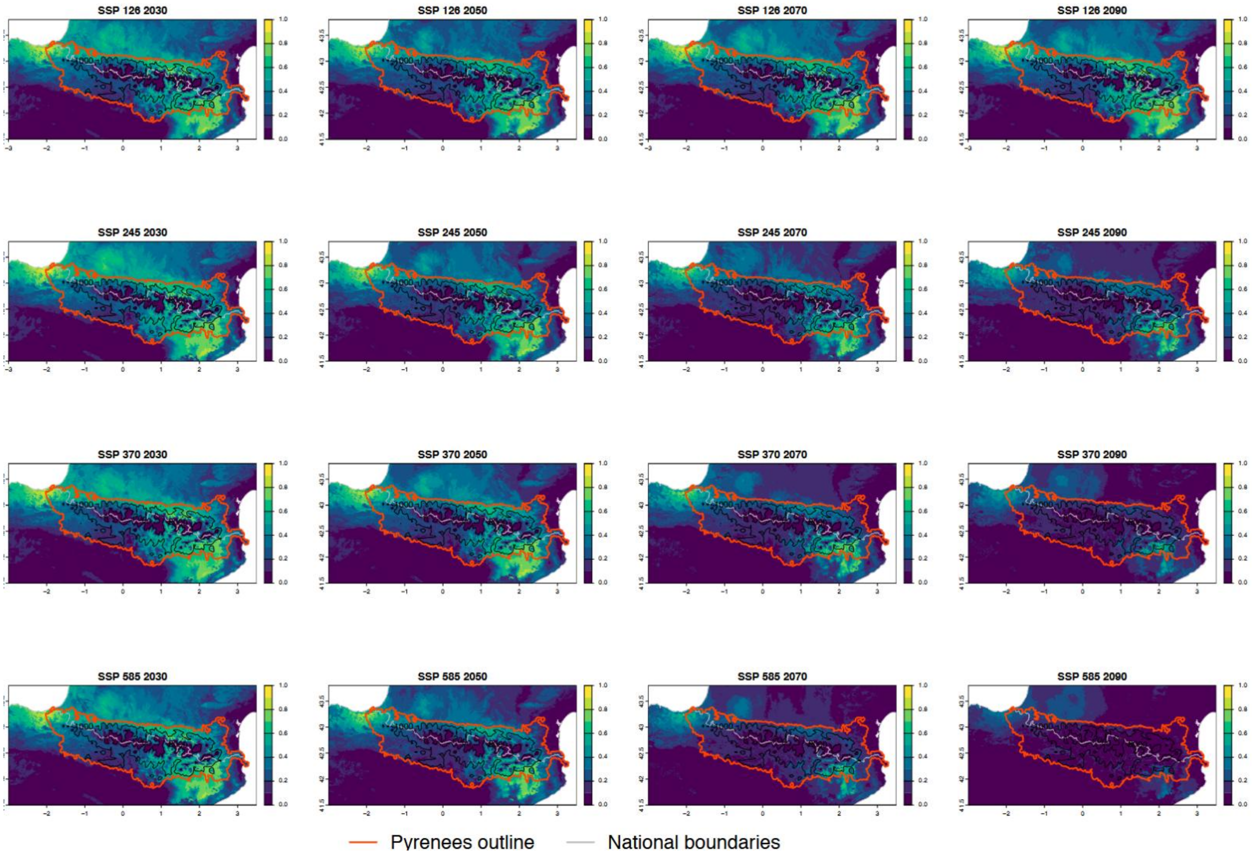

Current bioclimatic suitability in the Pyrenees for *Carpobrotus acinaciformis*

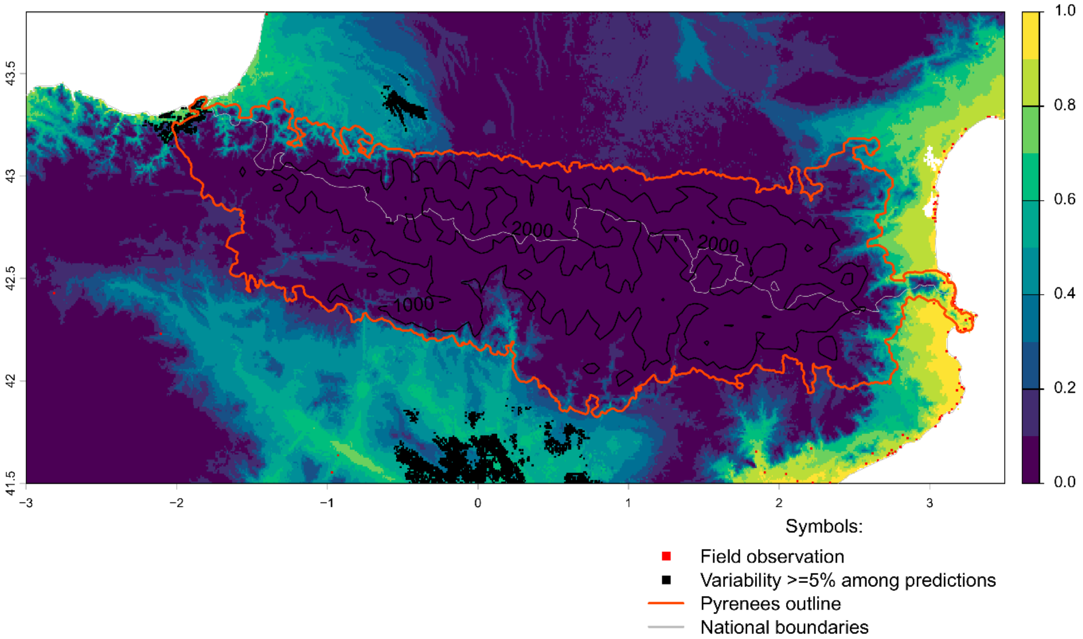


Future bioclimatic suitability in the Pyrenees by 2090 for *Carpobrotus acinaciformis*

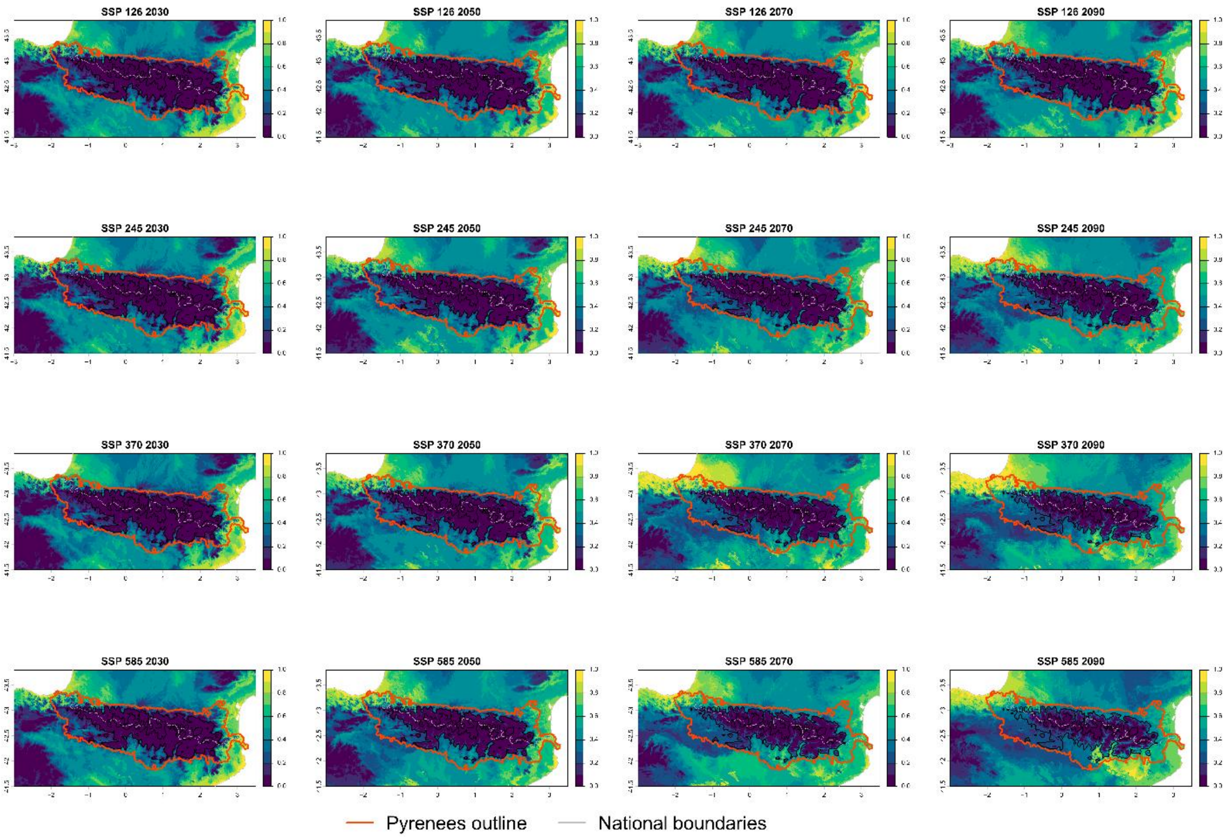

Current bioclimatic suitability in the Pyrenees for *Carpobrotus edulis*

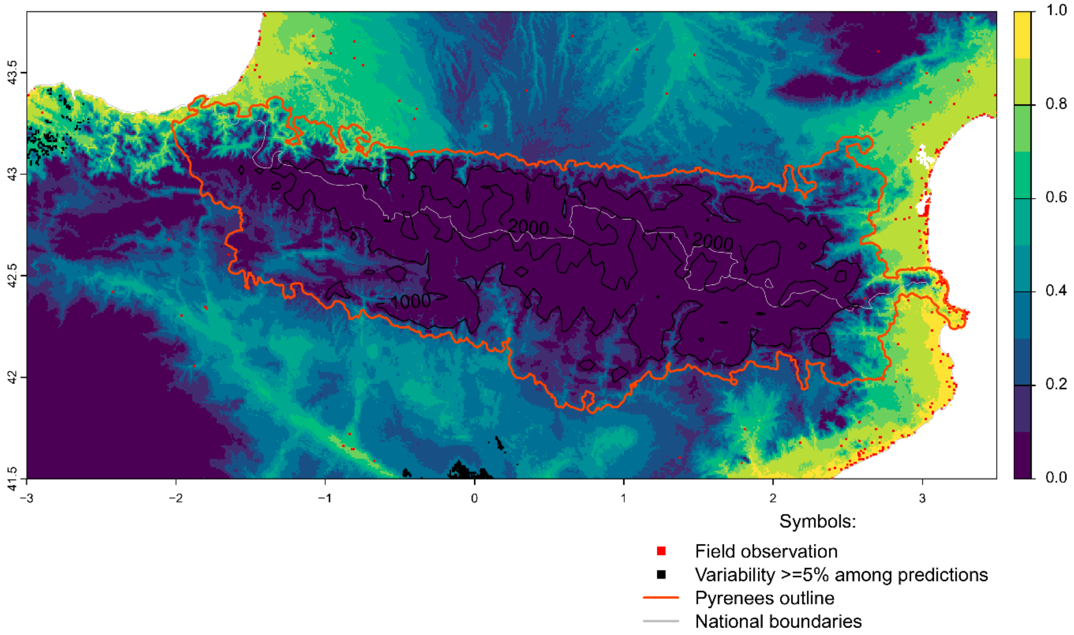


Future bioclimatic suitability in the Pyrenees by 2090 for *Carpobrotus edulis*

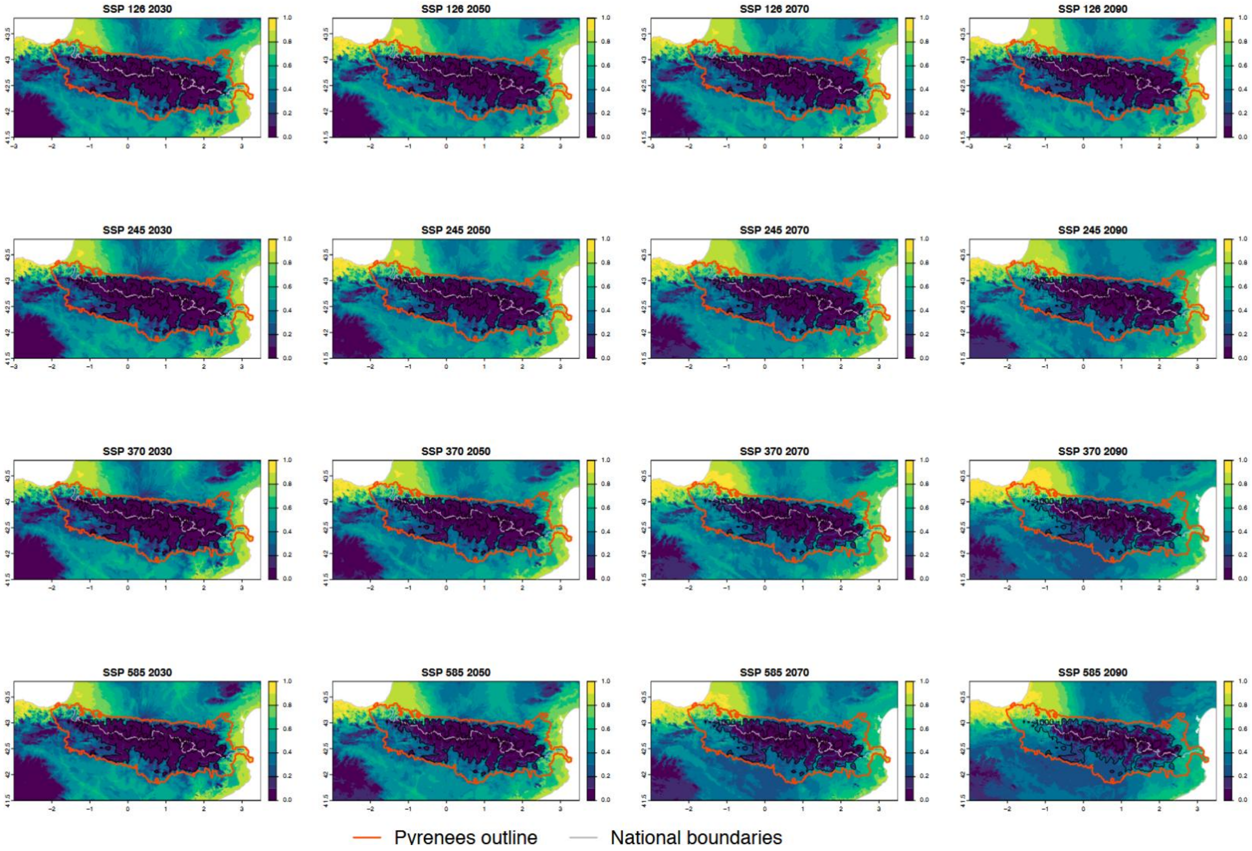

Current bioclimatic suitability in the Pyrenees for *Cortaderia selloana*

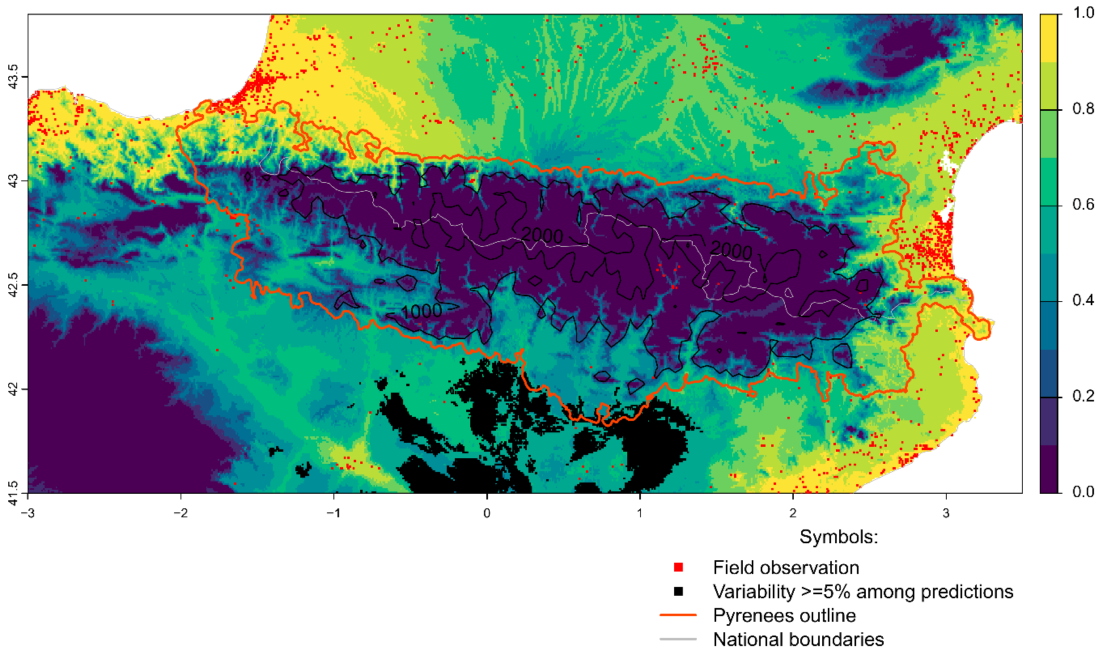


Future bioclimatic suitability in the Pyrenees by 2090 for *Cortaderia selloana*

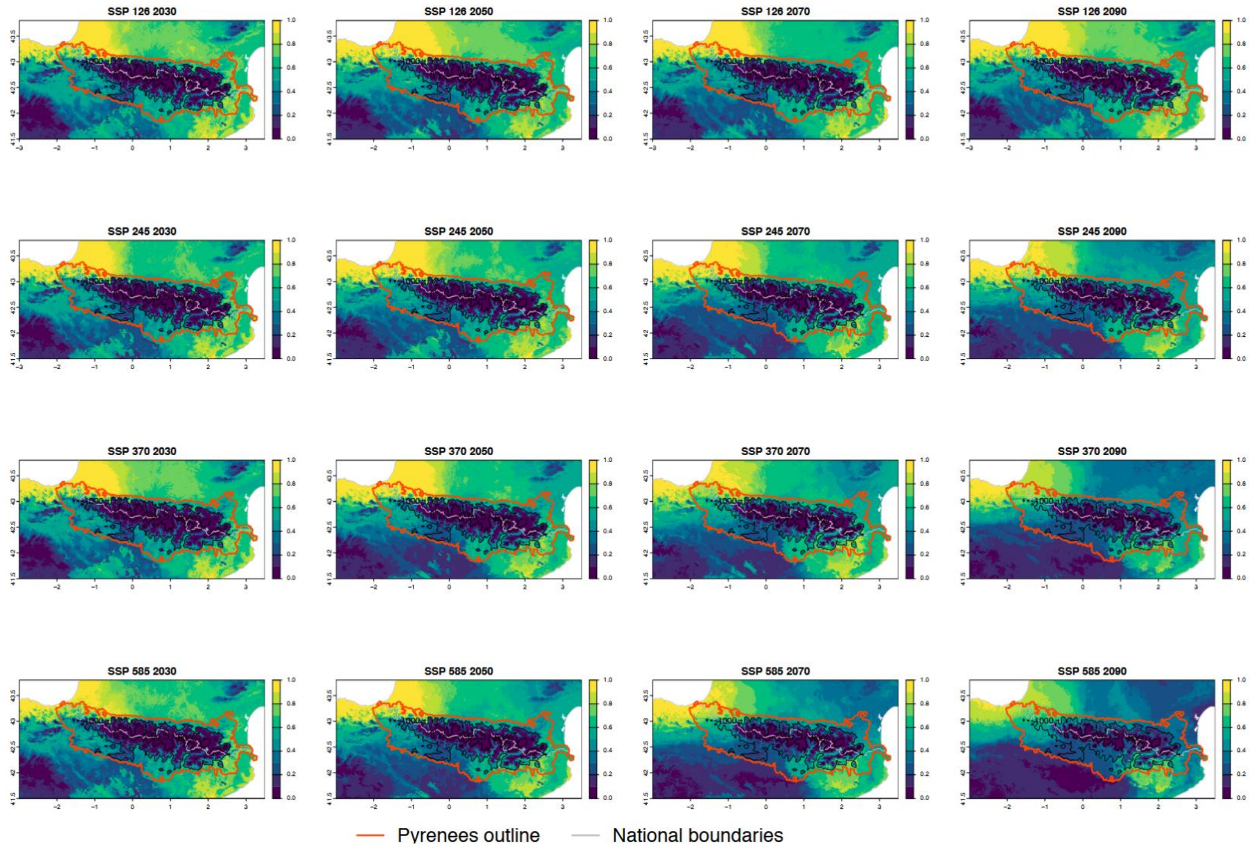

Current bioclimatic suitability in the Pyrenees for *Cyperus eragrostis*

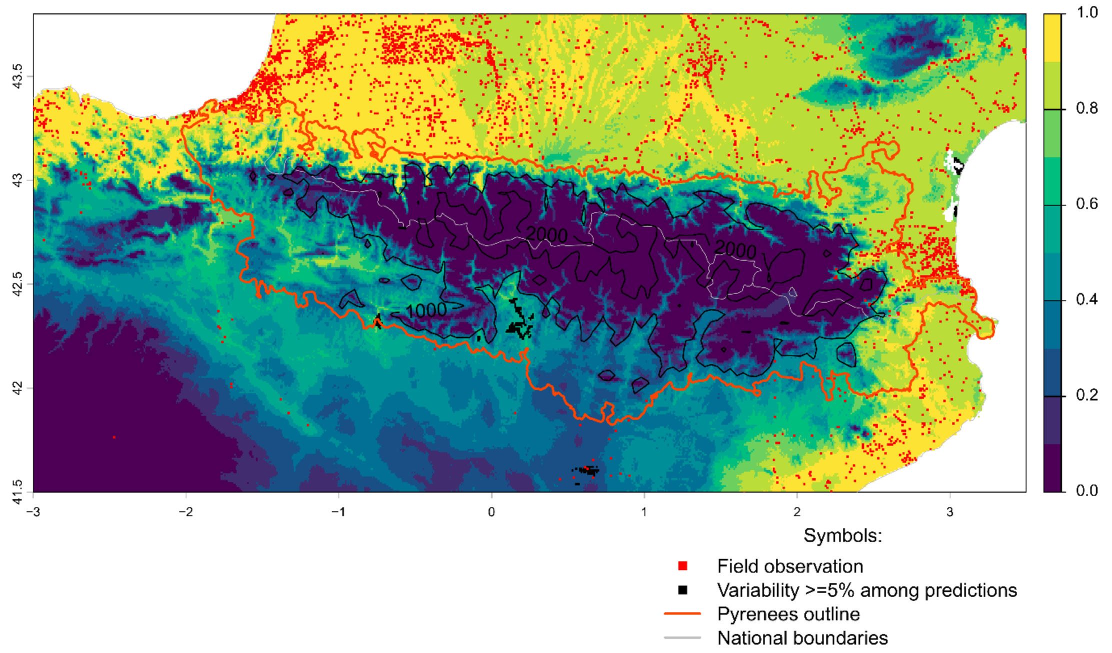


Future bioclimatic suitability in the Pyrenees by 2090 for *Cyperus eragrostis*

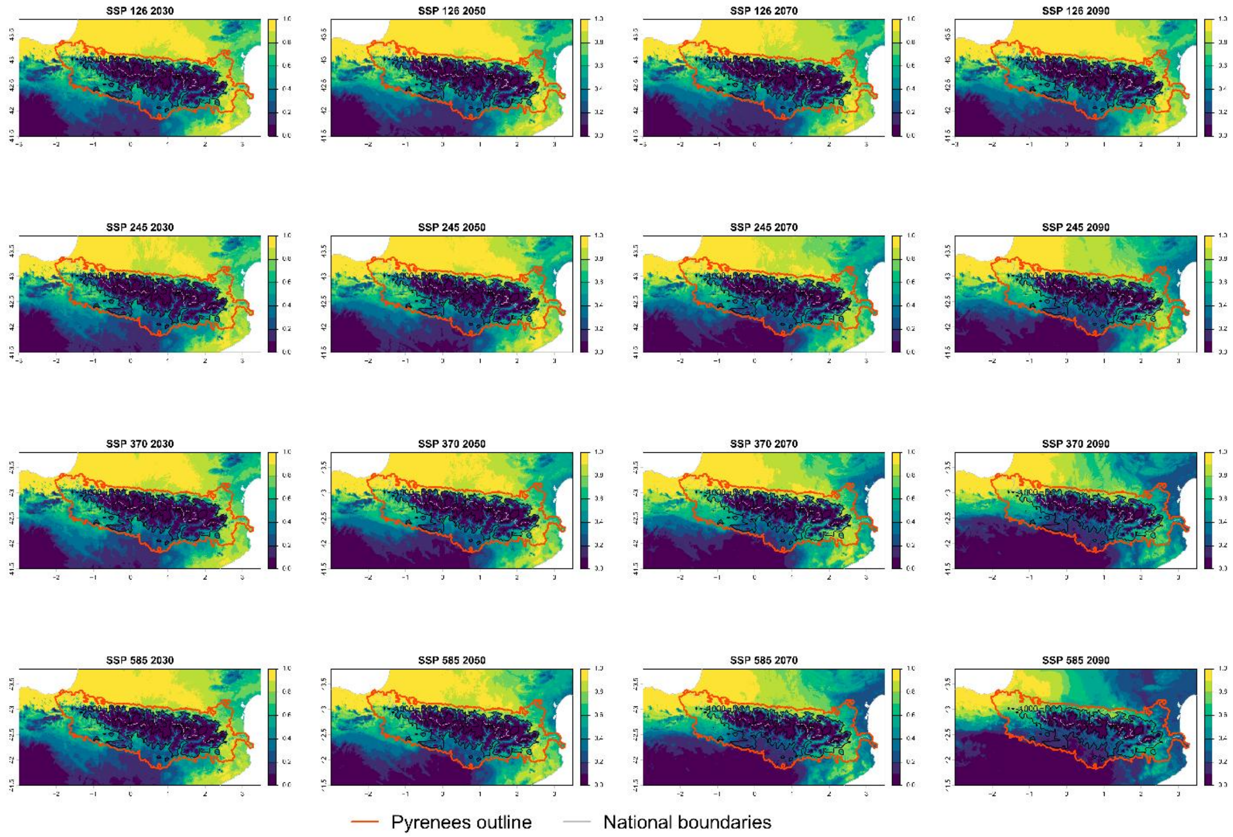

### Current bioclimatic suitability in the Pyrenees for *Elaeagnus angustifolia*

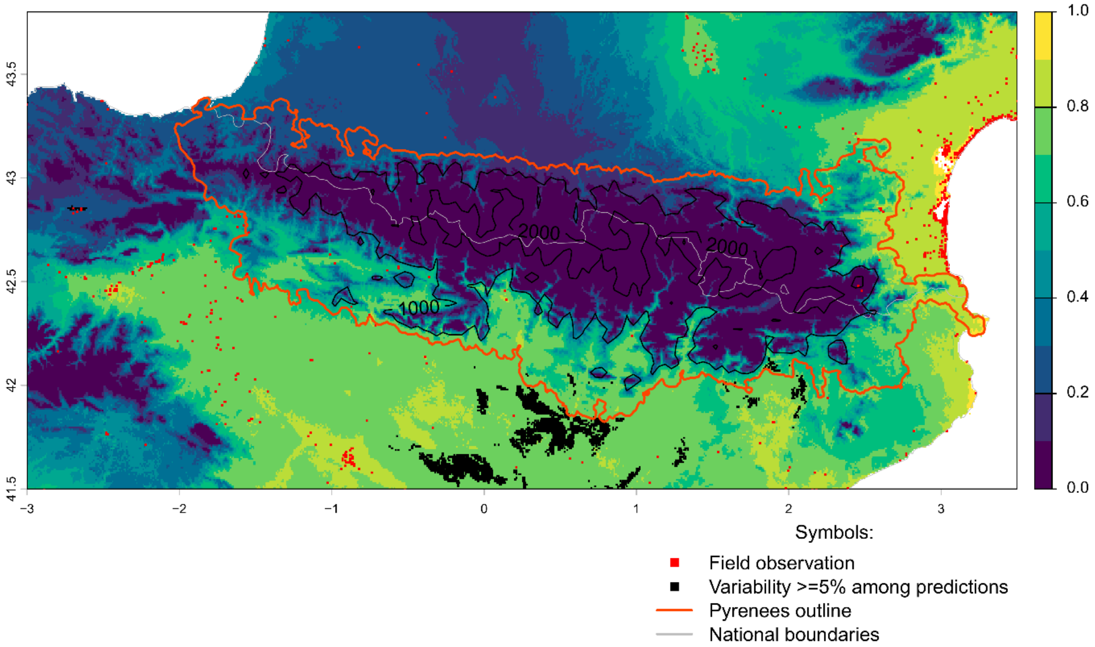


### Future bioclimatic suitability in the Pyrenees by 2090 for *Elaeagnus angustifolia*

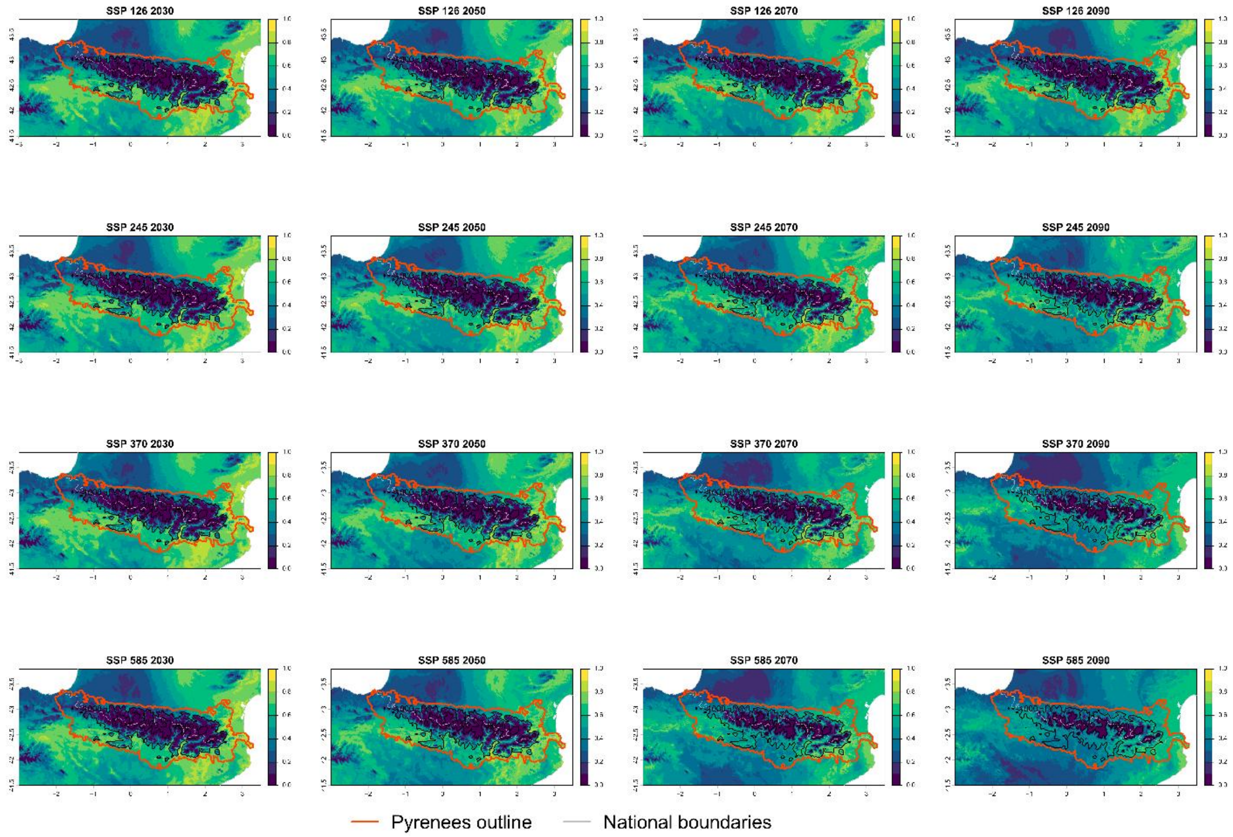

### Current bioclimatic suitability in the Pyrenees for *Epilobium ciliatum*

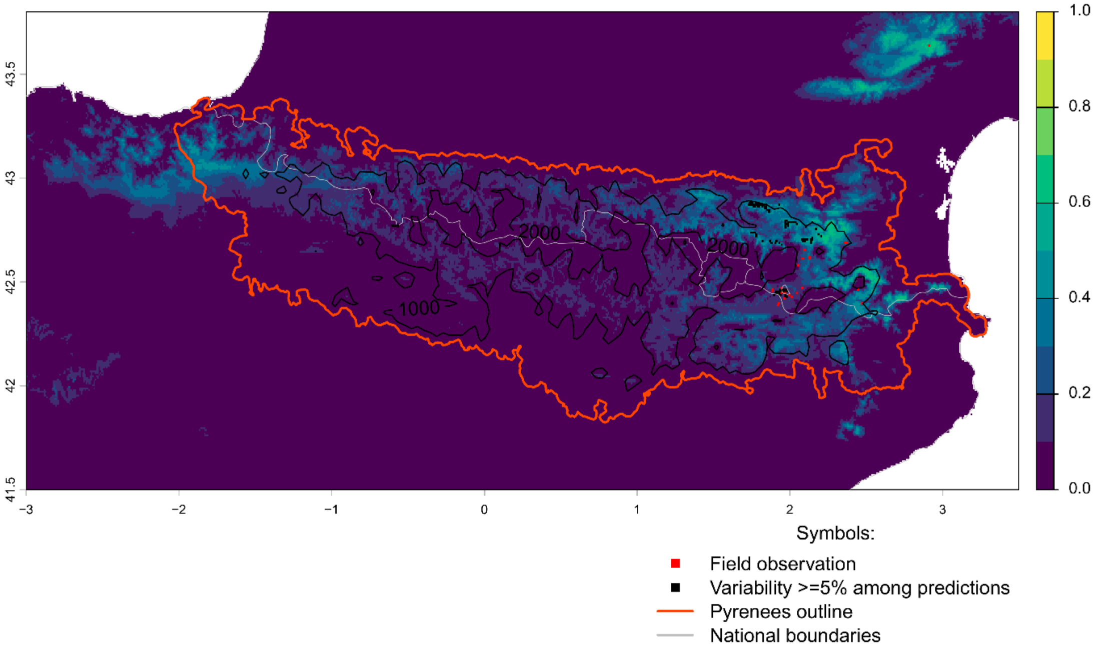


Future bioclimatic suitability in the Pyrenees by 2090 for *Epilobium ciliatum*

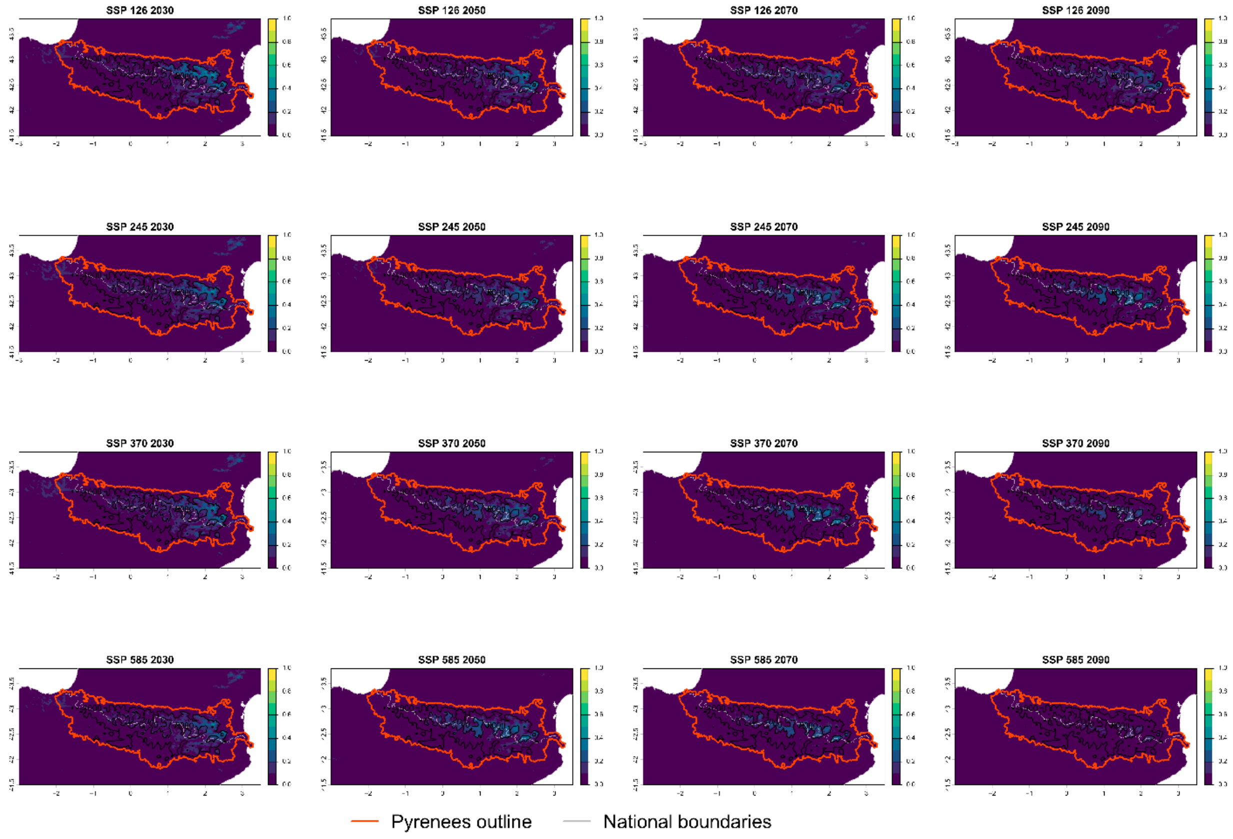

Current bioclimatic suitability in the Pyrenees for *Erigeron canadensis*

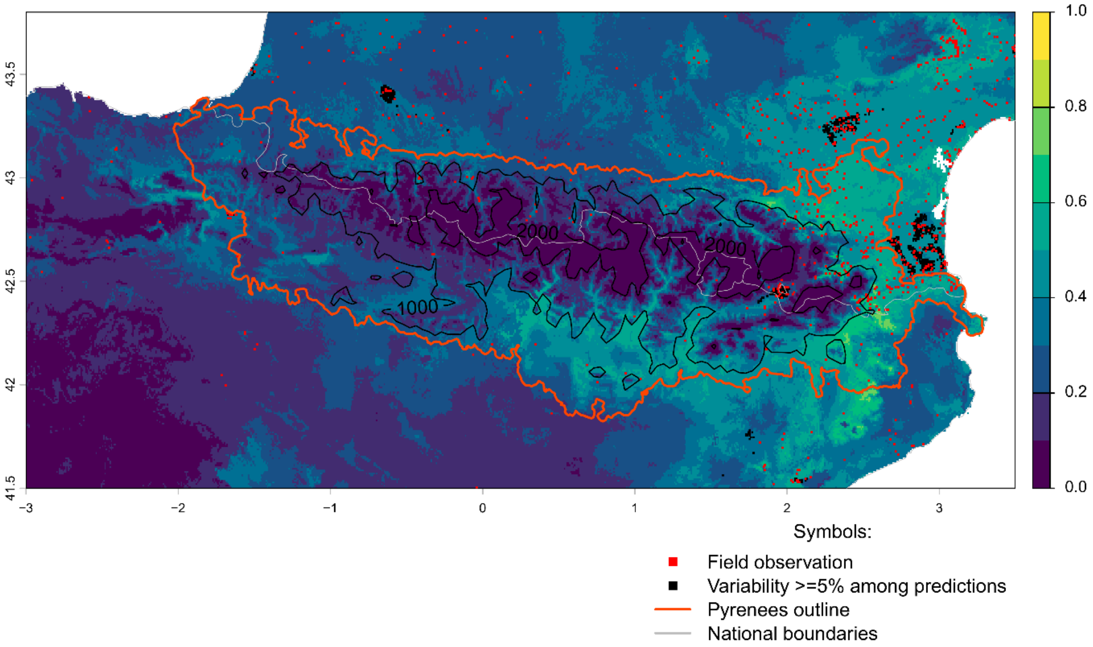


Future bioclimatic suitability in the Pyrenees by 2090 for *Erigeron canadensis*

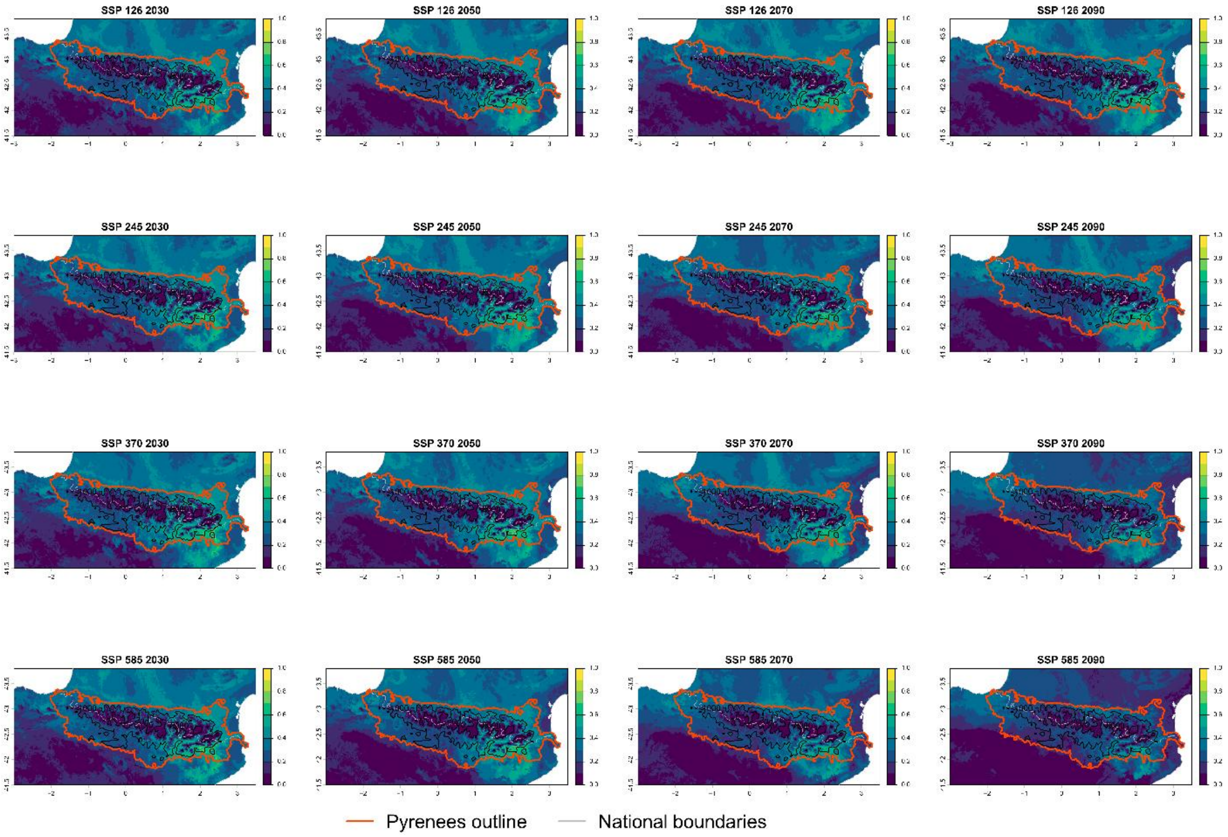

Current bioclimatic suitability in the Pyrenees for *Erigeron karvinskianus*

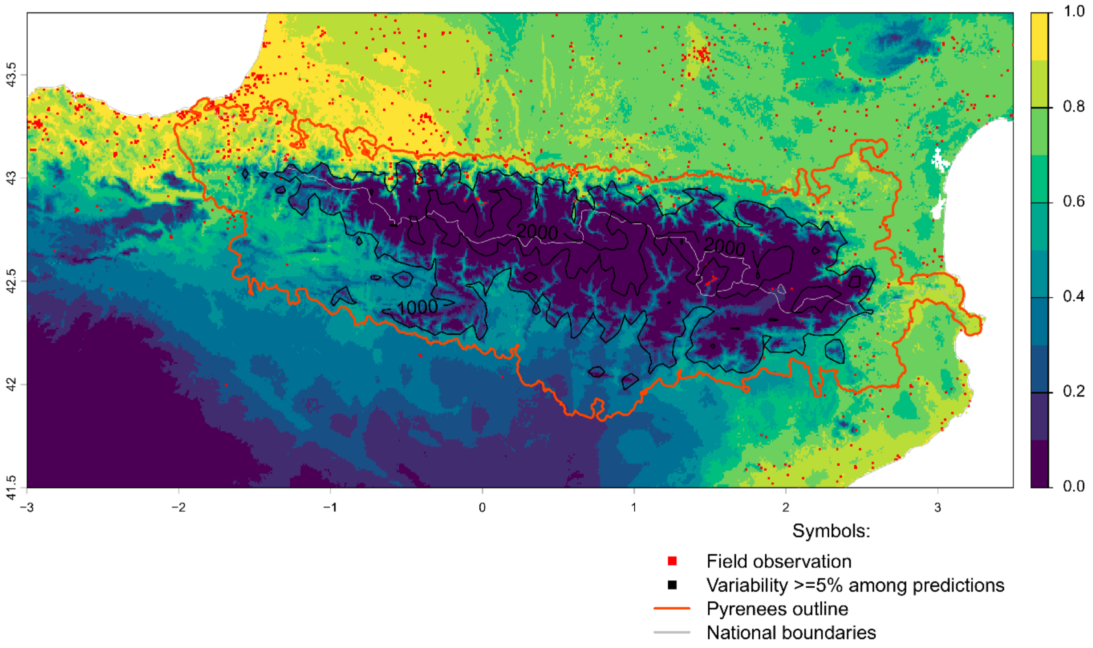


Future bioclimatic suitability in the Pyrenees by 2090 for *Erigeron karvinskianus*

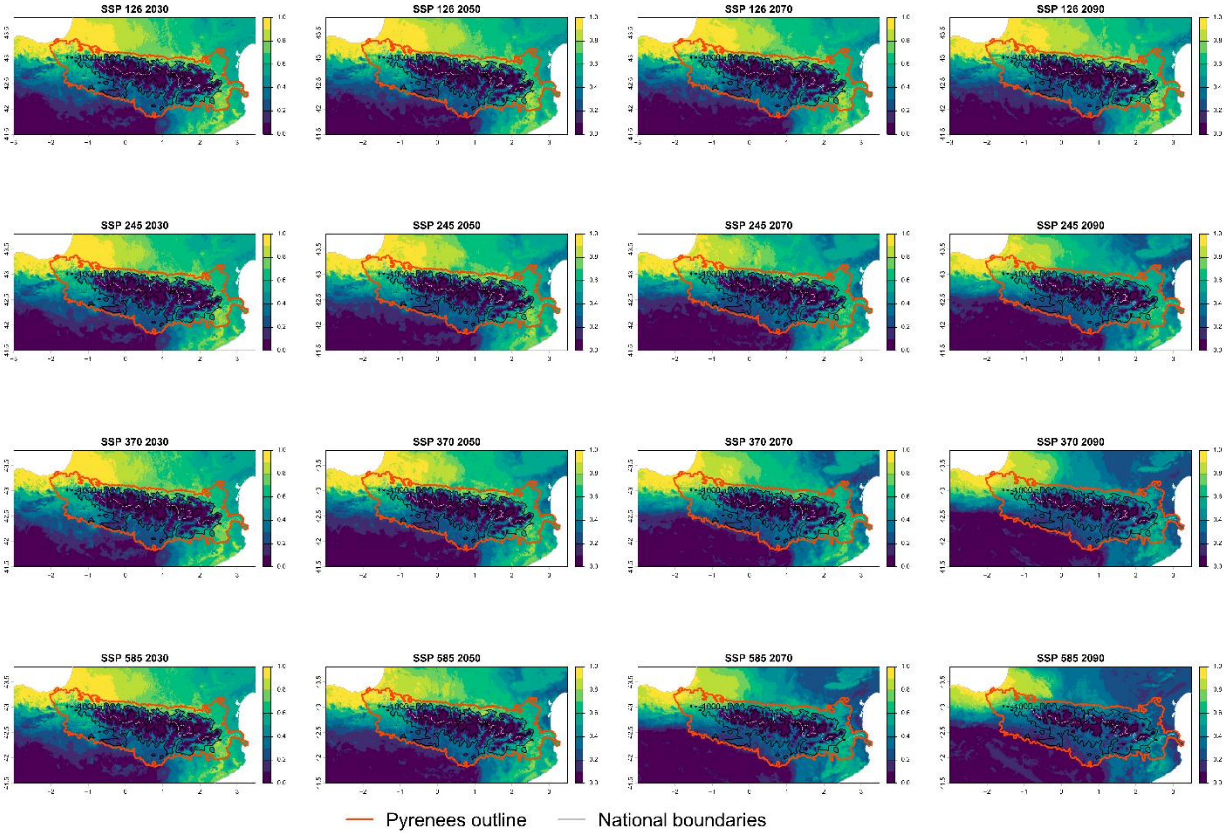

Current bioclimatic suitability in the Pyrenees for *Erigeron sumatrensis*

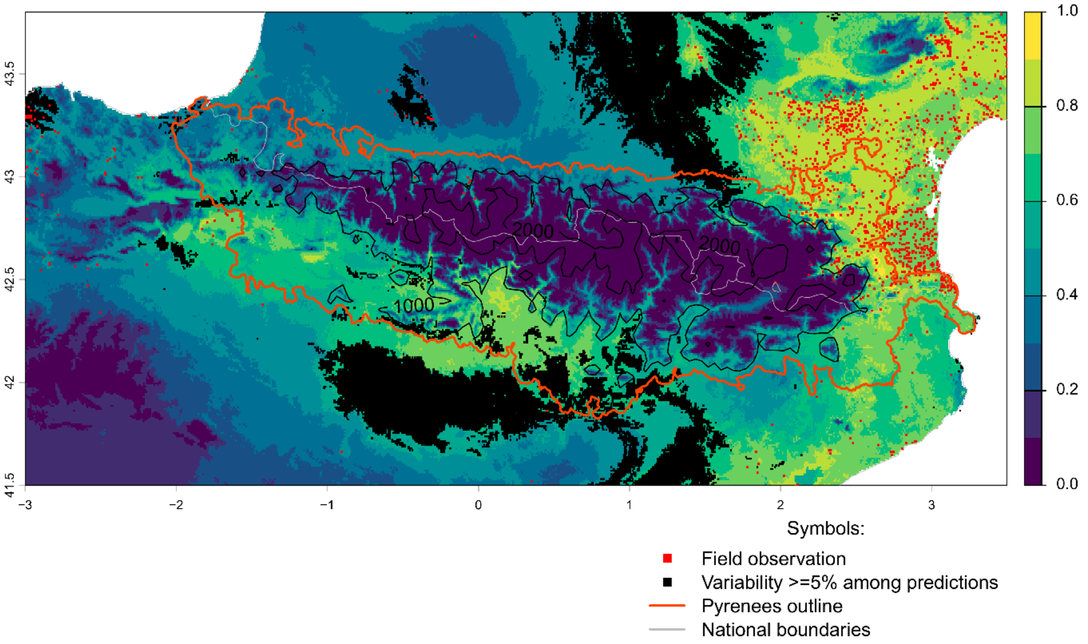


Future bioclimatic suitability in the Pyrenees by 2090 for *Erigeron sumatrensis*

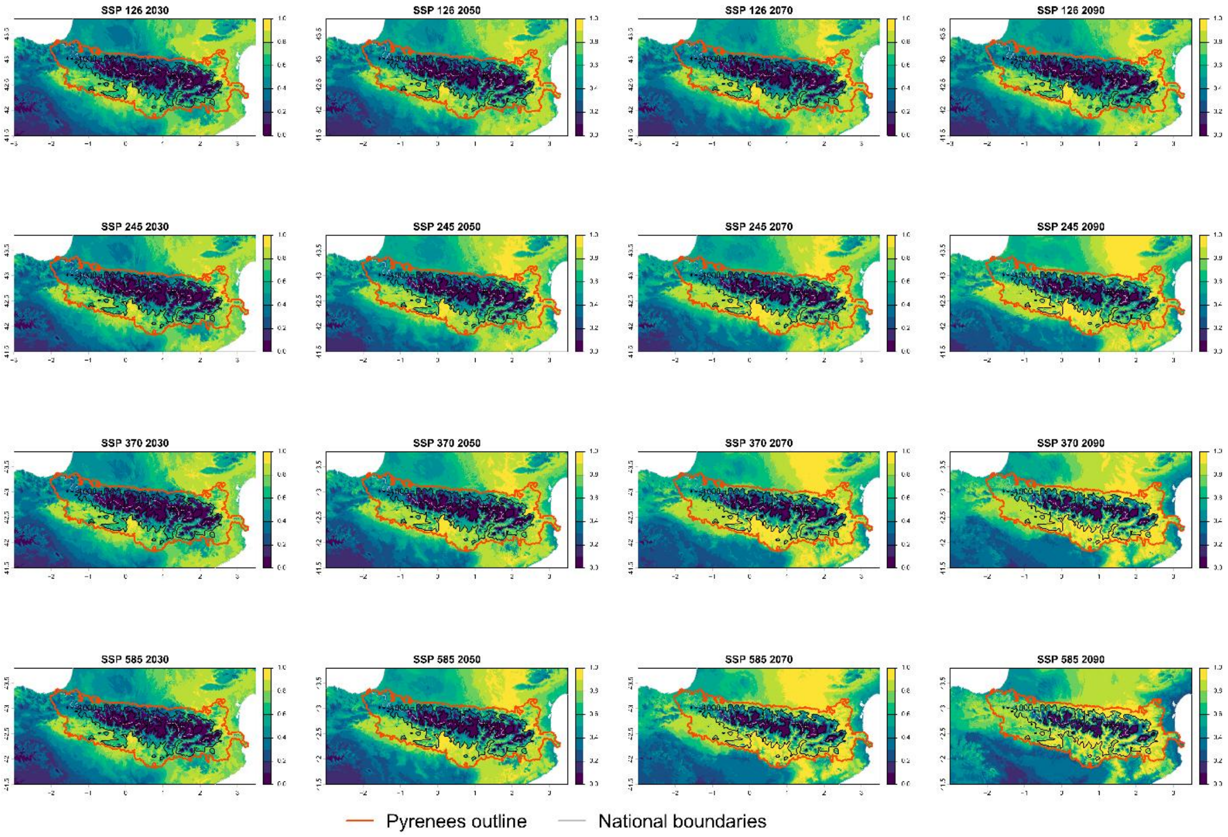

Current bioclimatic suitability in the Pyrenees for *Euphorbia prostrata*

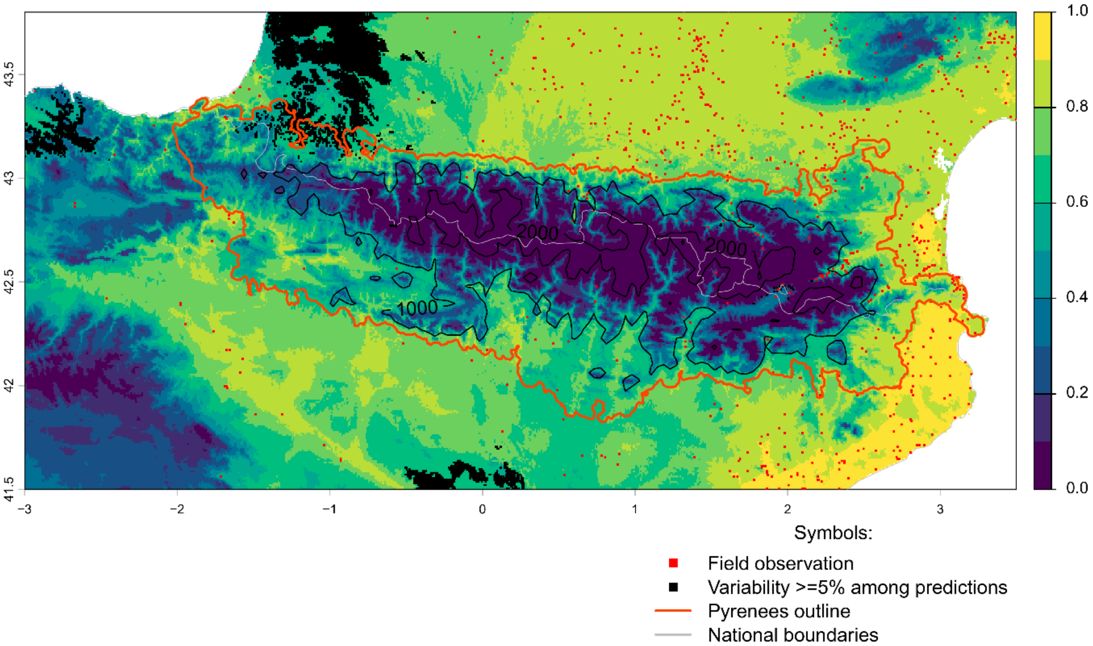


Future bioclimatic suitability in the Pyrenees by 2090 for *Euphorbia prostrata*

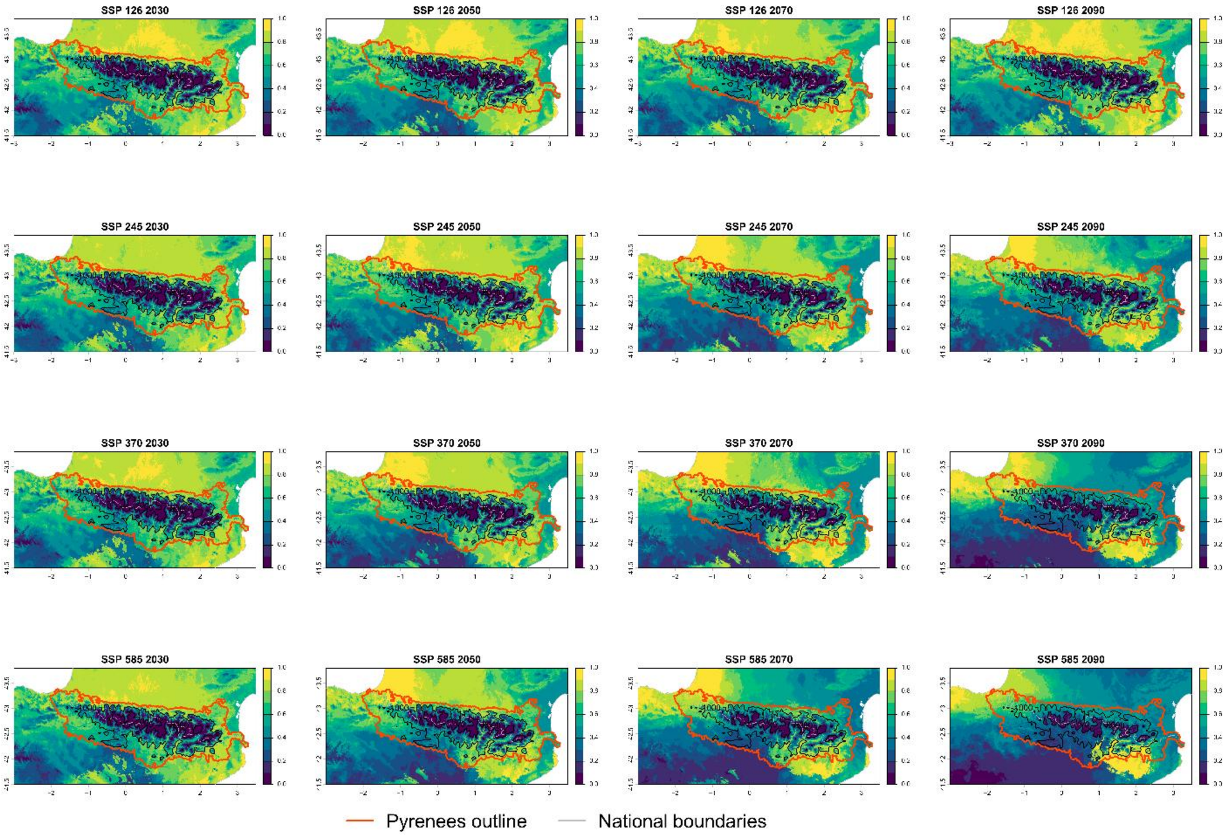

Current bioclimatic suitability in the Pyrenees for *Fallopia baldschuanica*

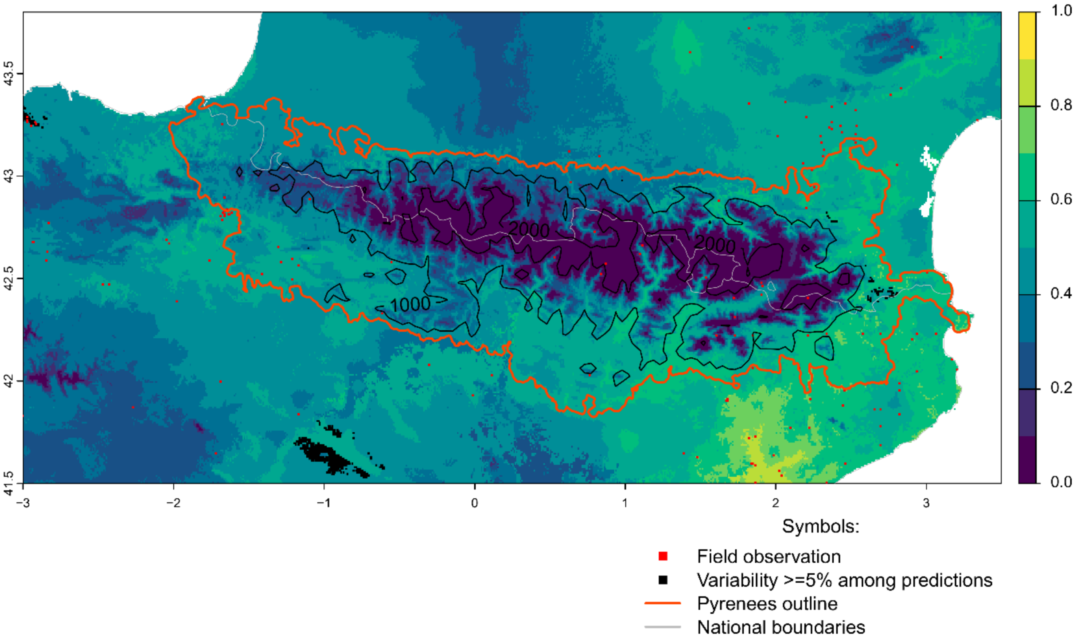


Future bioclimatic suitability in the Pyrenees by 2090 for *Fallopia baldschuanica*

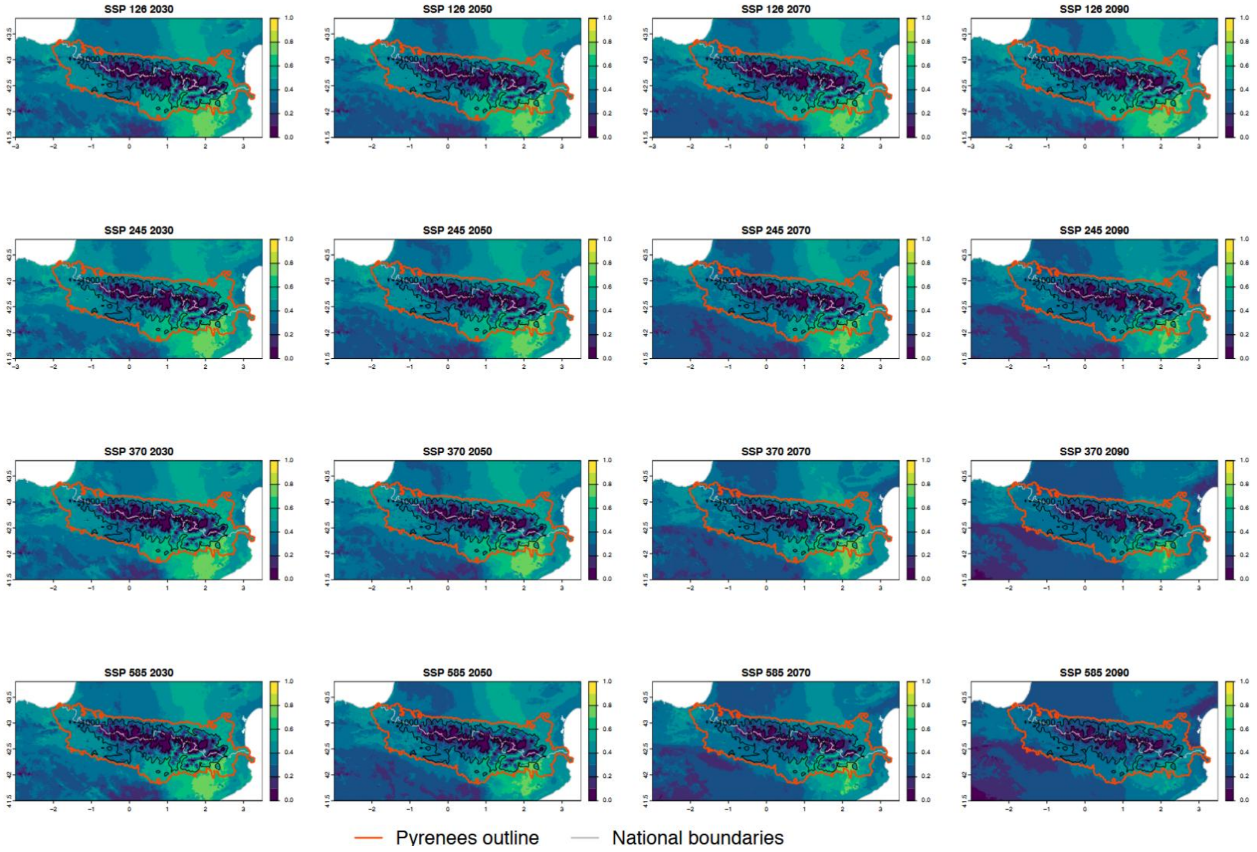

Current bioclimatic suitability in the Pyrenees for *Gleditsia triacanthos*

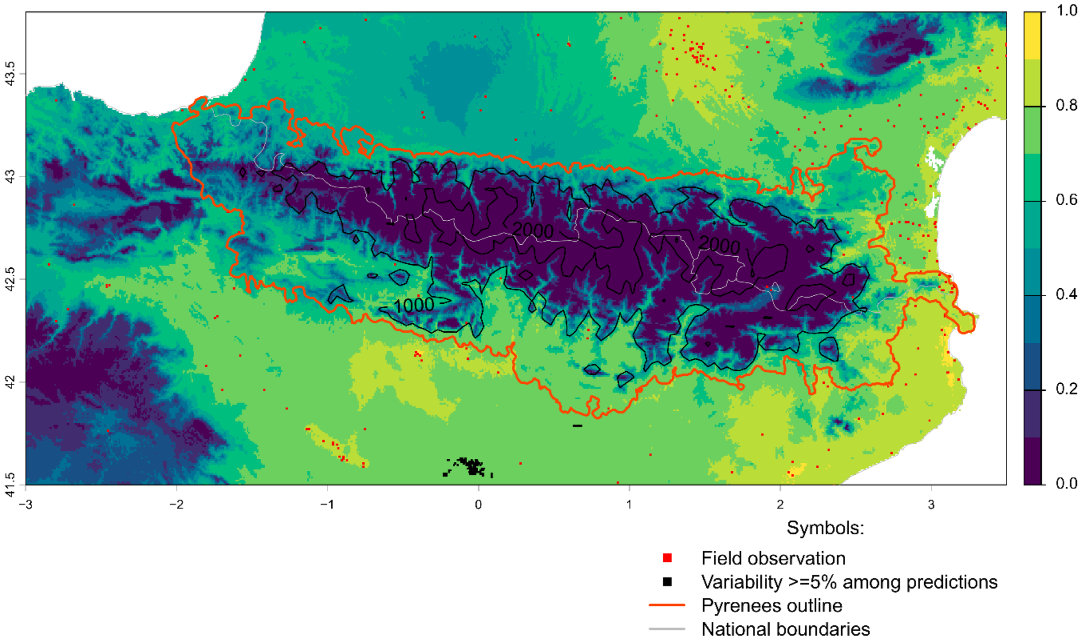


Future bioclimatic suitability in the Pyrenees by 2090 for *Gleditsia triacanthos*

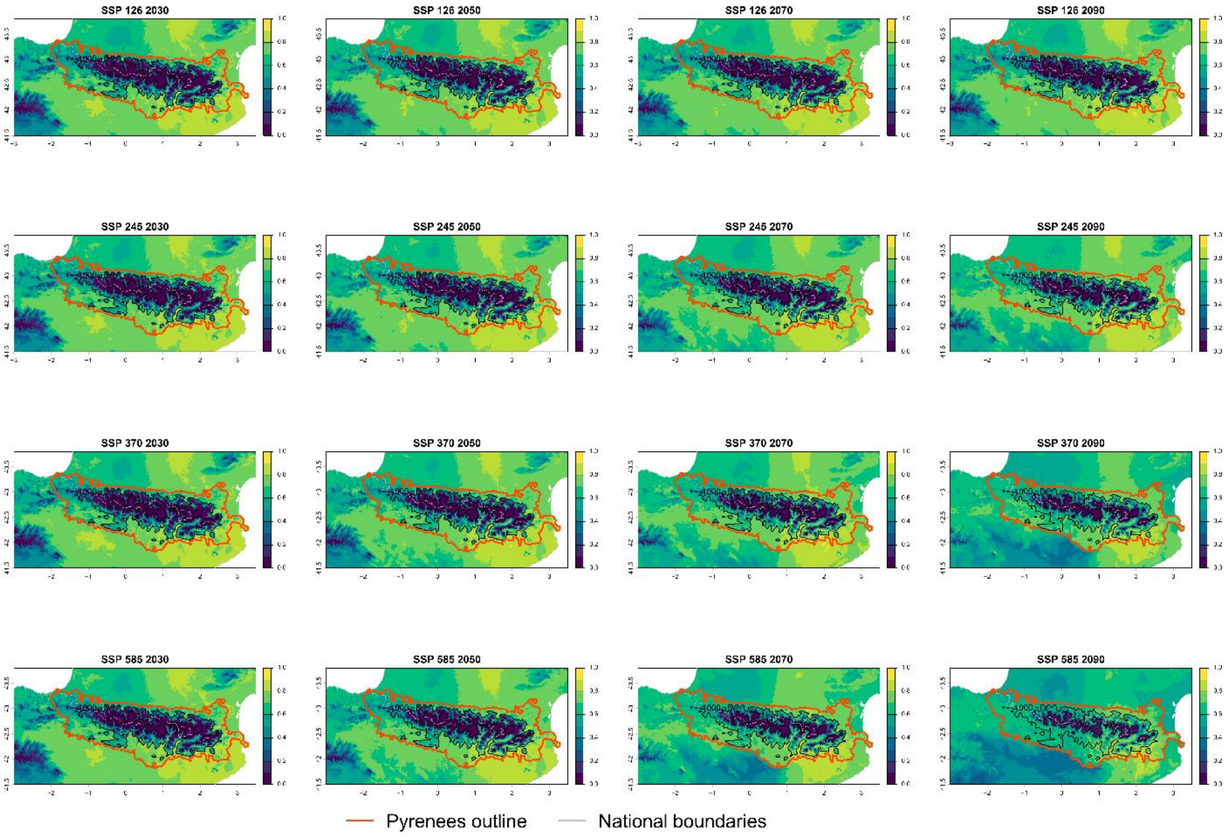

Current bioclimatic suitability in the Pyrenees for *Helianthus tuberosus*

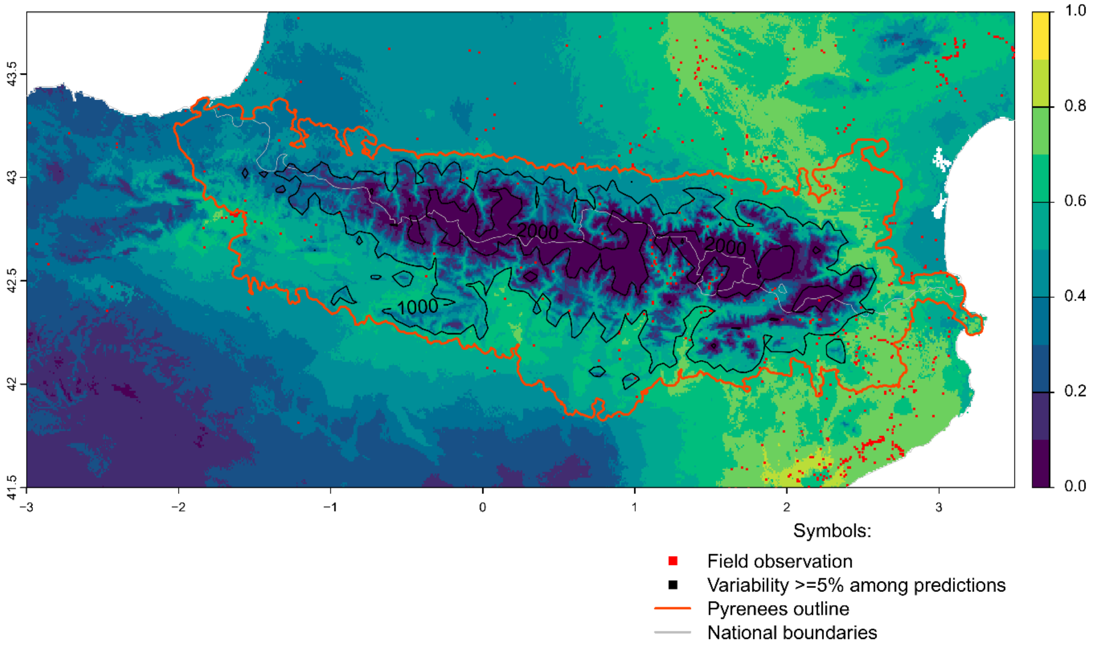


Future bioclimatic suitability in the Pyrenees by 2090 for *Helianthus tuberosus*

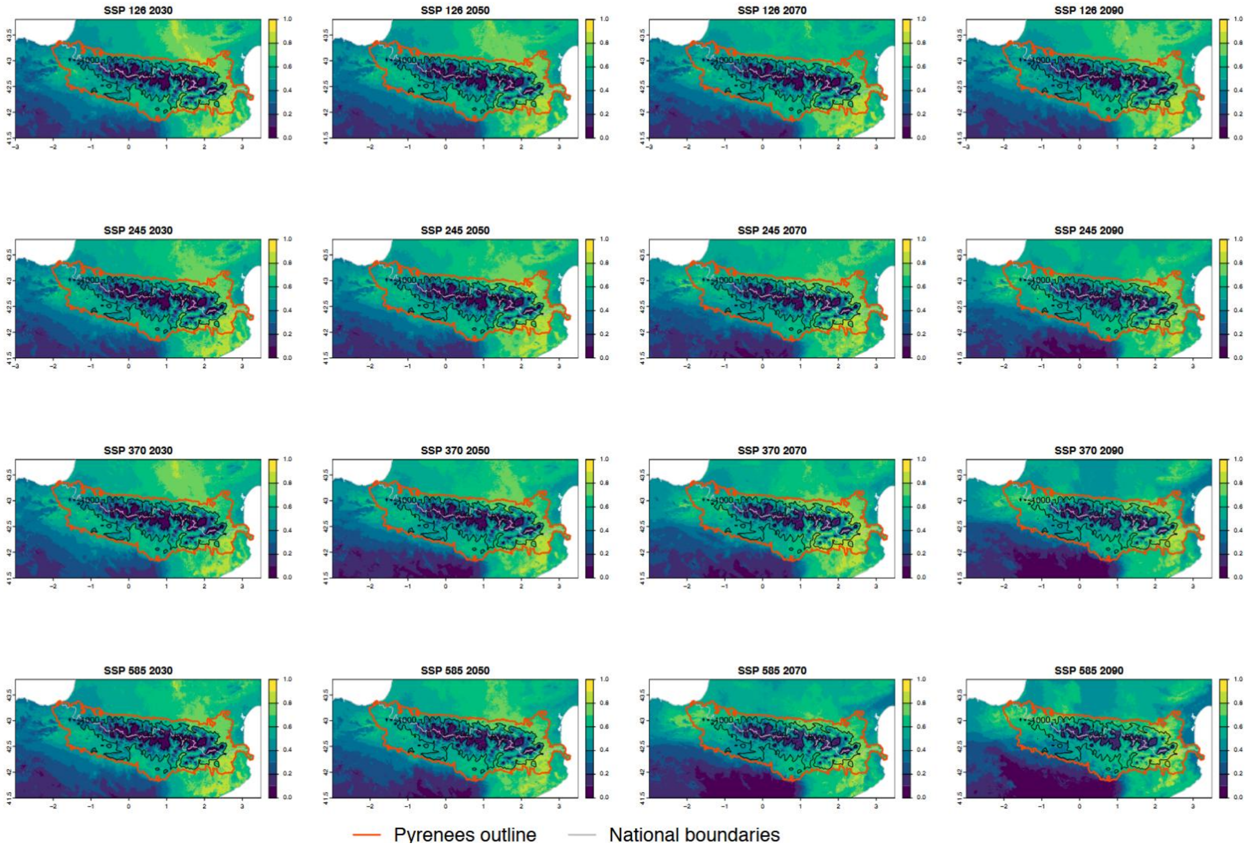

Current bioclimatic suitability in the Pyrenees for *Impatiens balfourii*

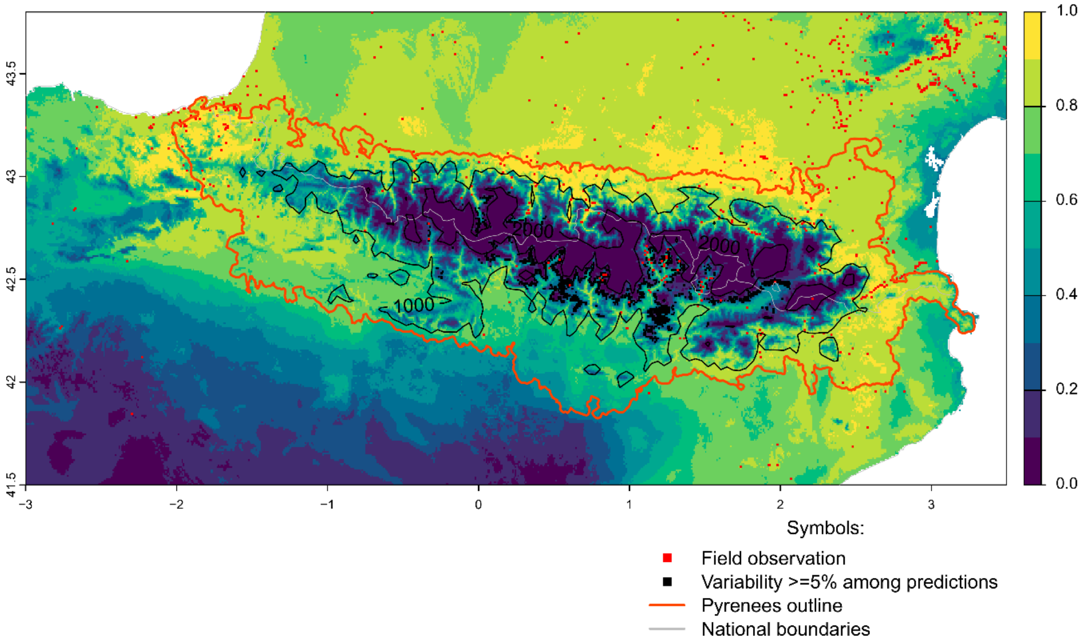


Future bioclimatic suitability in the Pyrenees by 2090 for *Impatiens balfourii*

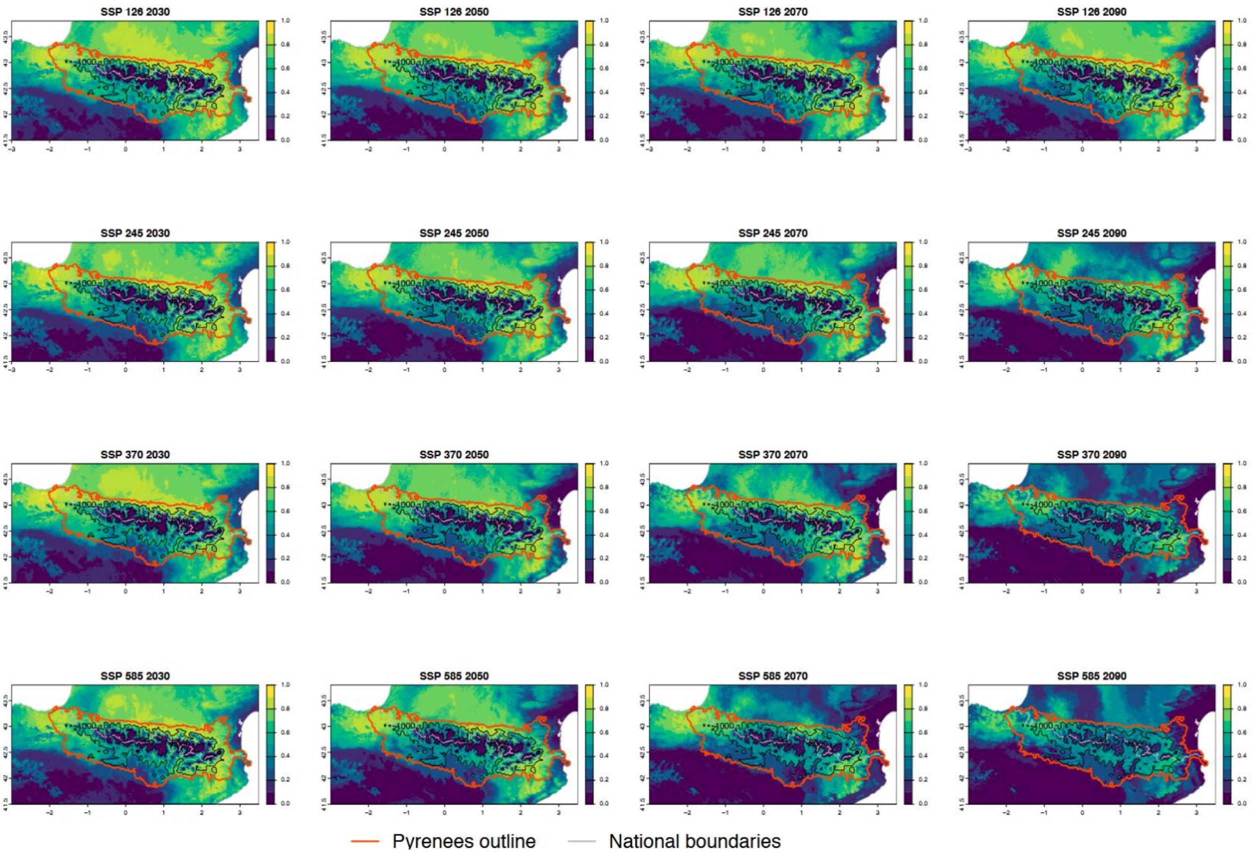

Current bioclimatic suitability in the Pyrenees for *Impatiens glandulifera*

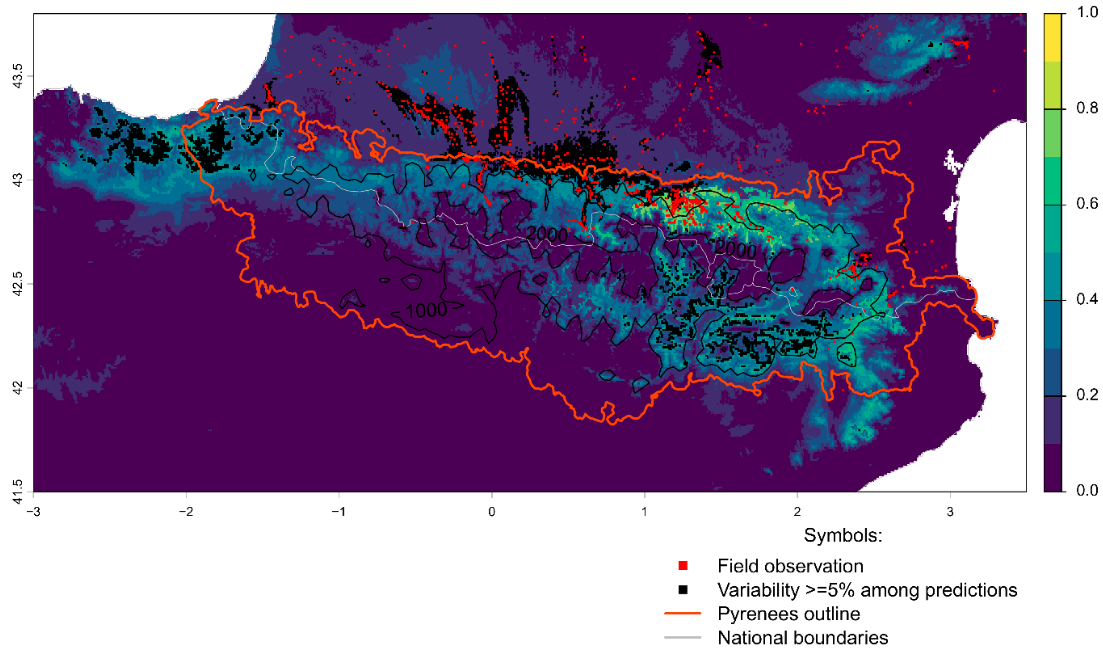


Future bioclimatic suitability in the Pyrenees by 2090 for *Impatiens glandulifera*

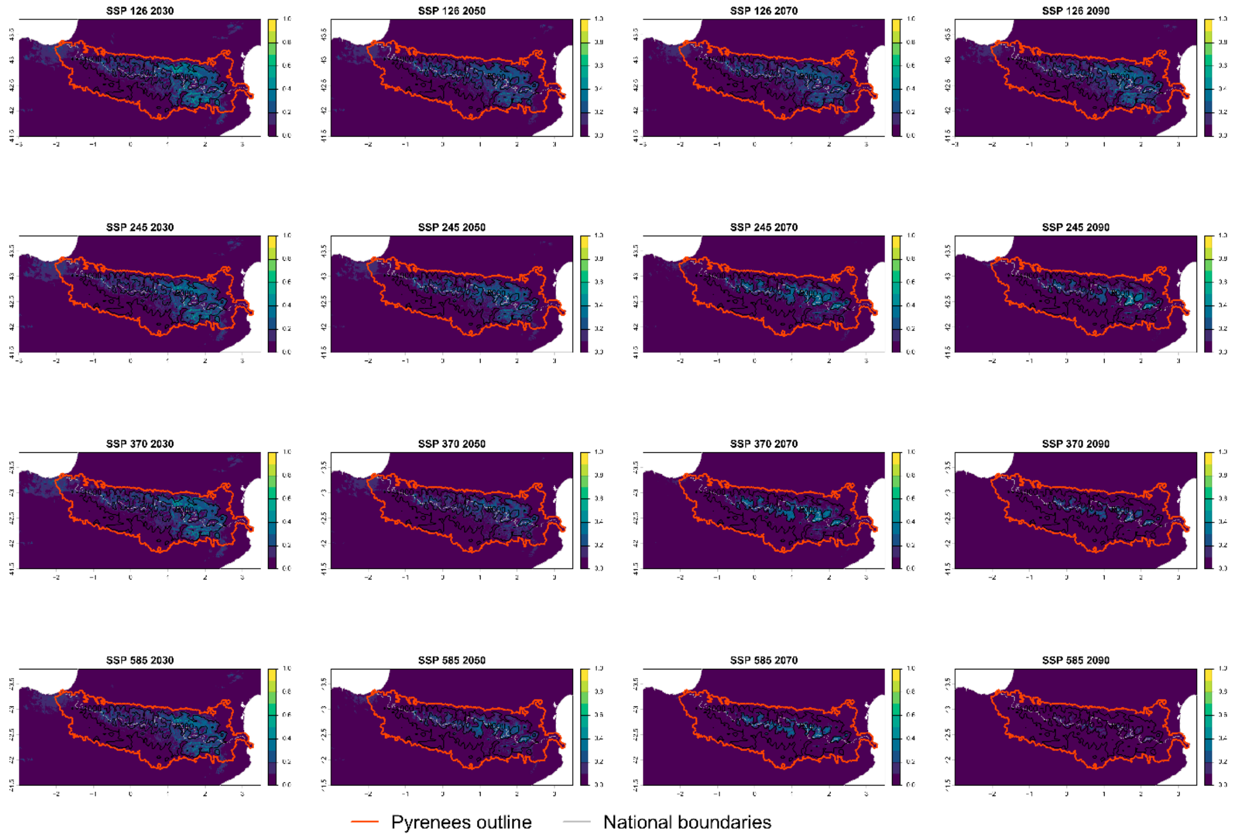

Current bioclimatic suitability in the Pyrenees for *Ligustrum lucidum*

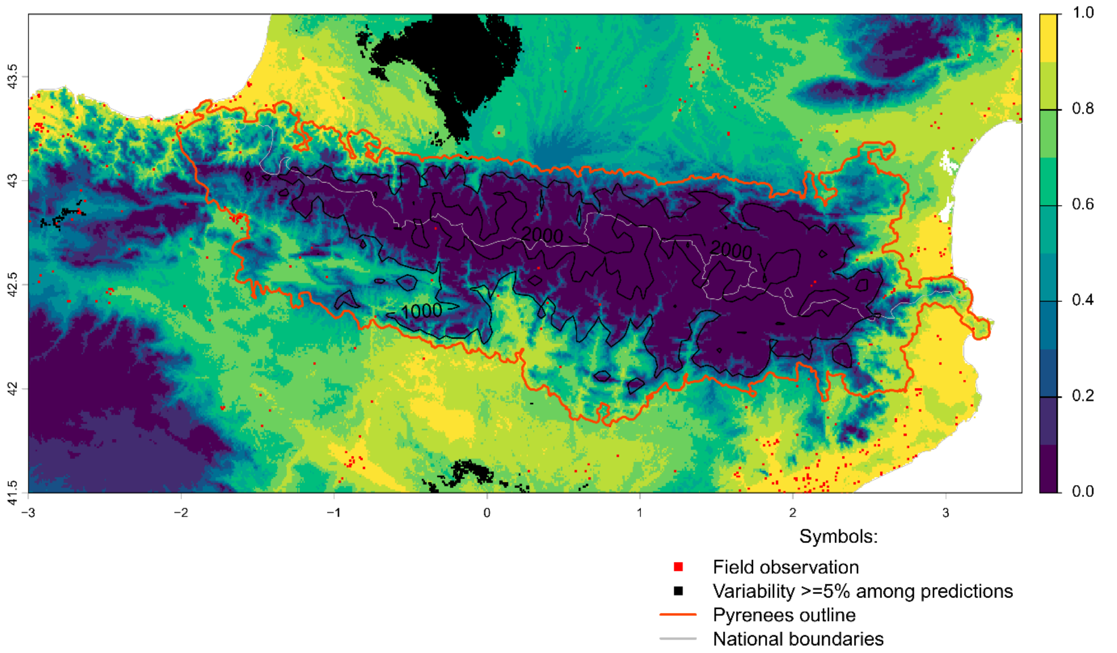


Future bioclimatic suitability in the Pyrenees by 2090 for *Ligustrum lucidum*

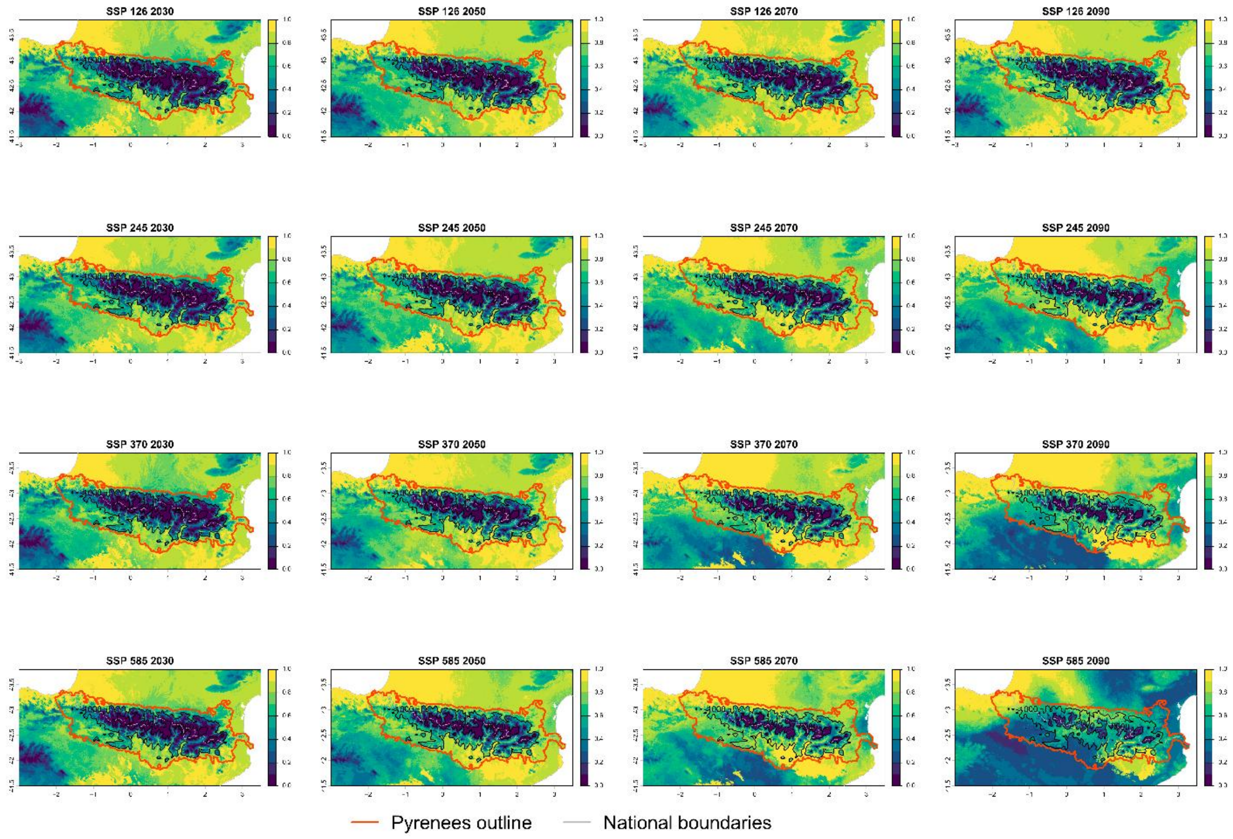

Current bioclimatic suitability in the Pyrenees for *Lonicera japonica*

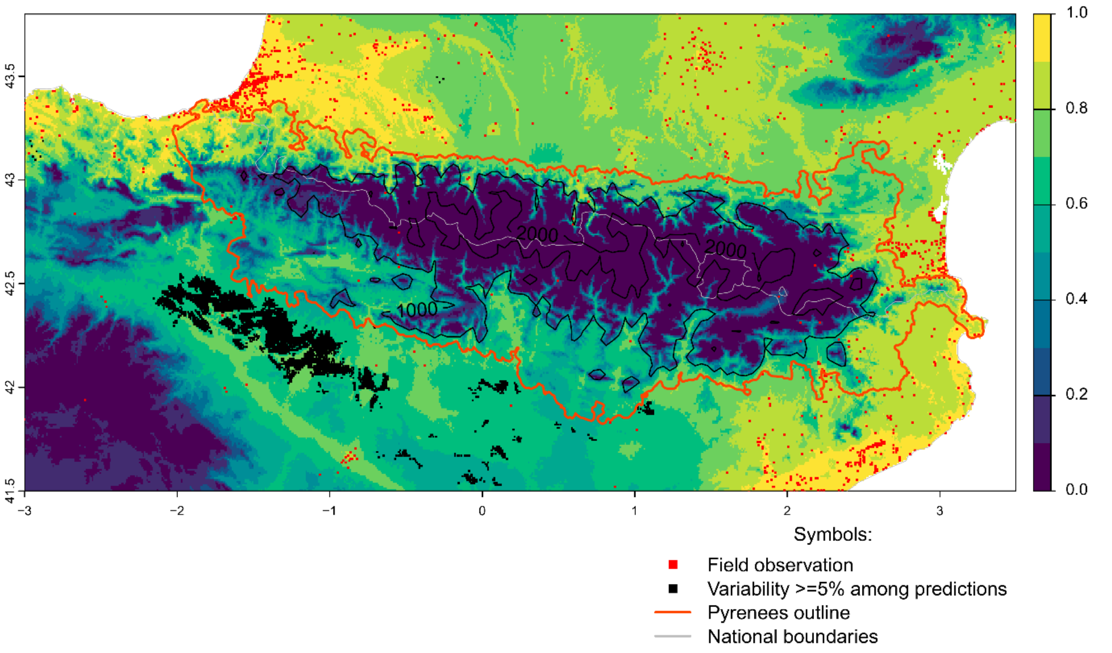


Future bioclimatic suitability in the Pyrenees by 2090 for *Lonicera japonica*

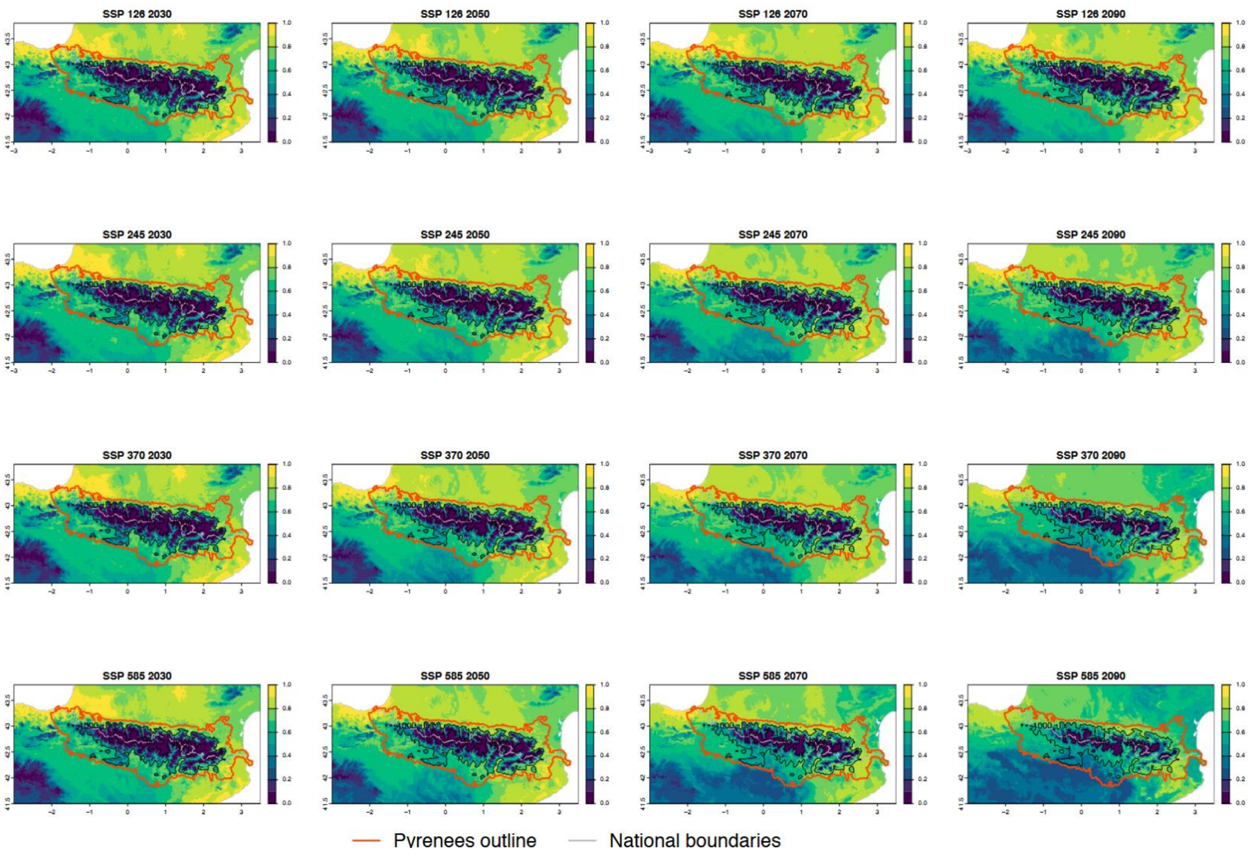

Current bioclimatic suitability in the Pyrenees for *Ludwigia peploides*

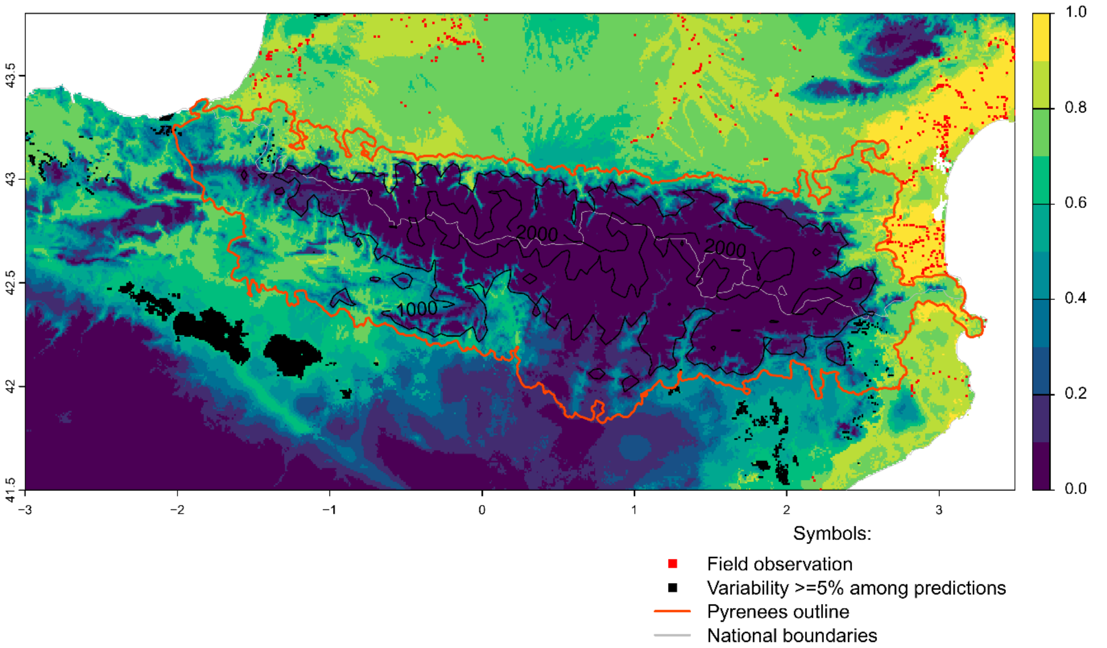


Future bioclimatic suitability in the Pyrenees by 2090 for *Ludwigia peploides*

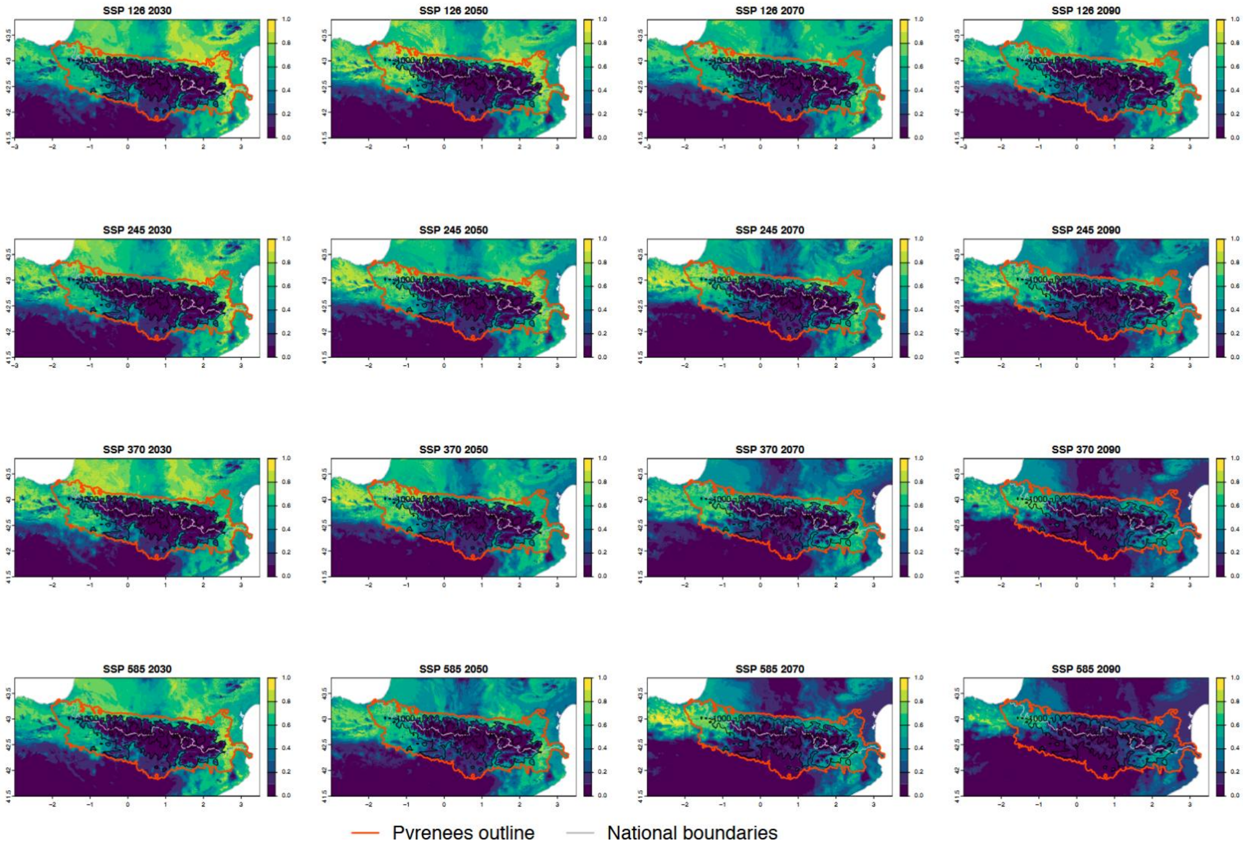

Current bioclimatic suitability in the Pyrenees for *Opuntia stricta*

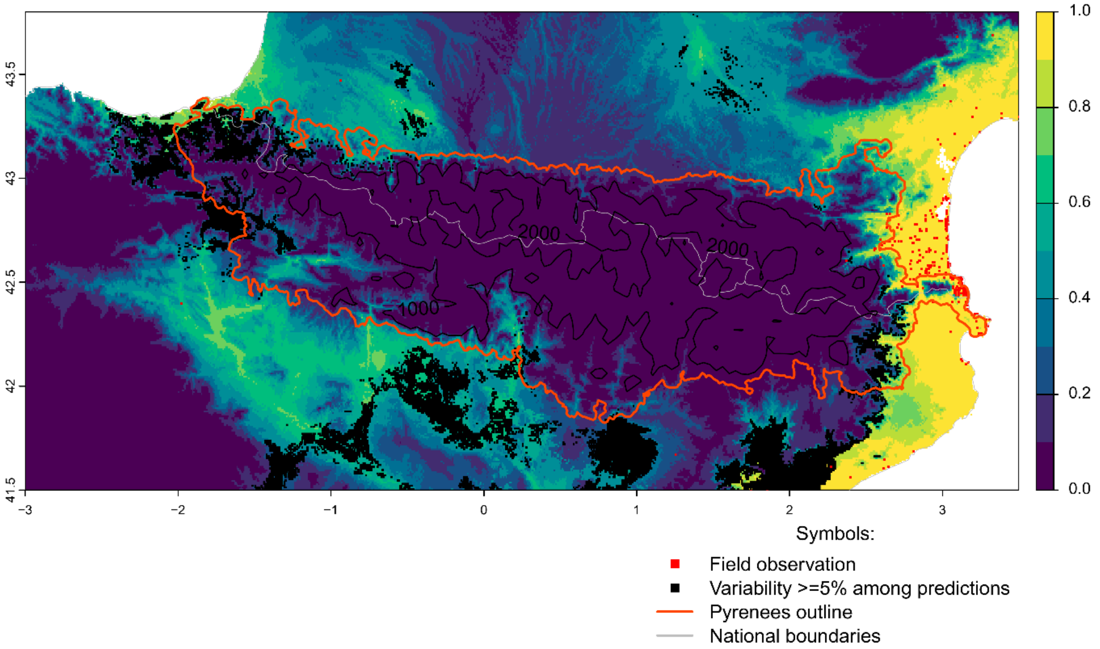


Future bioclimatic suitability in the Pyrenees by 2090 for *Opuntia stricta*

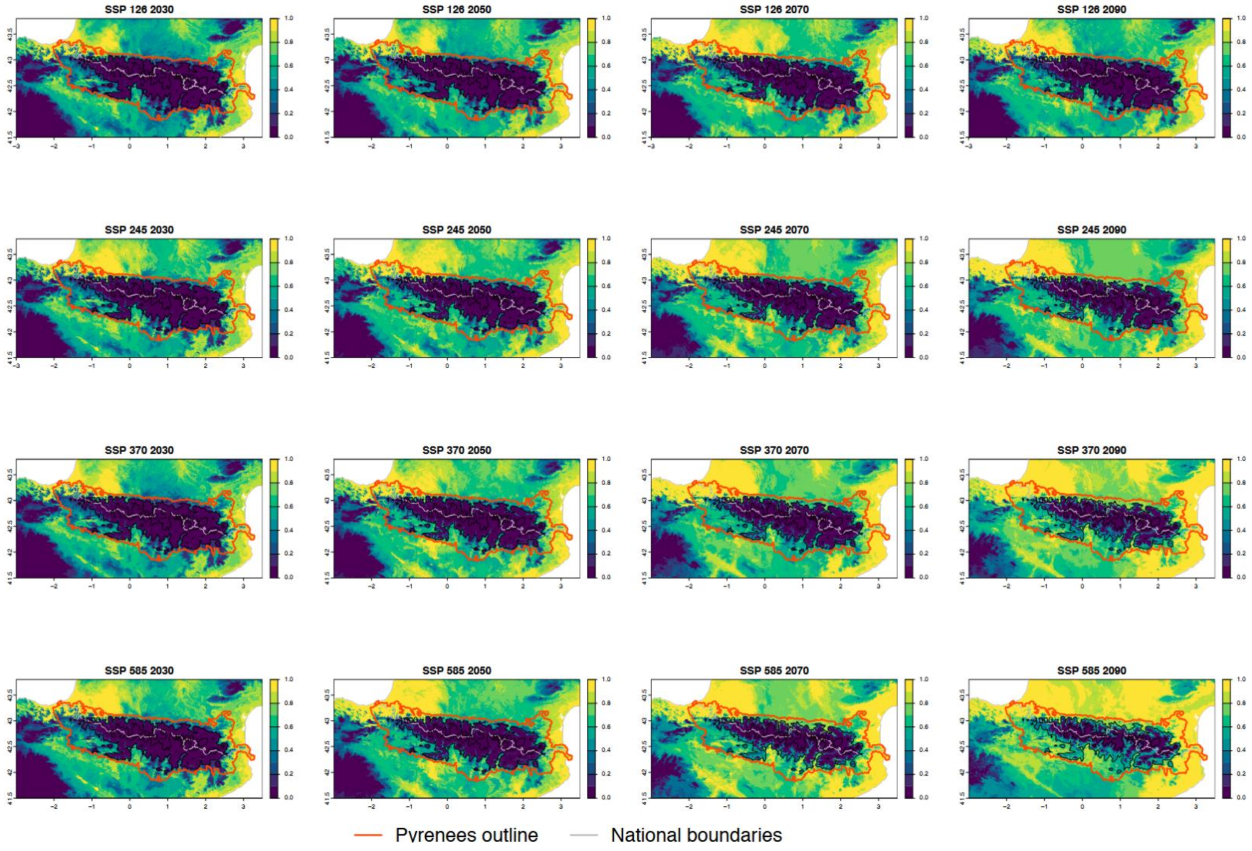

Current bioclimatic suitability in the Pyrenees for *Parthenocissus inserta*

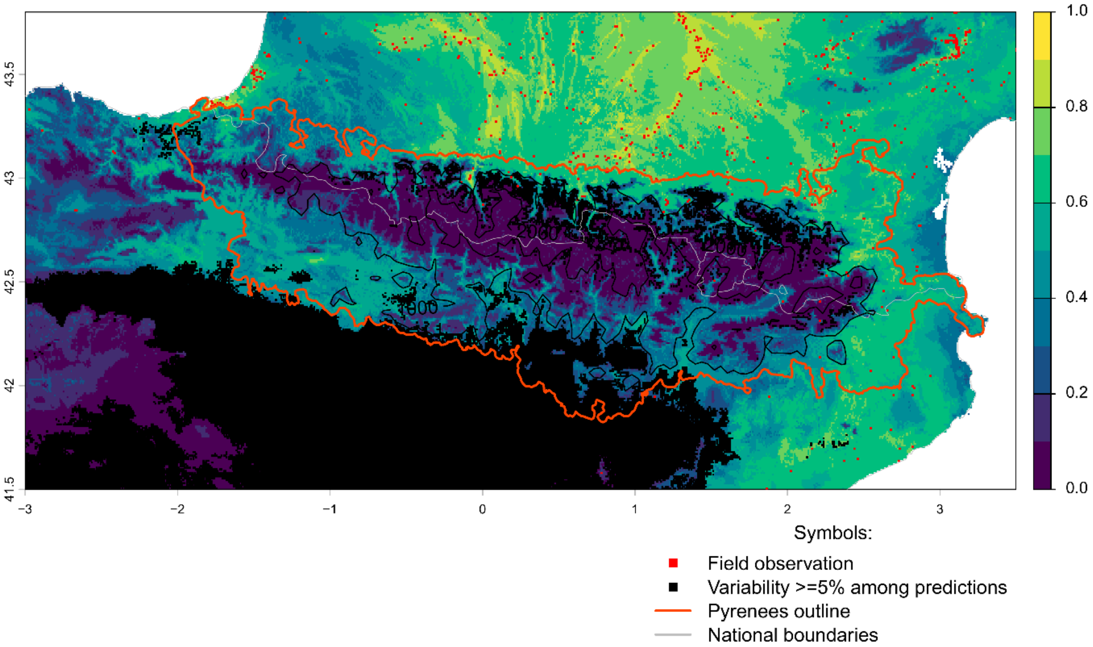


Future bioclimatic suitability in the Pyrenees by 2090 for *Parthenocissus inserta*

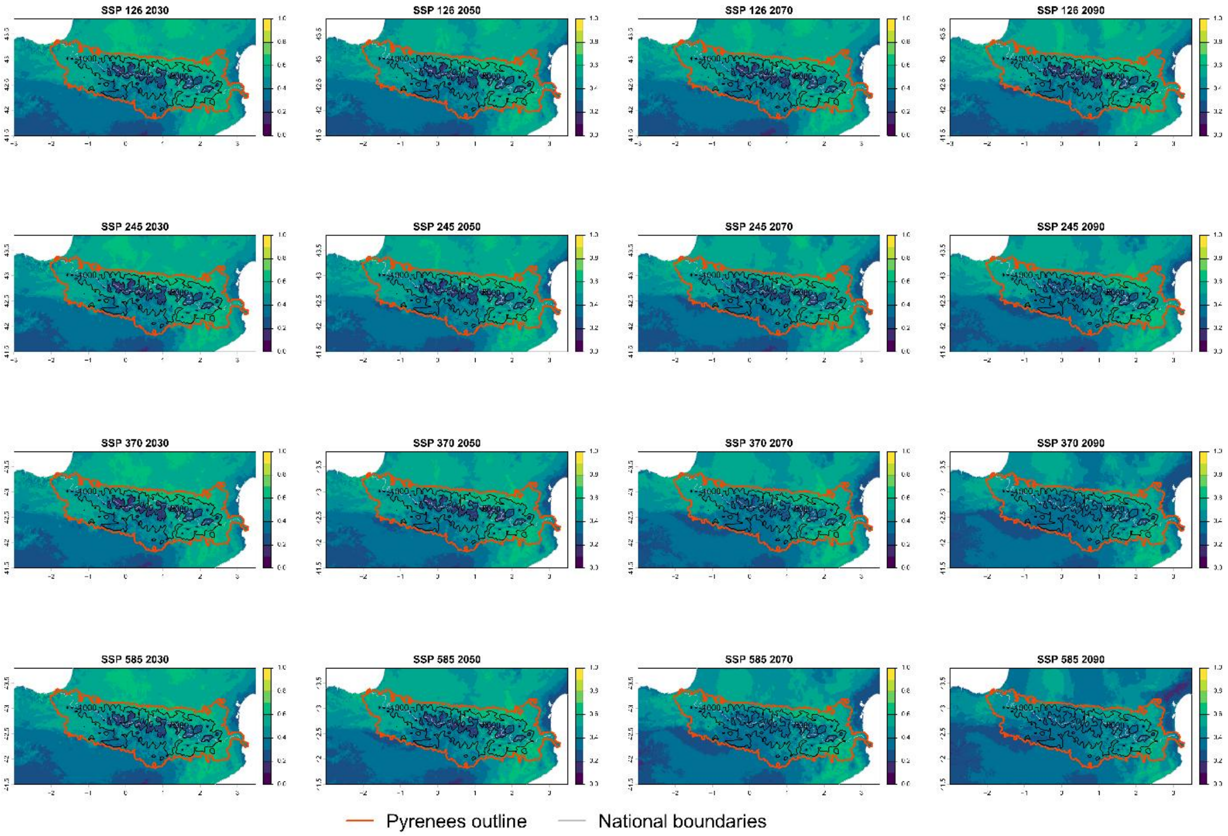

Current bioclimatic suitability in the Pyrenees for *Paspalum dilatatum*

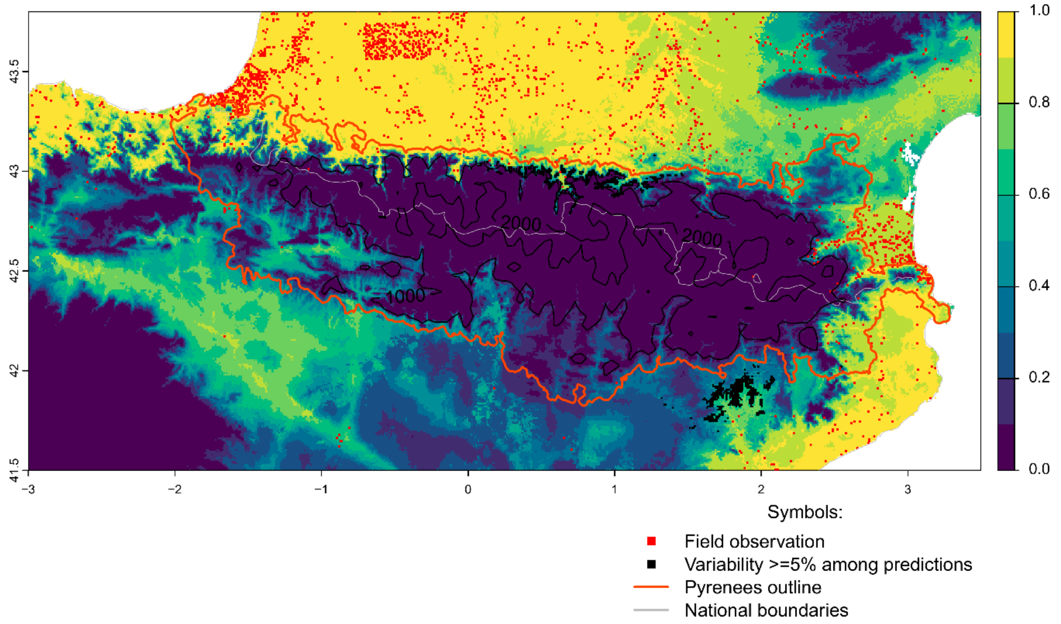


Future bioclimatic suitability in the Pyrenees by 2090 for *Paspalum dilatatum*

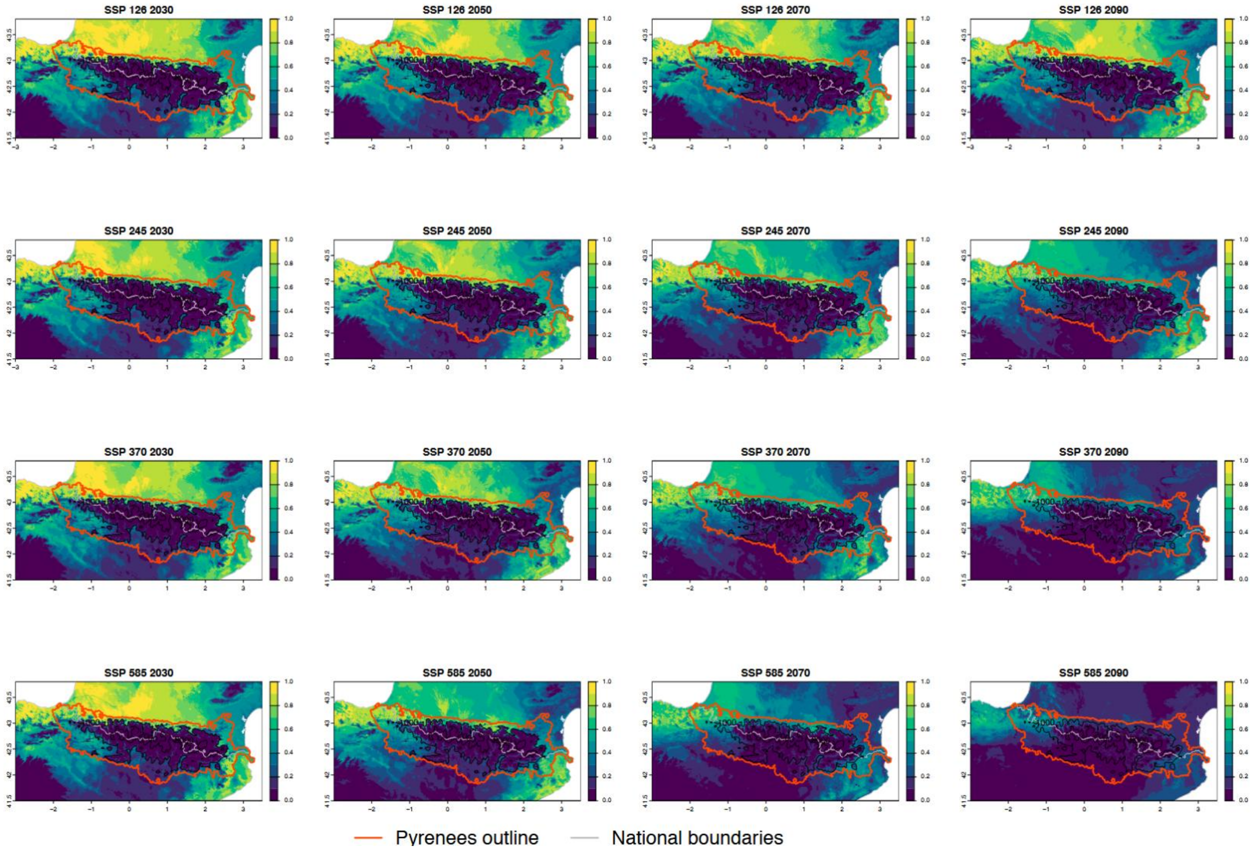

Current bioclimatic suitability in the Pyrenees for *Periploca graeca*

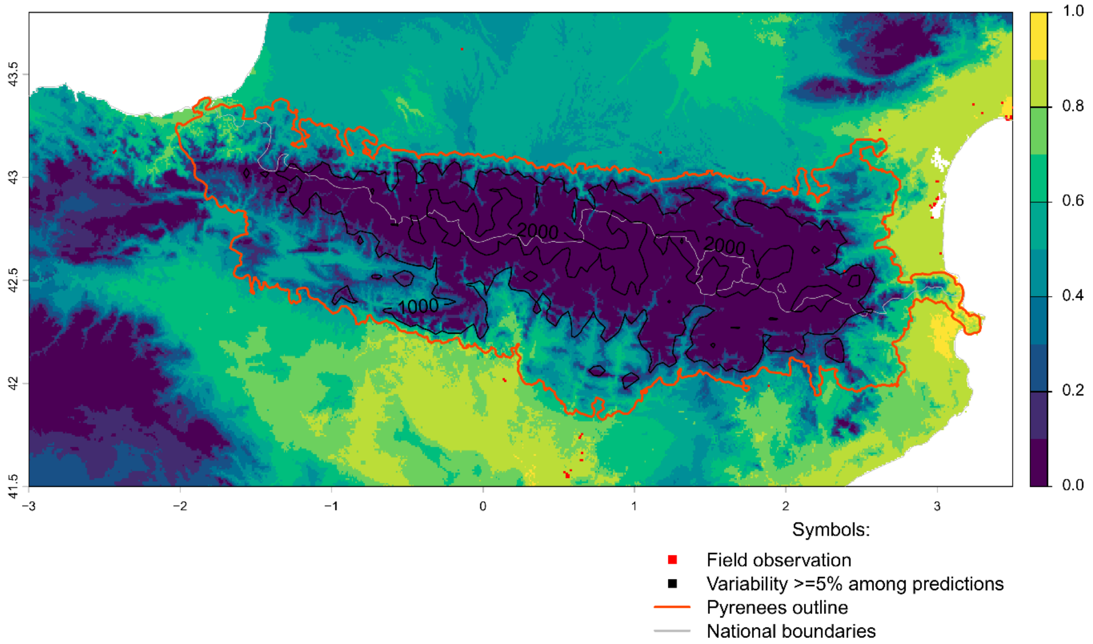


Future bioclimatic suitability in the Pyrenees by 2090 for *Periploca graeca*

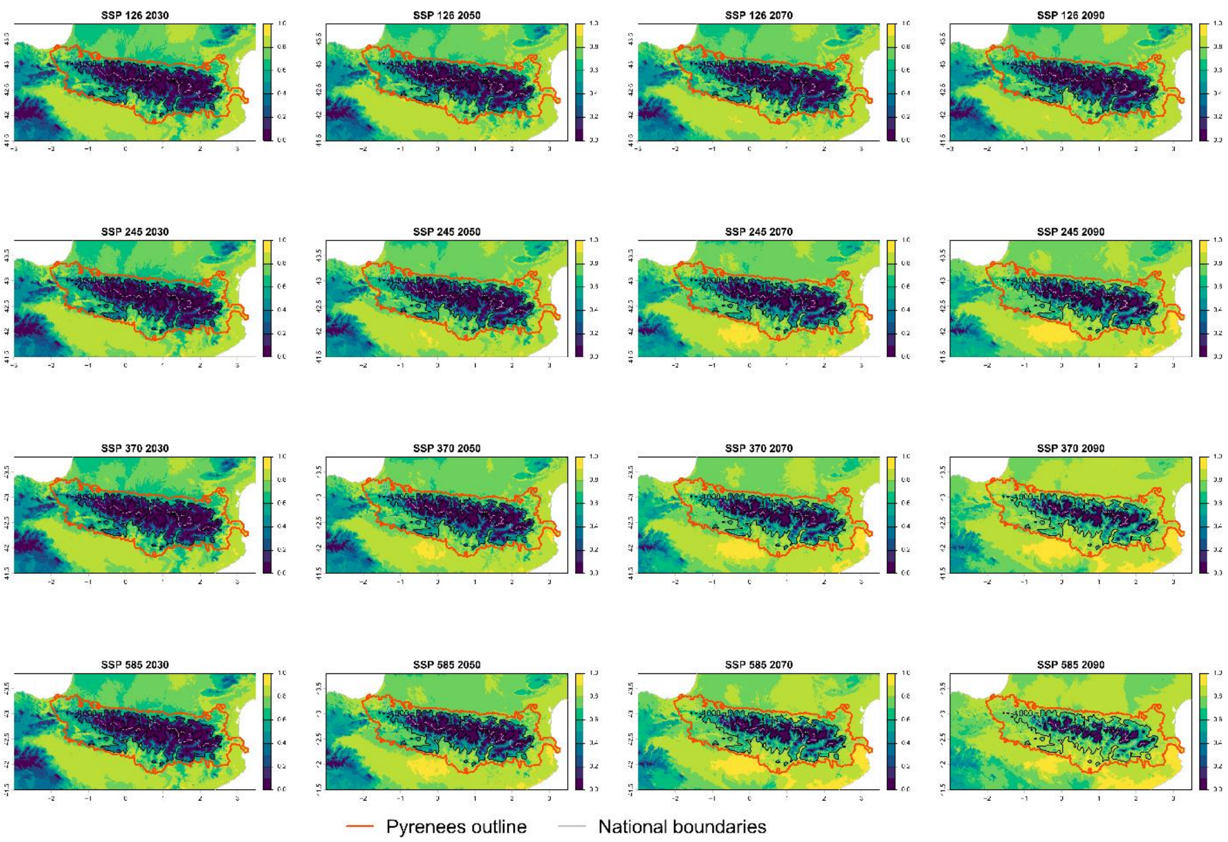

Current bioclimatic suitability in the Pyrenees for *Phyllostachys aurea*

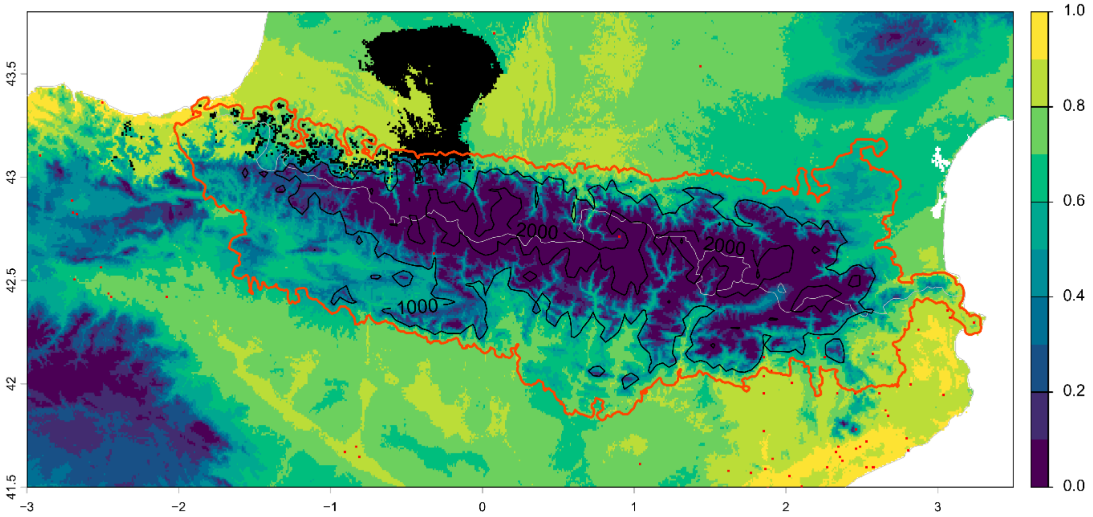


Symbols:

- Field observation
- Variability >=5% among predictions
- Pyrenees outline
- National boundaries

Future bioclimatic suitability in the Pyrenees by 2090 for *Phyllostachys aurea*

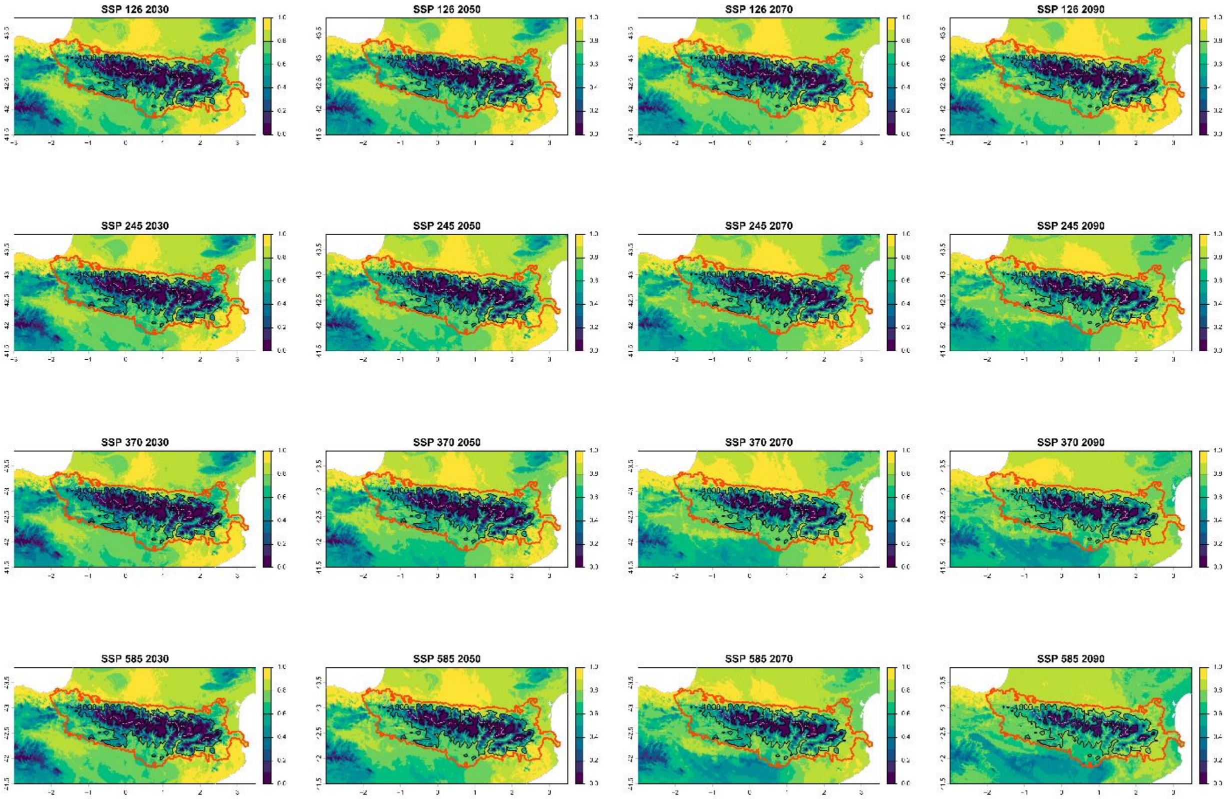


Pyrenees outline National boundaries

Current bioclimatic suitability in the Pyrenees for *Phyllostachys bambusoides*

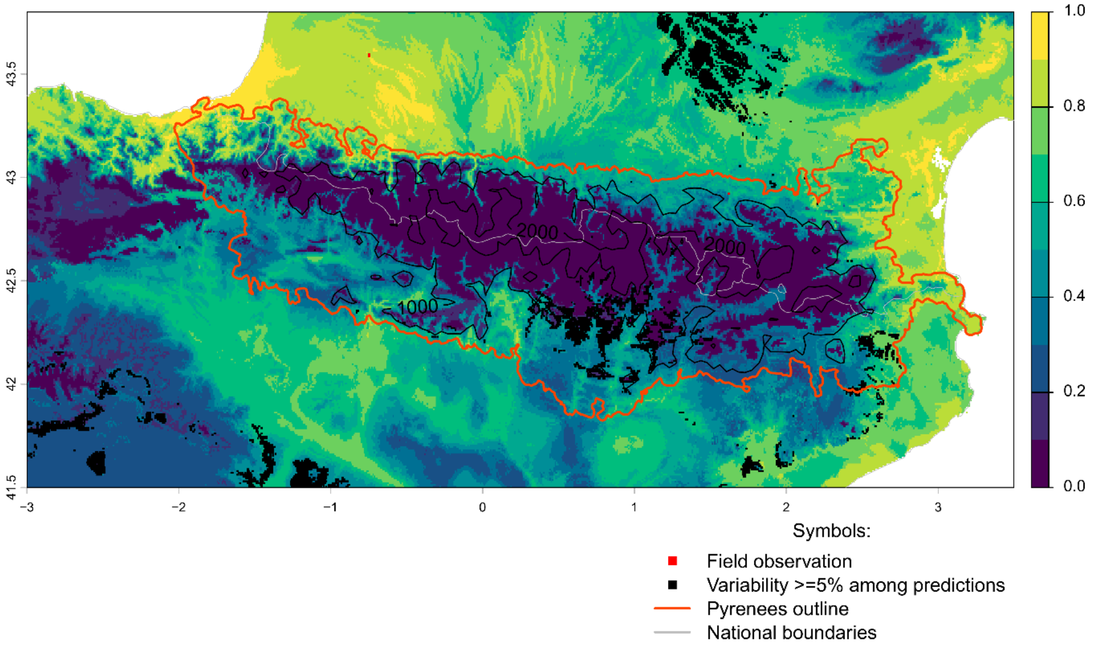


Future bioclimatic suitability in the Pyrenees by 2090 for *Phyllostachys bambusoides*

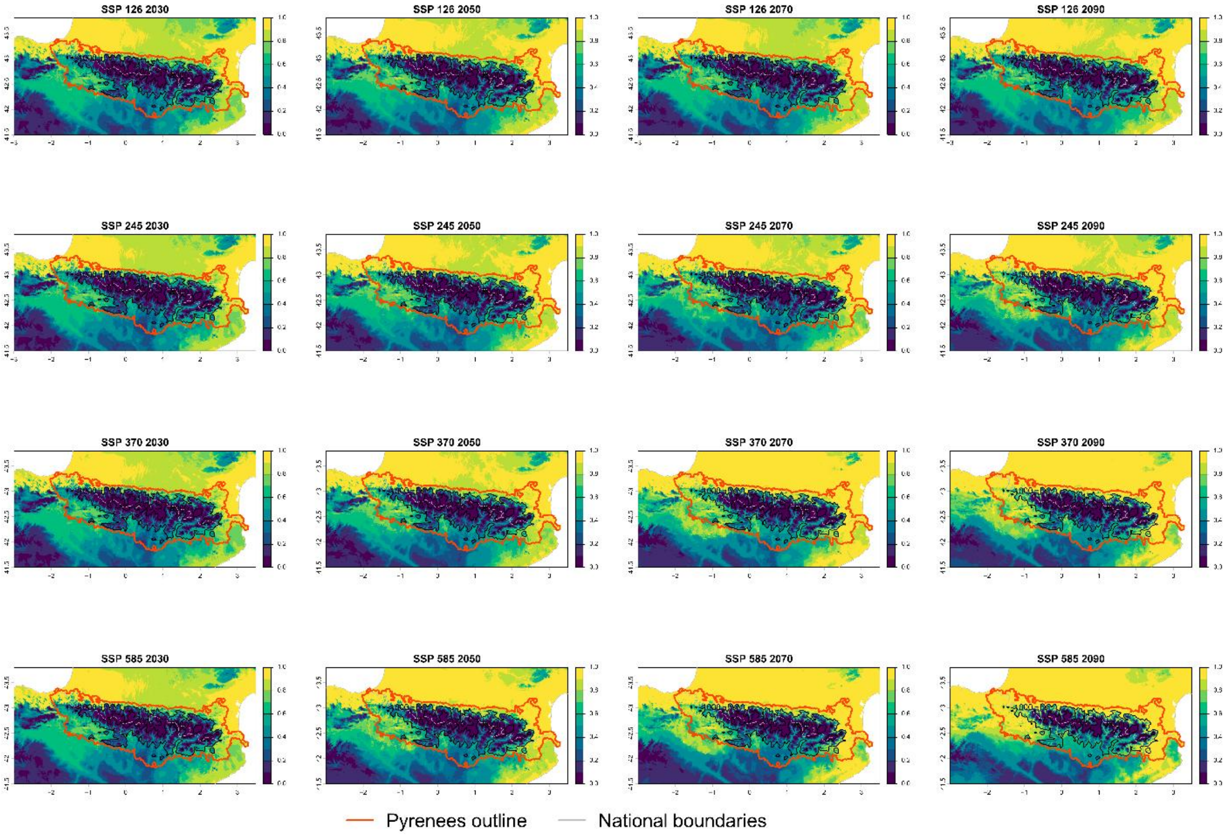

Current bioclimatic suitability in the Pyrenees for *Phyllostachys nigra*

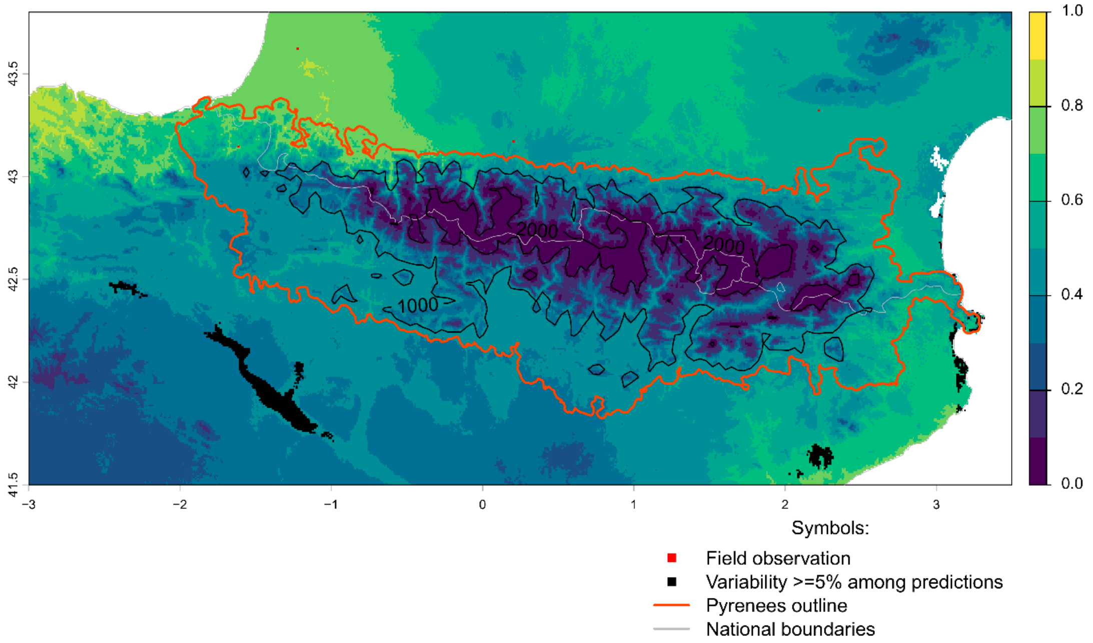


Future bioclimatic suitability in the Pyrenees by 2090 for *Phyllostachys nigra*

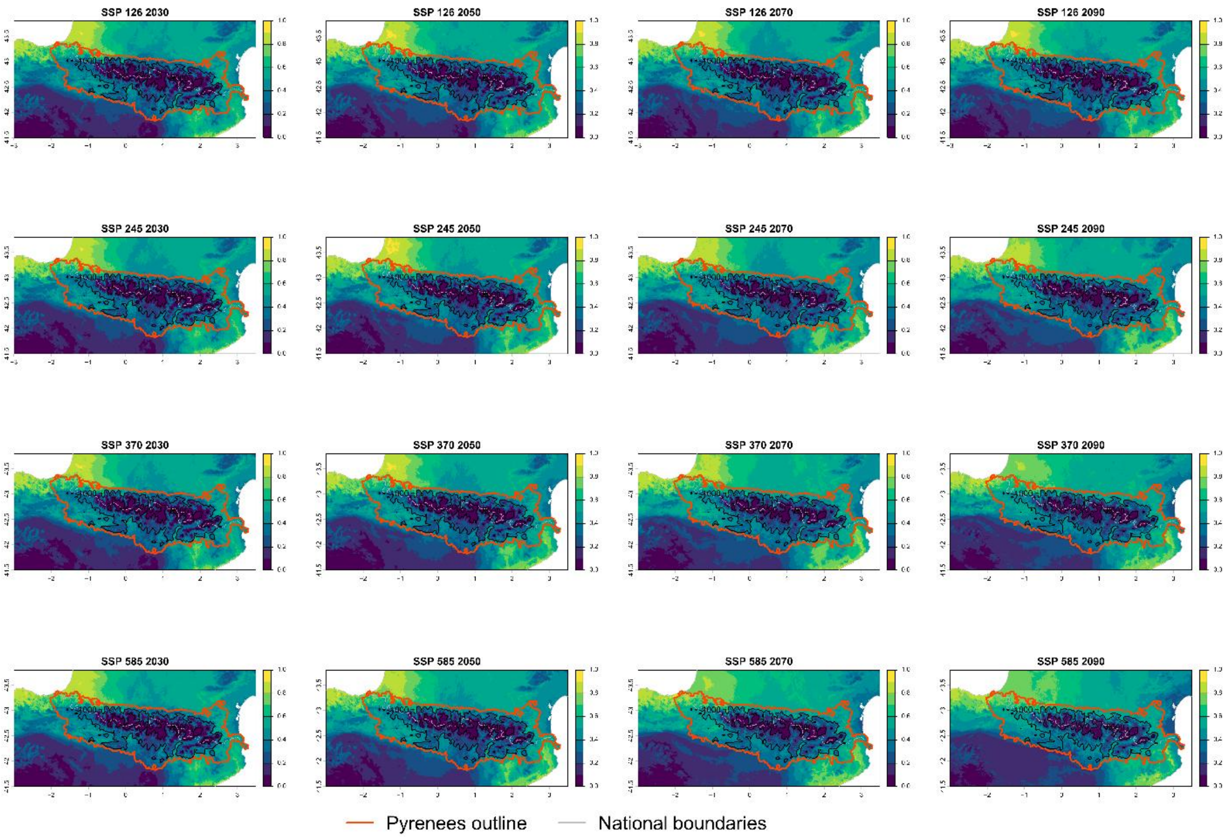

Current bioclimatic suitability in the Pyrenees for *Phytolacca americana*

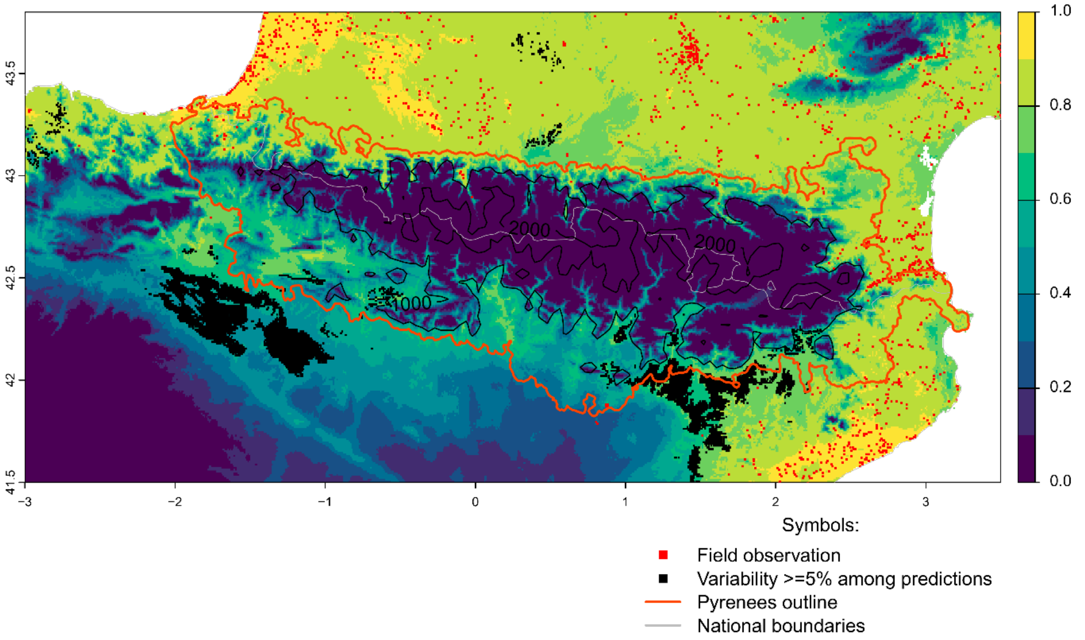


Future bioclimatic suitability in the Pyrenees by 2090 for *Phytolacca americana*

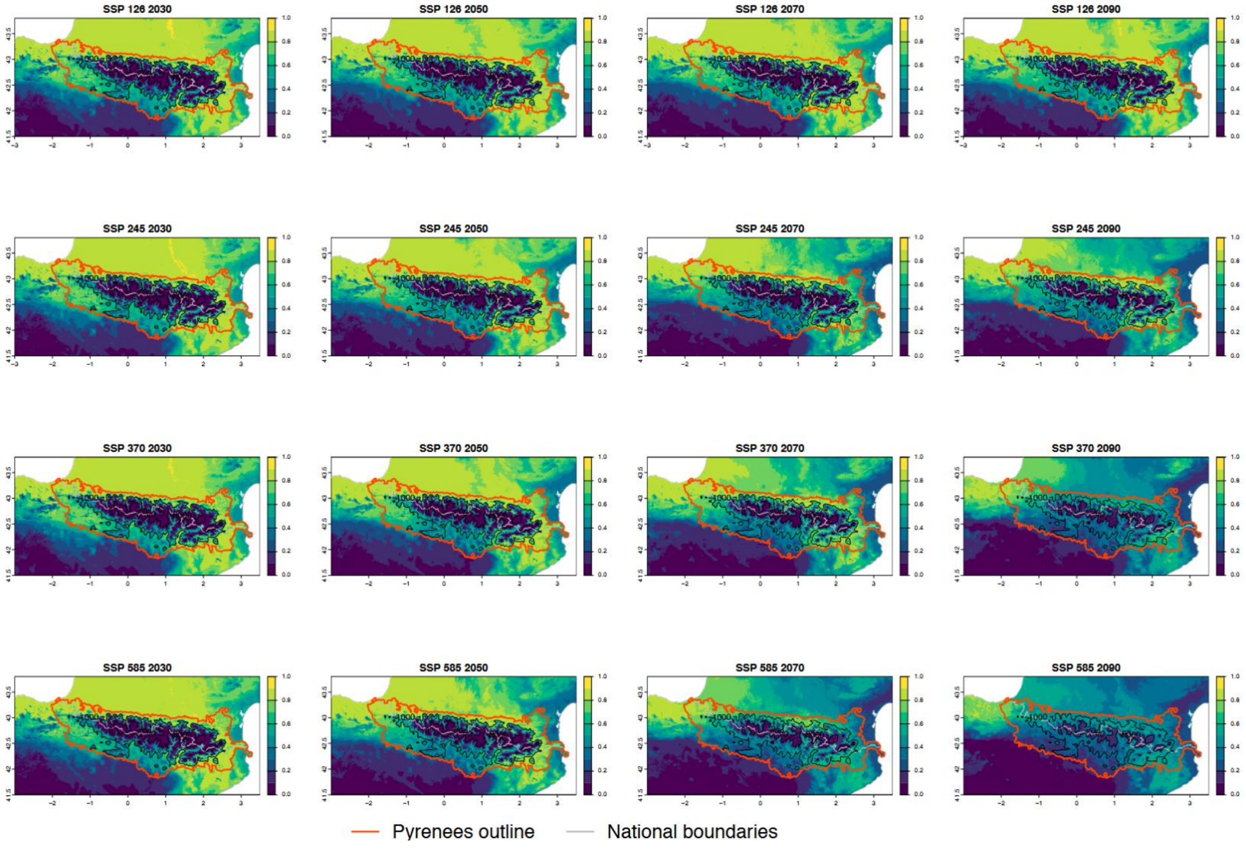

Current bioclimatic suitability in the Pyrenees for *Prunus cerasifera*

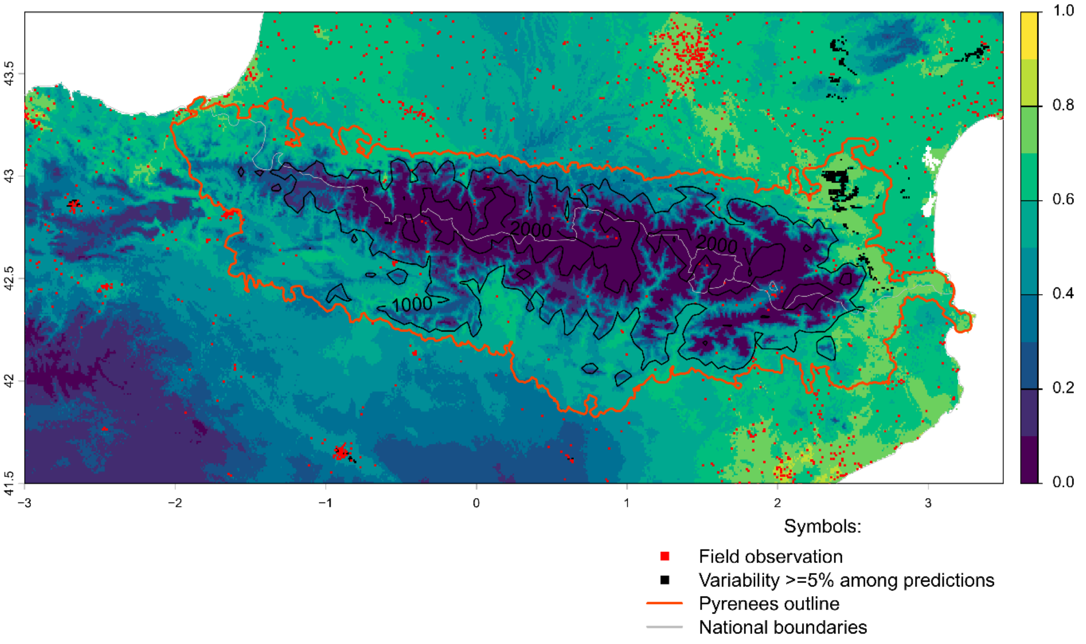


Future bioclimatic suitability in the Pyrenees by 2090 for *Prunus cerasifera*

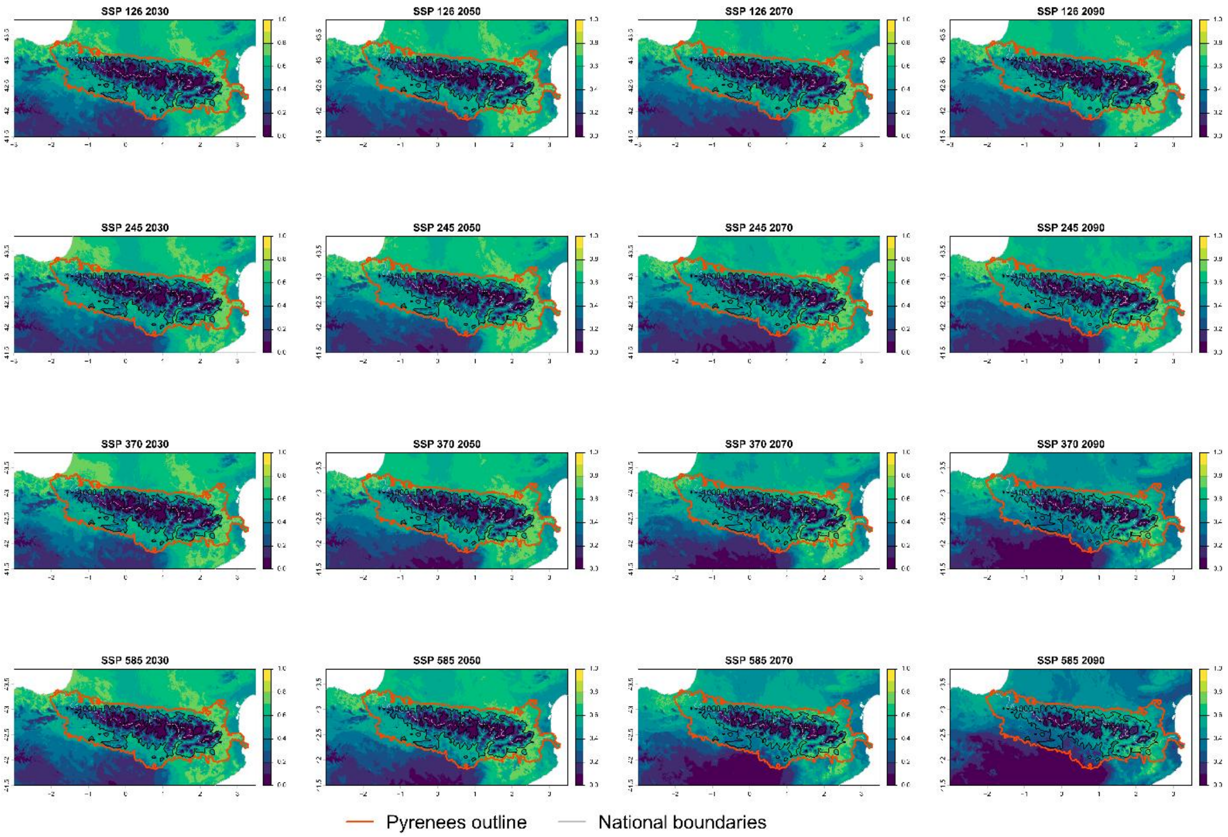

Current bioclimatic suitability in the Pyrenees for *Robinia pseudoacacia*

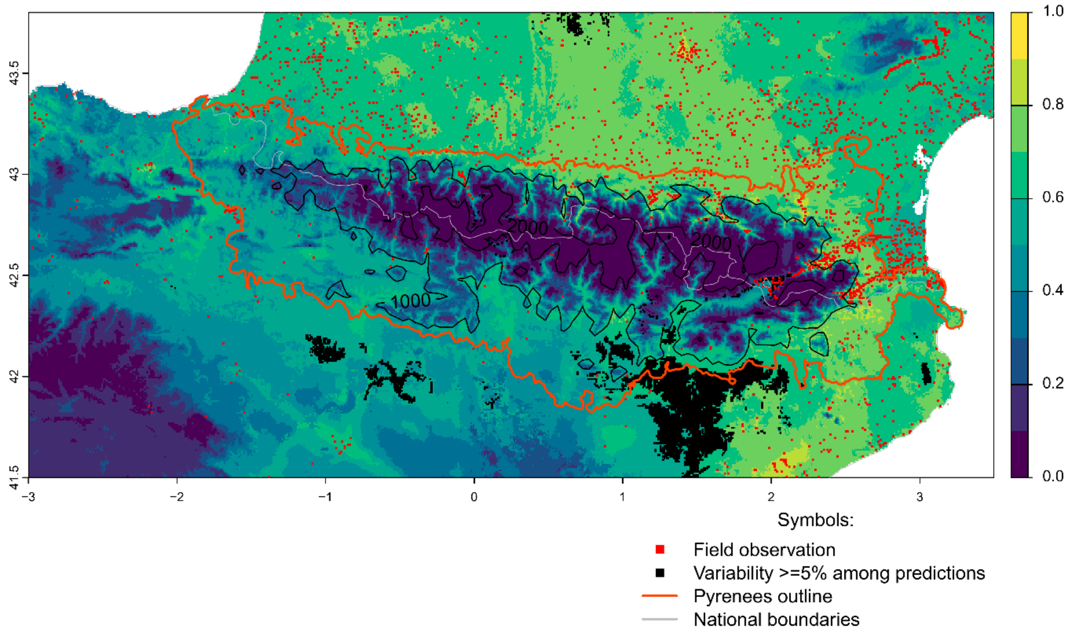


Future bioclimatic suitability in the Pyrenees by 2090 for *Robinia pseudoacacia*

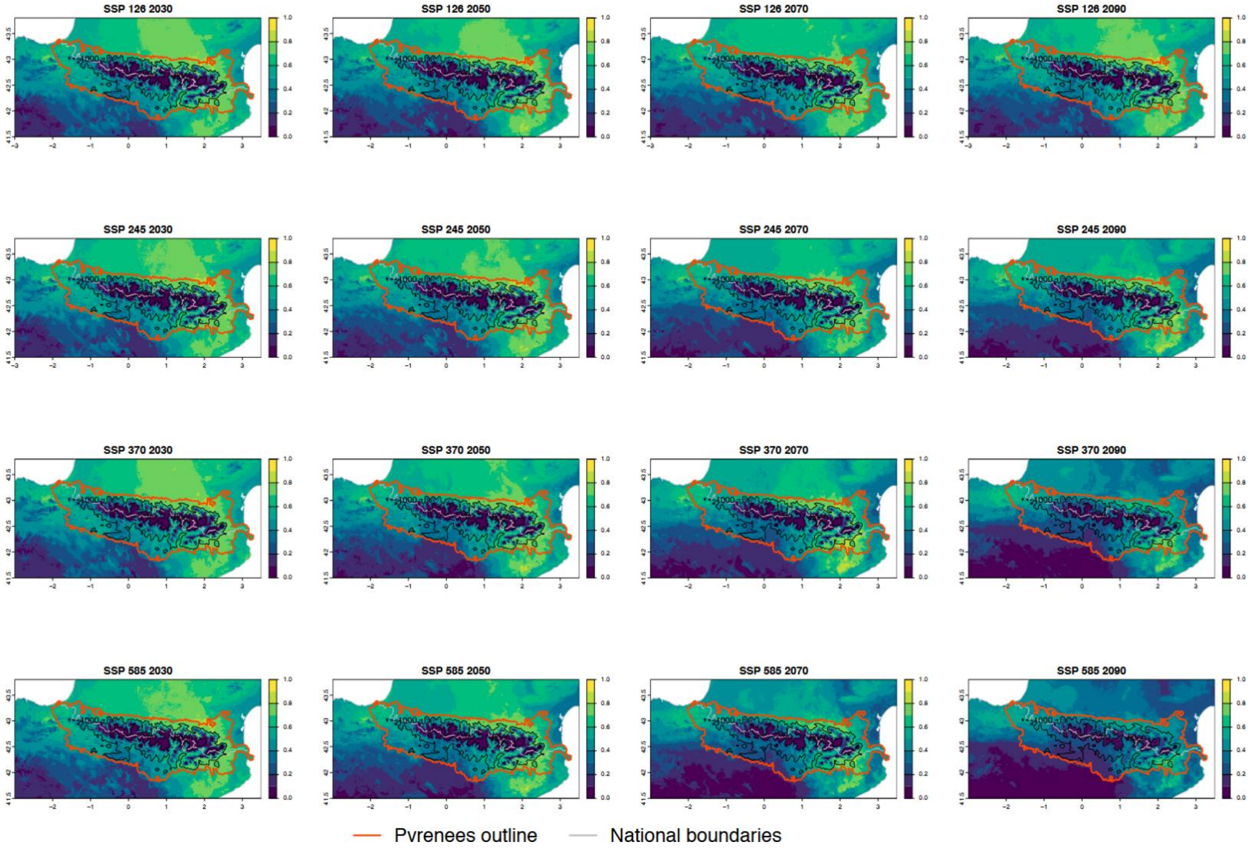

Current bioclimatic suitability in the Pyrenees for *Senecio inaequidens*

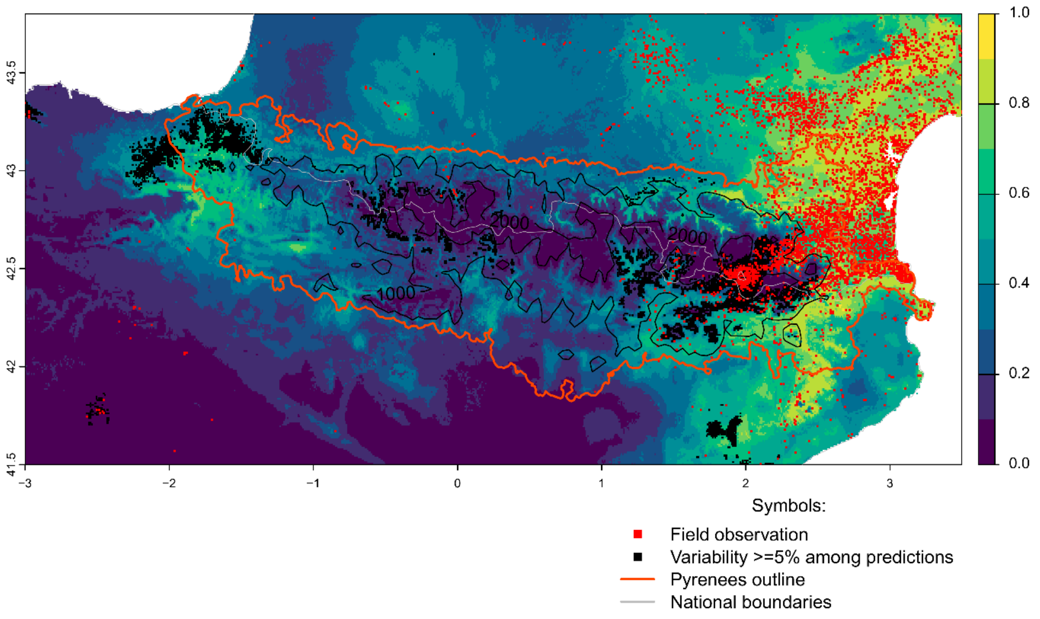


Future bioclimatic suitability in the Pyrenees by 2090 for *Senecio inaequidens*

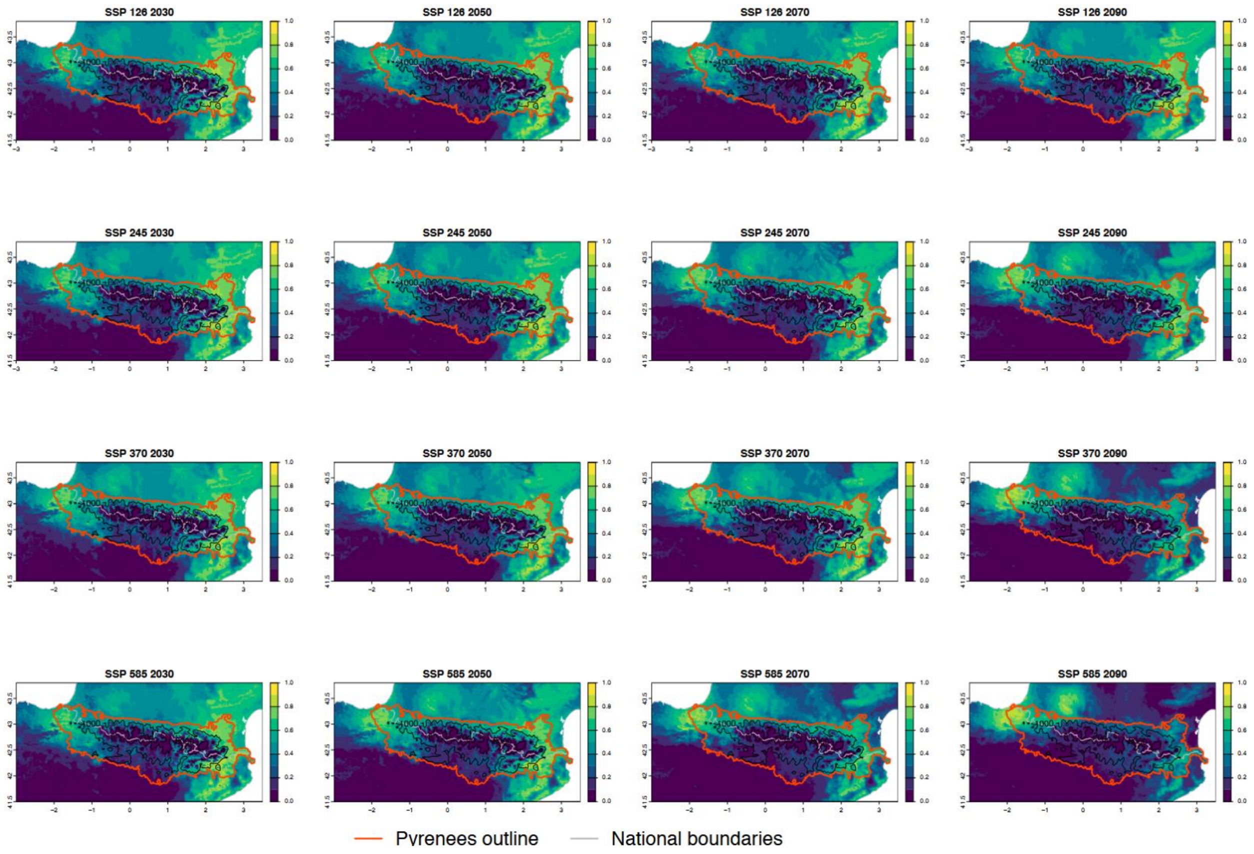

Current bioclimatic suitability in the Pyrenees for *Sicyos angulatus*

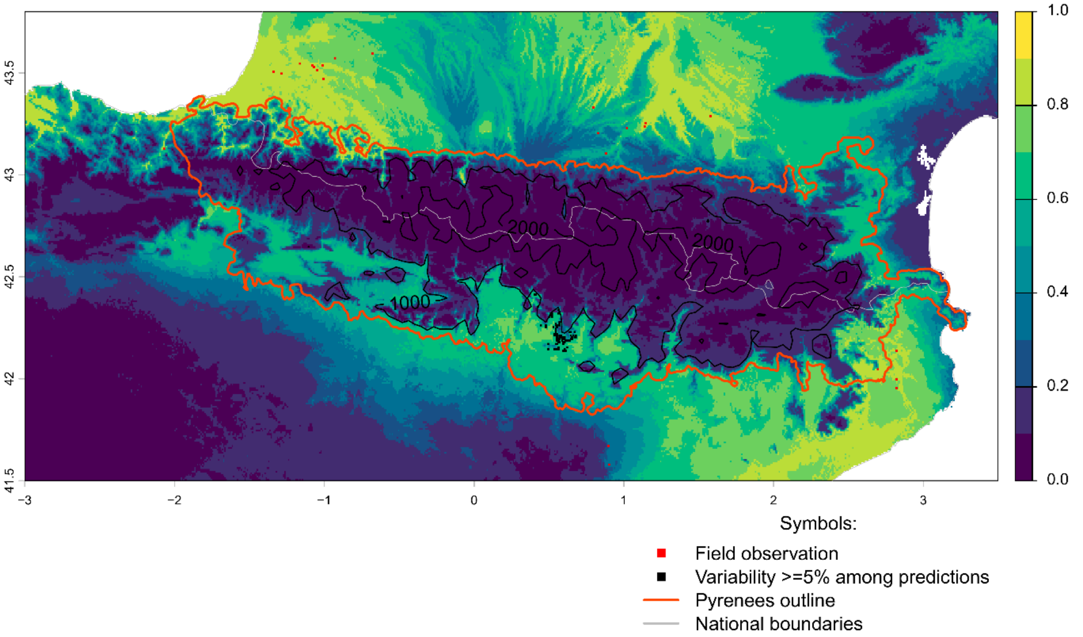


Future bioclimatic suitability in the Pyrenees by 2090 for *Sicyos angulatus*

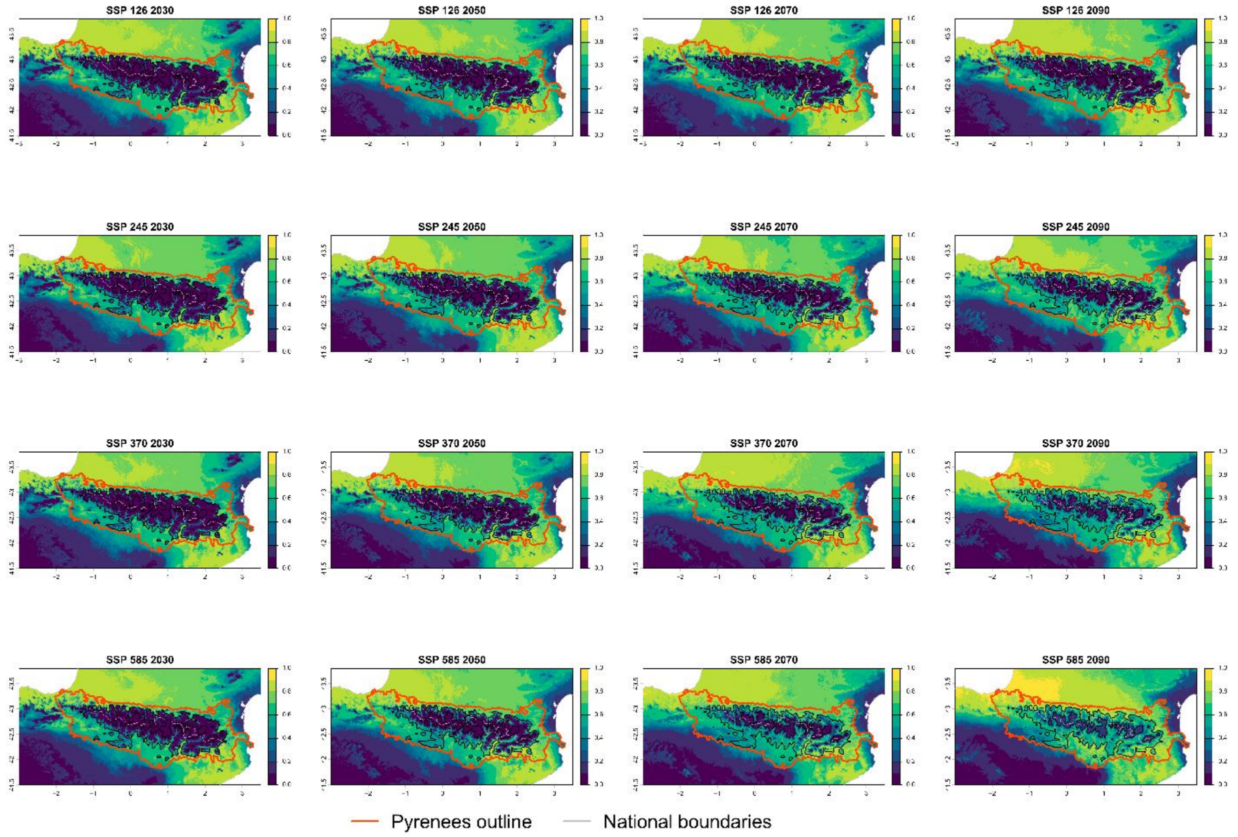

Current bioclimatic suitability in the Pyrenees for *Solidago canadensis*

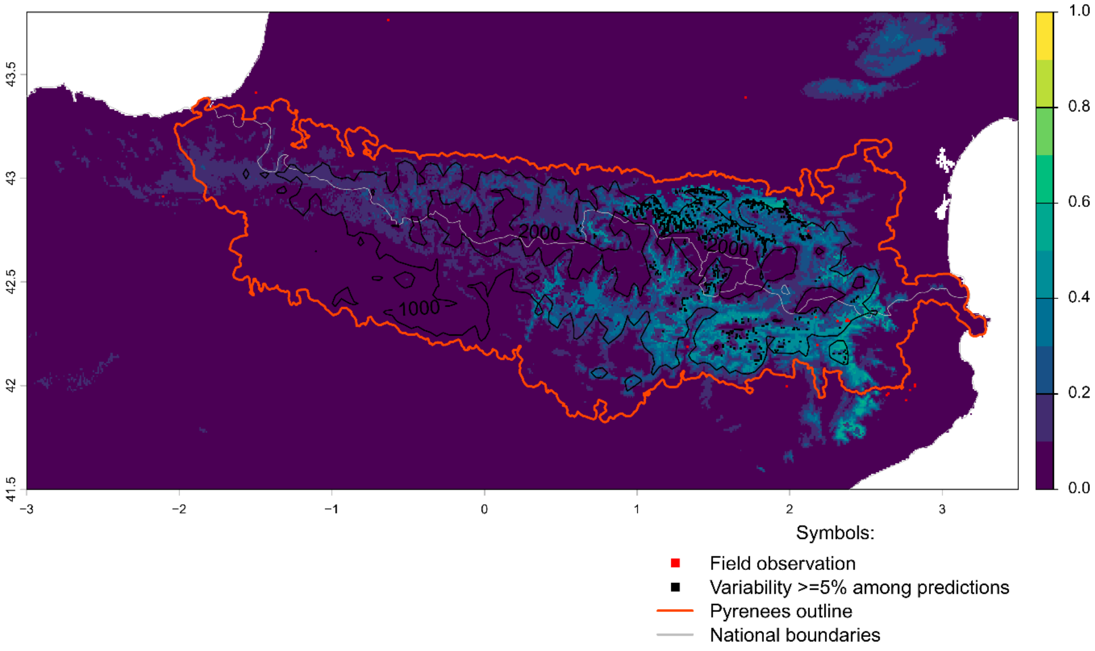


Future bioclimatic suitability in the Pyrenees by 2090 for *Solidago canadensis*

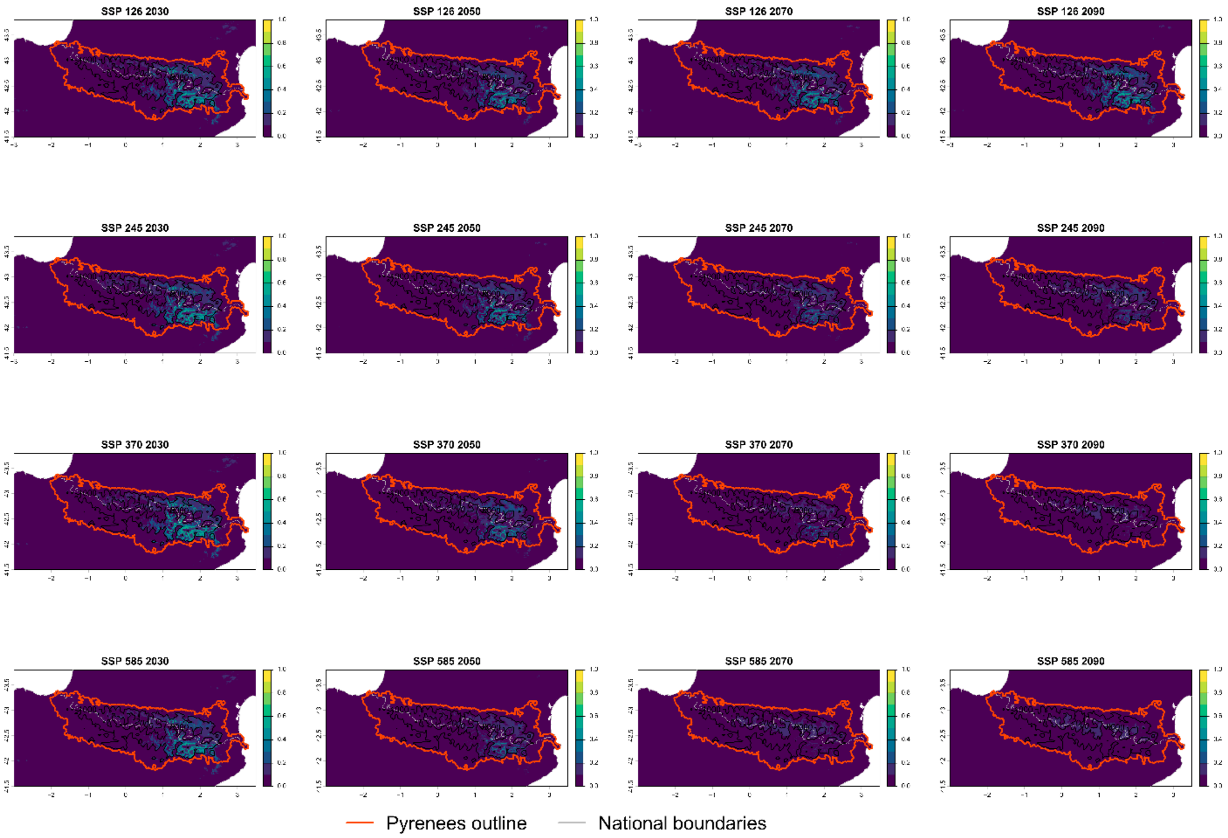

Current bioclimatic suitability in the Pyrenees for *Sorghum halepense*

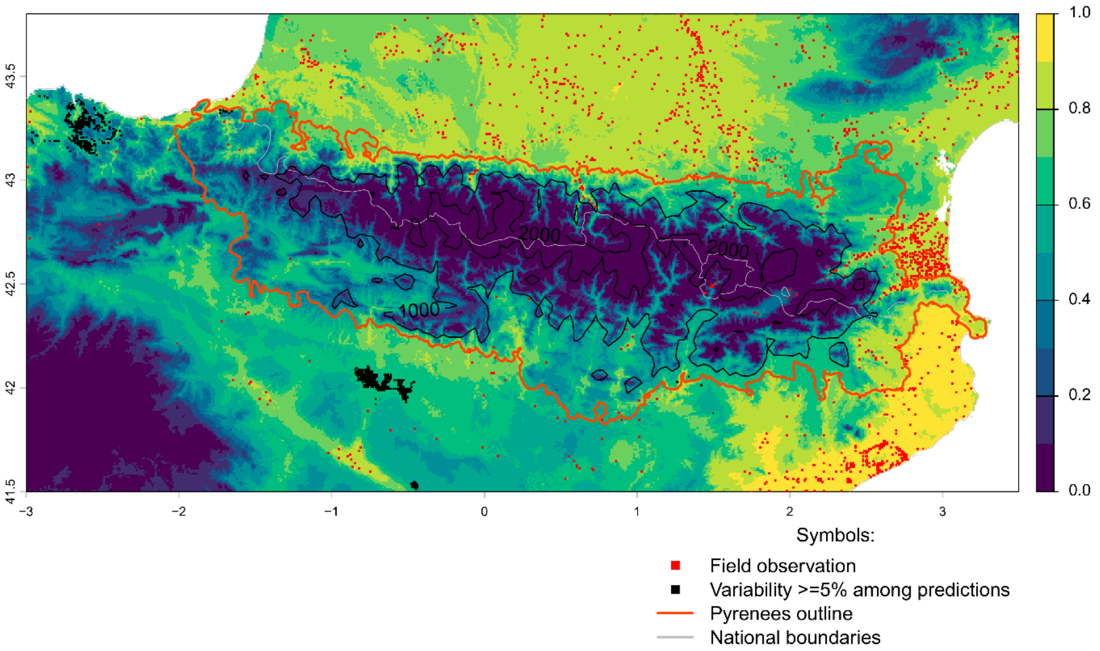


Future bioclimatic suitability in the Pyrenees by 2090 for *Sorghum halepense*

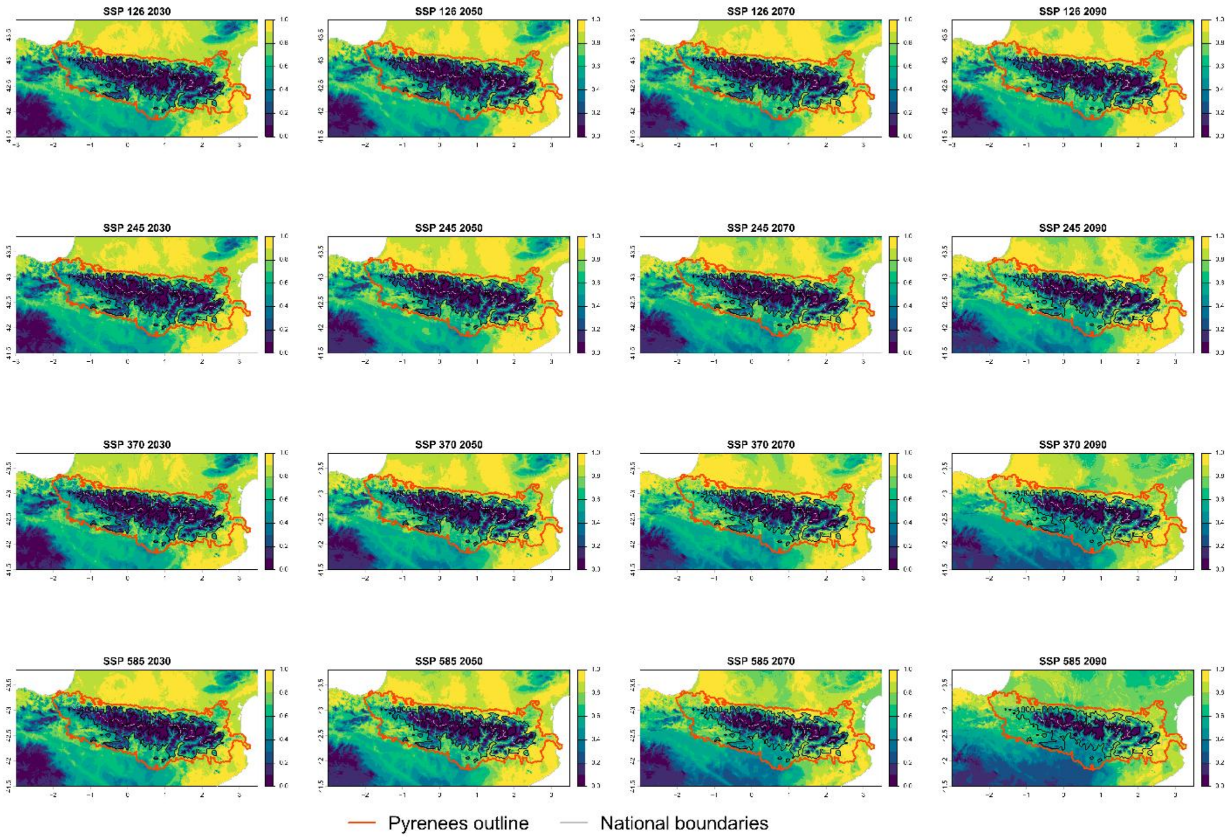

### Current bioclimatic suitability in the Pyrenees for *Sporobolus indicus*

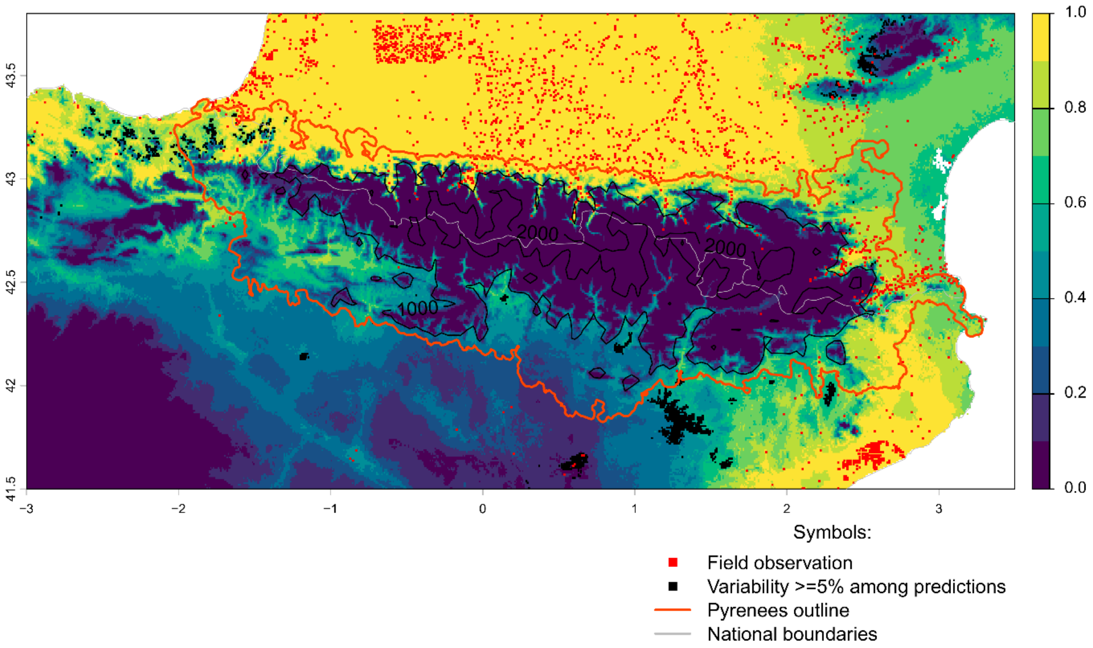


Future bioclimatic suitability in the Pyrenees by 2090 for *Sporobolus indicus*

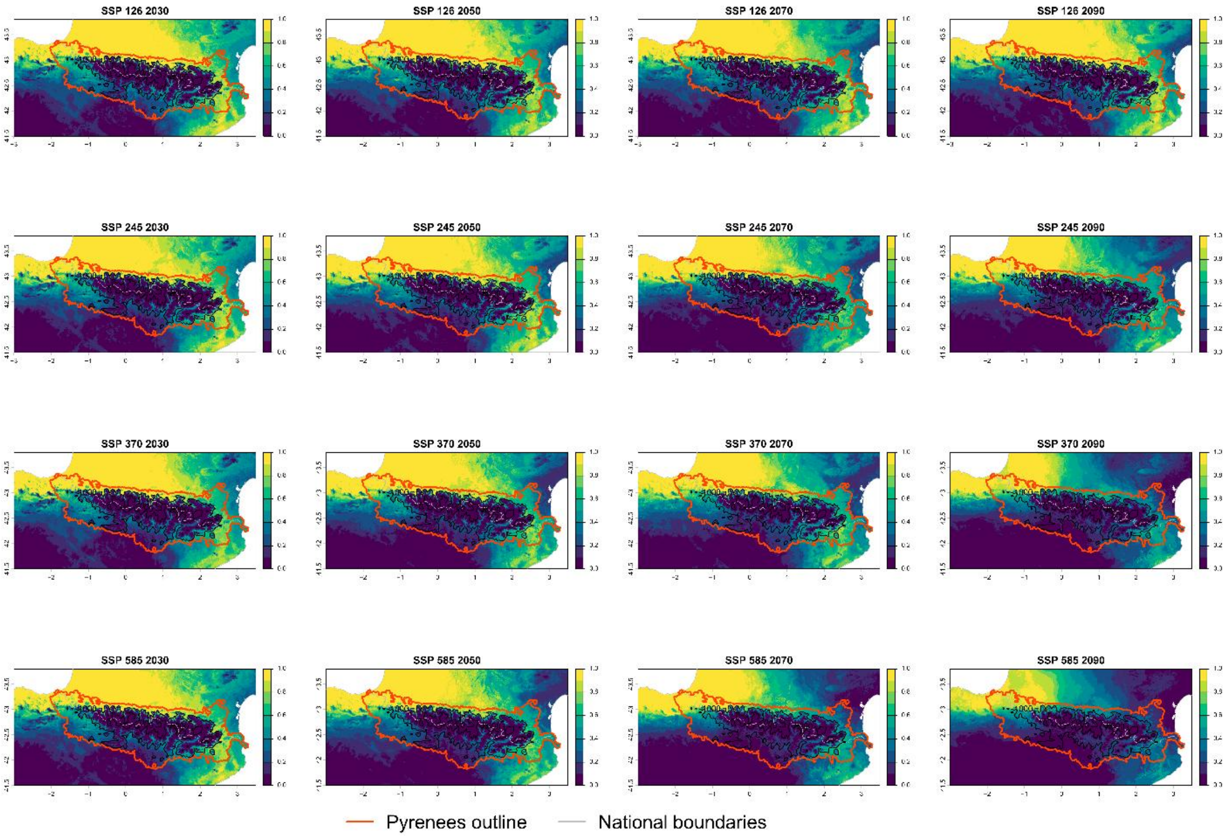

Current bioclimatic suitability in the Pyrenees for *Symphyotrichum pilosum*

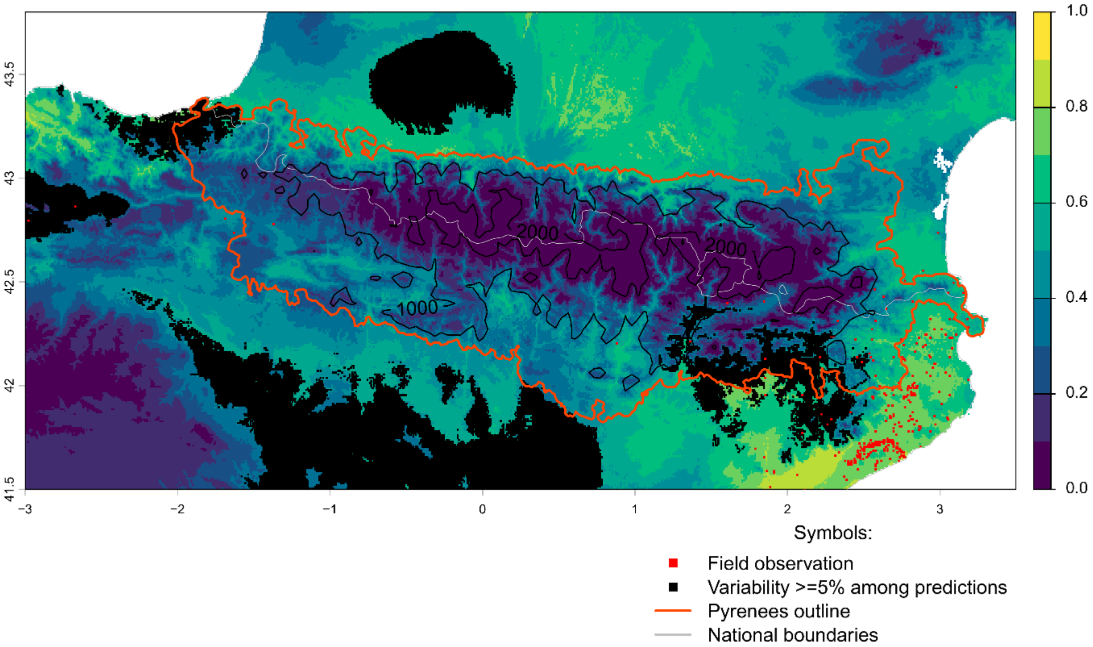


Future bioclimatic suitability in the Pyrenees by 2090 for *Symphyotrichum pilosum*

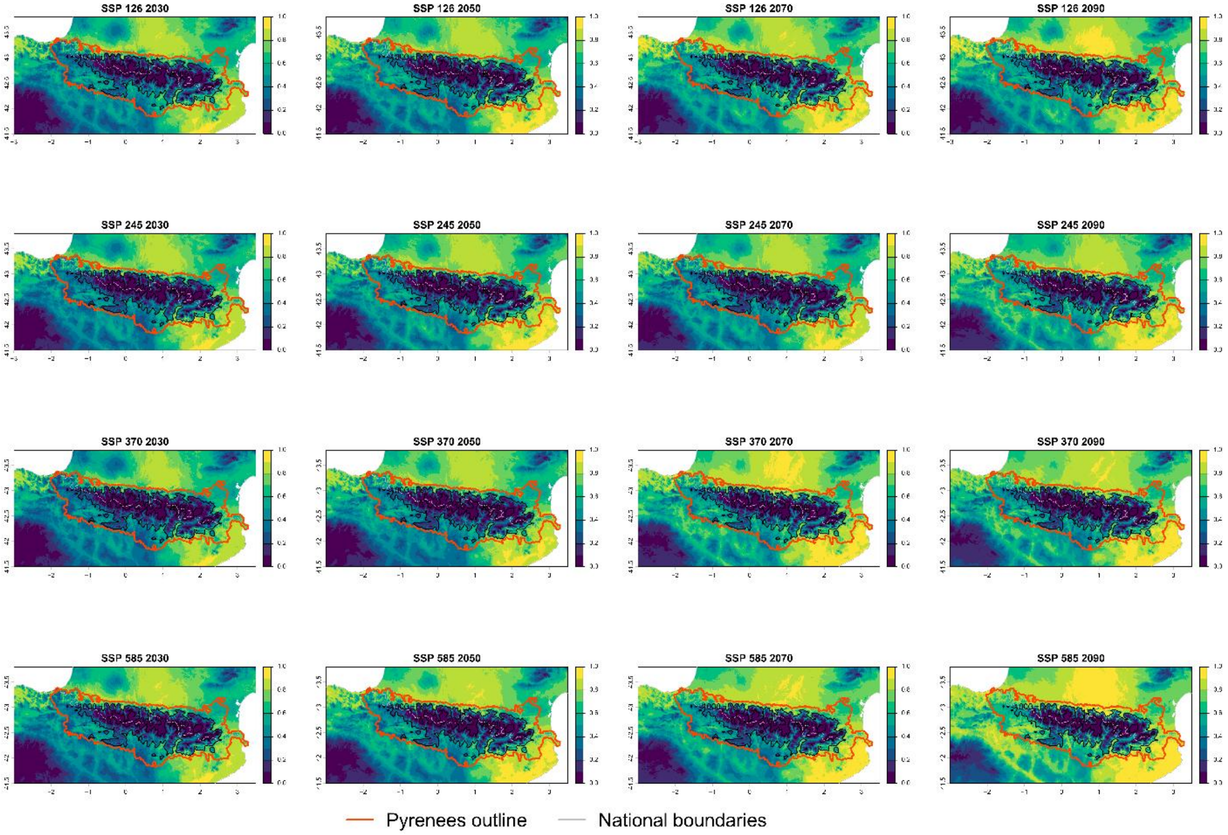

### Current bioclimatic suitability in the Pyrenees for *Symphyotrichum squamatum*

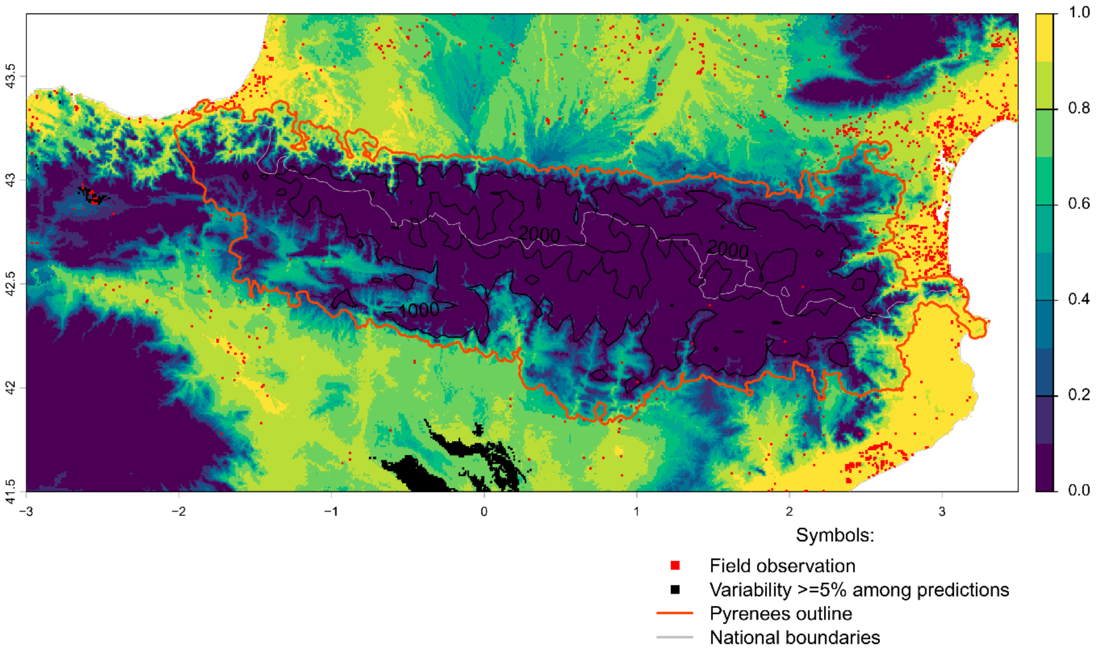


Future bioclimatic suitability in the Pyrenees by 2090 for *Symphyotrichum squamatum*

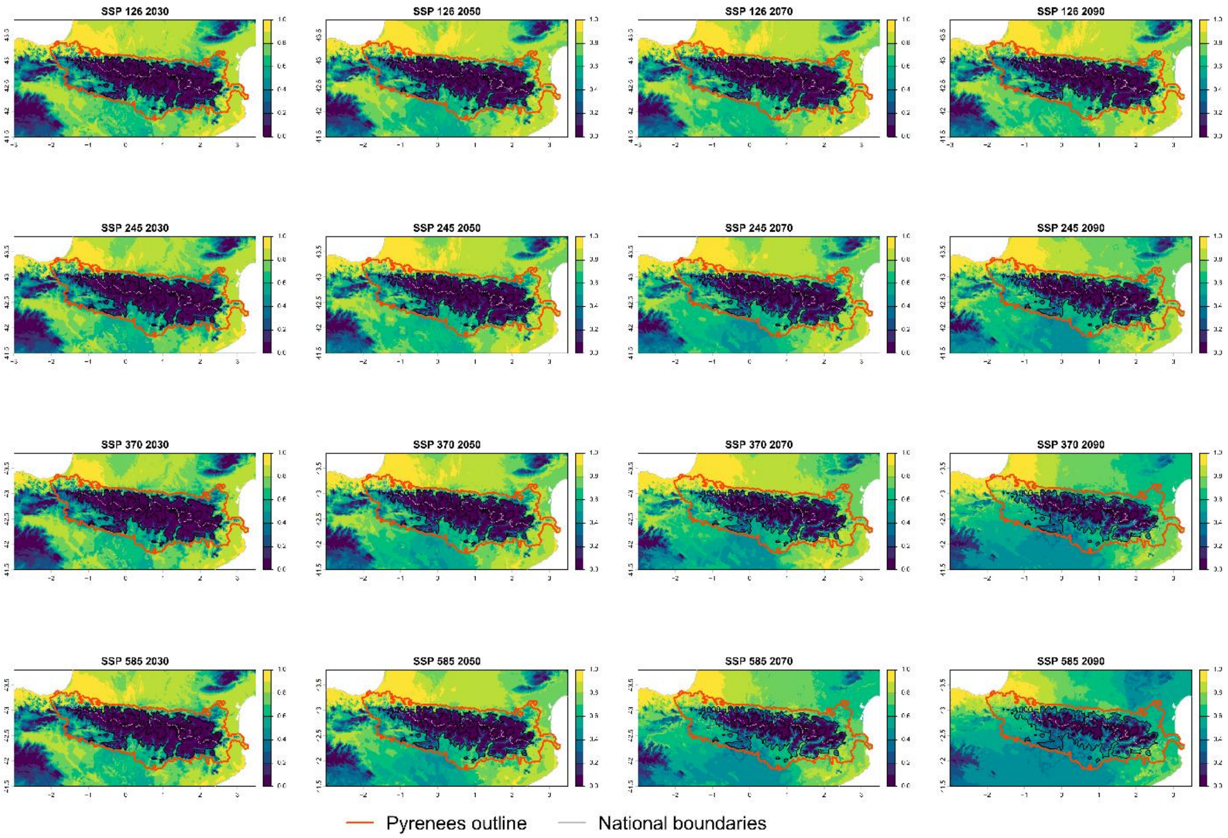

Current bioclimatic suitability in the Pyrenees for *Xanthium orientale*

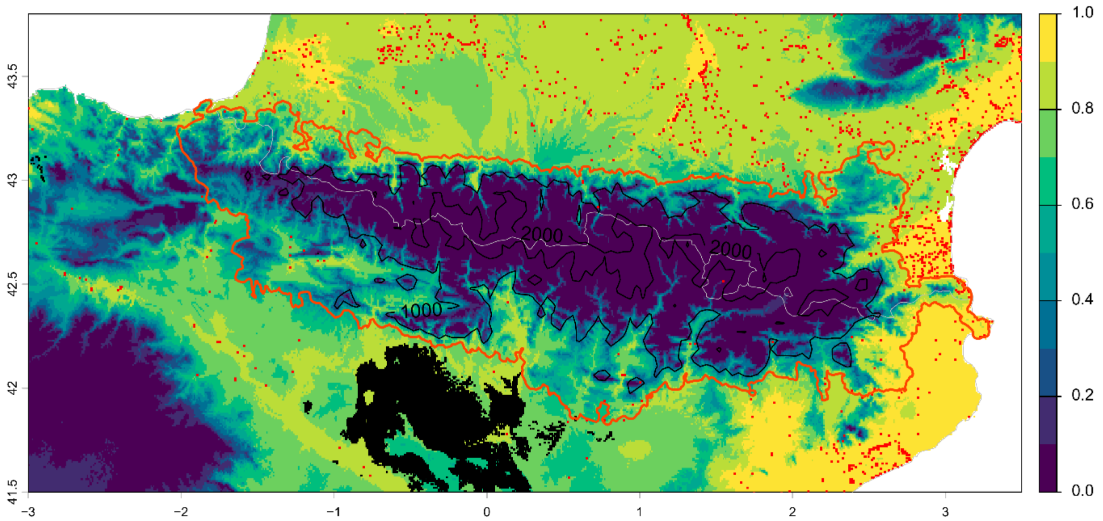


Symbols:

- Field observation
- Variability >=5% among predictions
- Pyrenees outline
- National boundaries

Future bioclimatic suitability in the Pyrenees by 2090 for *Xanthium orientale*

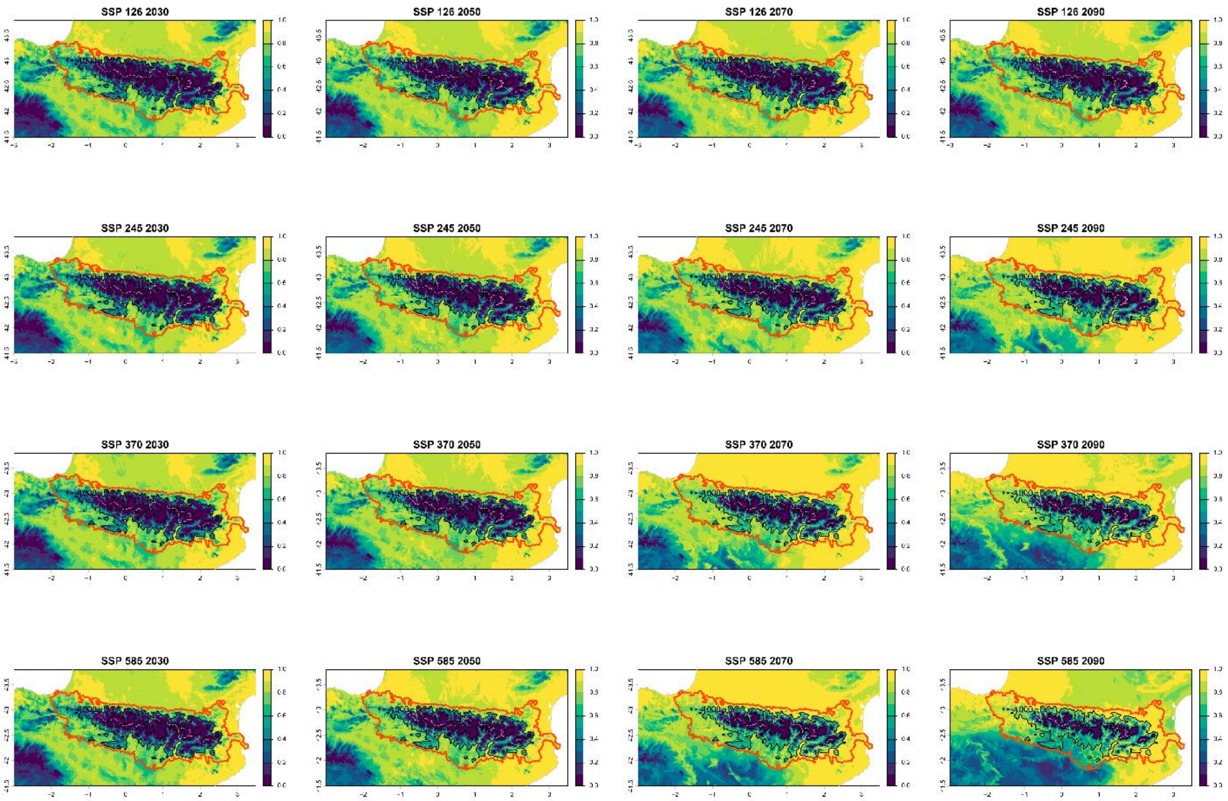


Pyrenees outline — National boundaries

**Appendix D: Longitudinal, latitudinal and elevational shifts in climatic suitability for invasive plant species in the Pyrenees under future climate scenarios for 2081-2100 period.**

For each species (rows), curves represent the difference between future and current climatic suitability calculated at the pixel level along each gradient, i.e. for all pixels sharing a given longitude (left), latitude (centre) or elevation class (right). Values therefore reflect the net gain or loss of suitability aggregated across pixels at each position along the gradient. Positive values indicate a gain in suitability, while negative values indicate a loss relative to present conditions (dashed horizontal line). Coloured lines correspond to the four SSP scenarios (SSP126, SSP245, SSP370, SSP585).

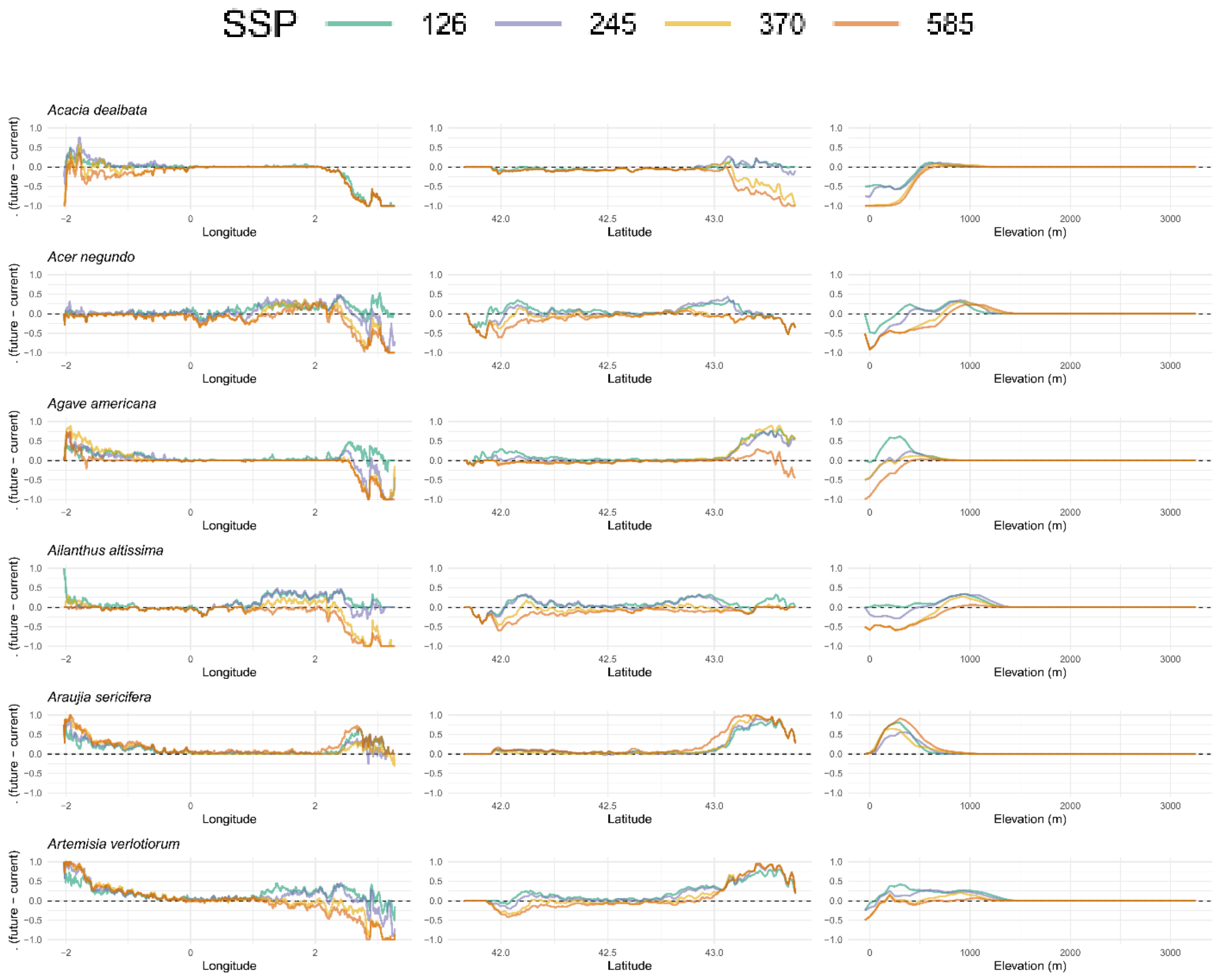

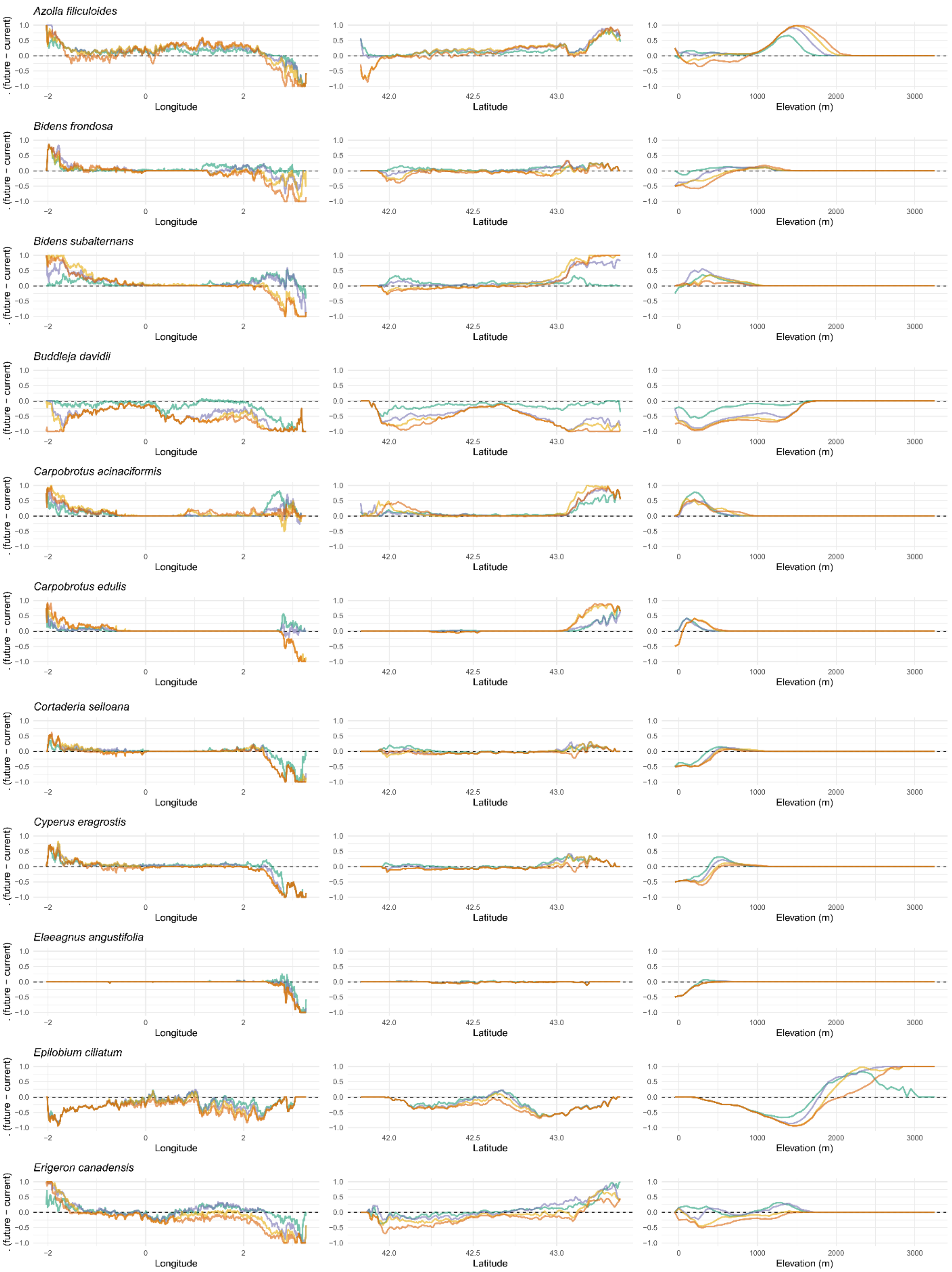

Azolla filiculoides
Bidens frondosa
Bidens subalternans
Buddleja davidii
Carpobrotus acinaciformis
Carpobrotus edulis
Cortaderia selloana
Cyperus eragrostis
Elaeagnus angustifolia
Epilobium ciliatum
Erigeron canadensis
. (future − current)
Longitude
Latitude
Elevation (m)

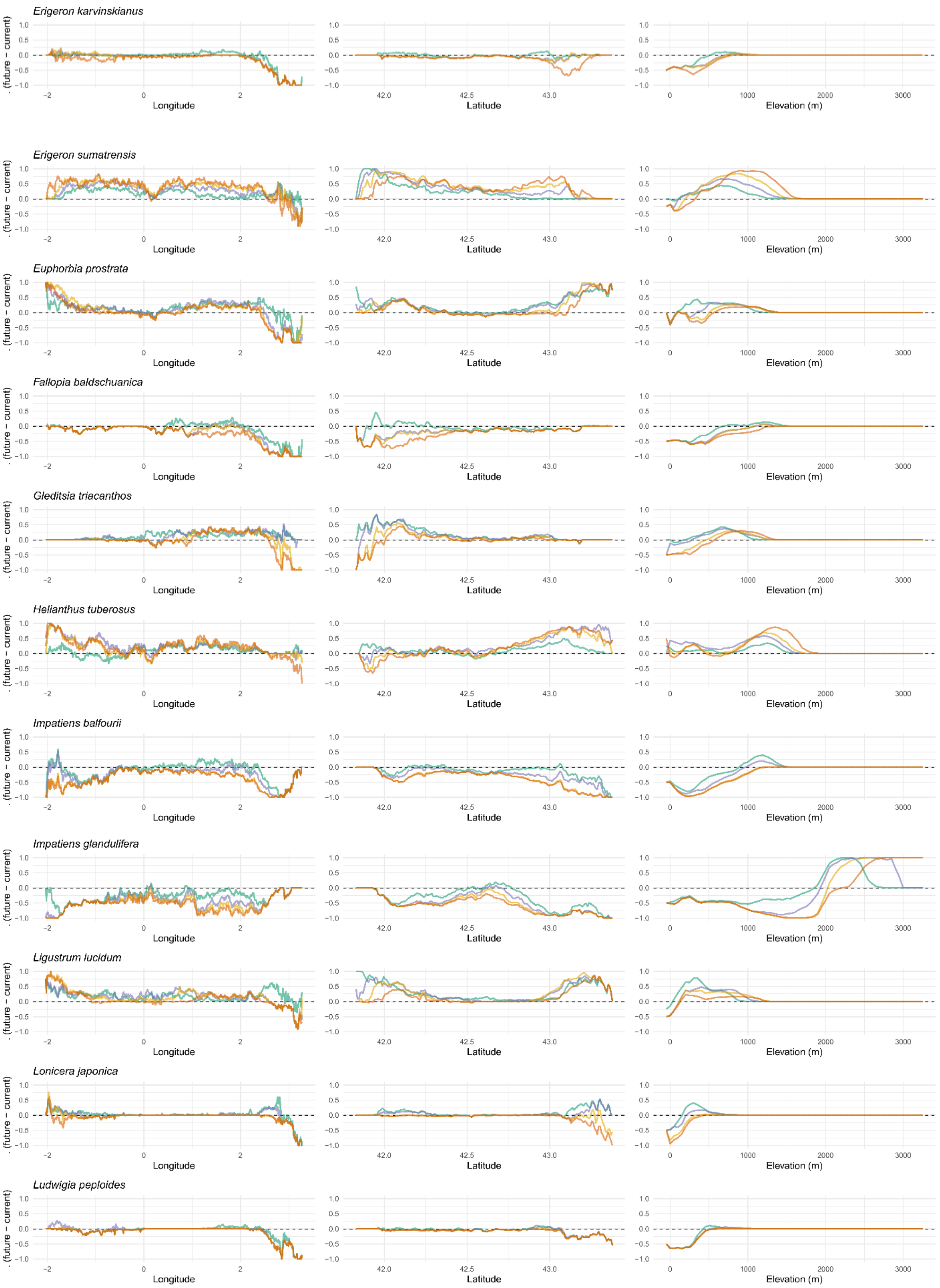
Erigeron karvinskianus
Erigeron sumatrensis
Euphorbia prostrata
Fallopia baldschuanica
Gleditsia triacanthos
Helianthus tuberosus
Impatiens balfourii
Impatiens glandulifera
Ligustrum lucidum
Lonicera japonica
Ludwigia peploides
(future – current)
Longitude
Latitude
Elevation (m)

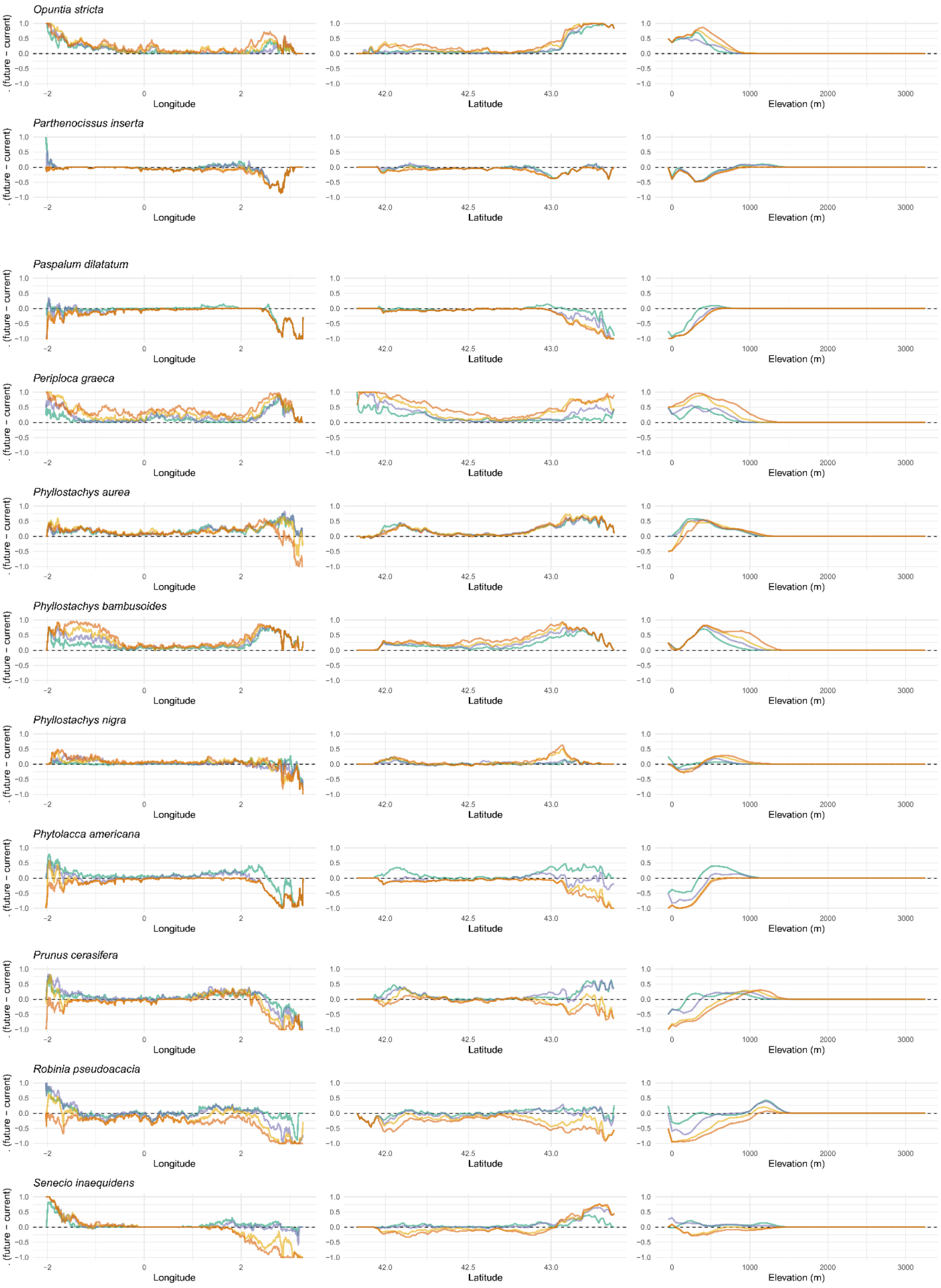
Opuntia stricta
Parthenocissus inserta
Paspalum dilatatum
Periploca graeca
Phyllostachys aurea
Phyllostachys bambusoides
Phyllostachys nigra
Phytolacca americana
Prunus cerasifera
Robinia pseudoacacia
Senecio inaequidens
(future − current)
Longitude
Latitude
Elevation (m)

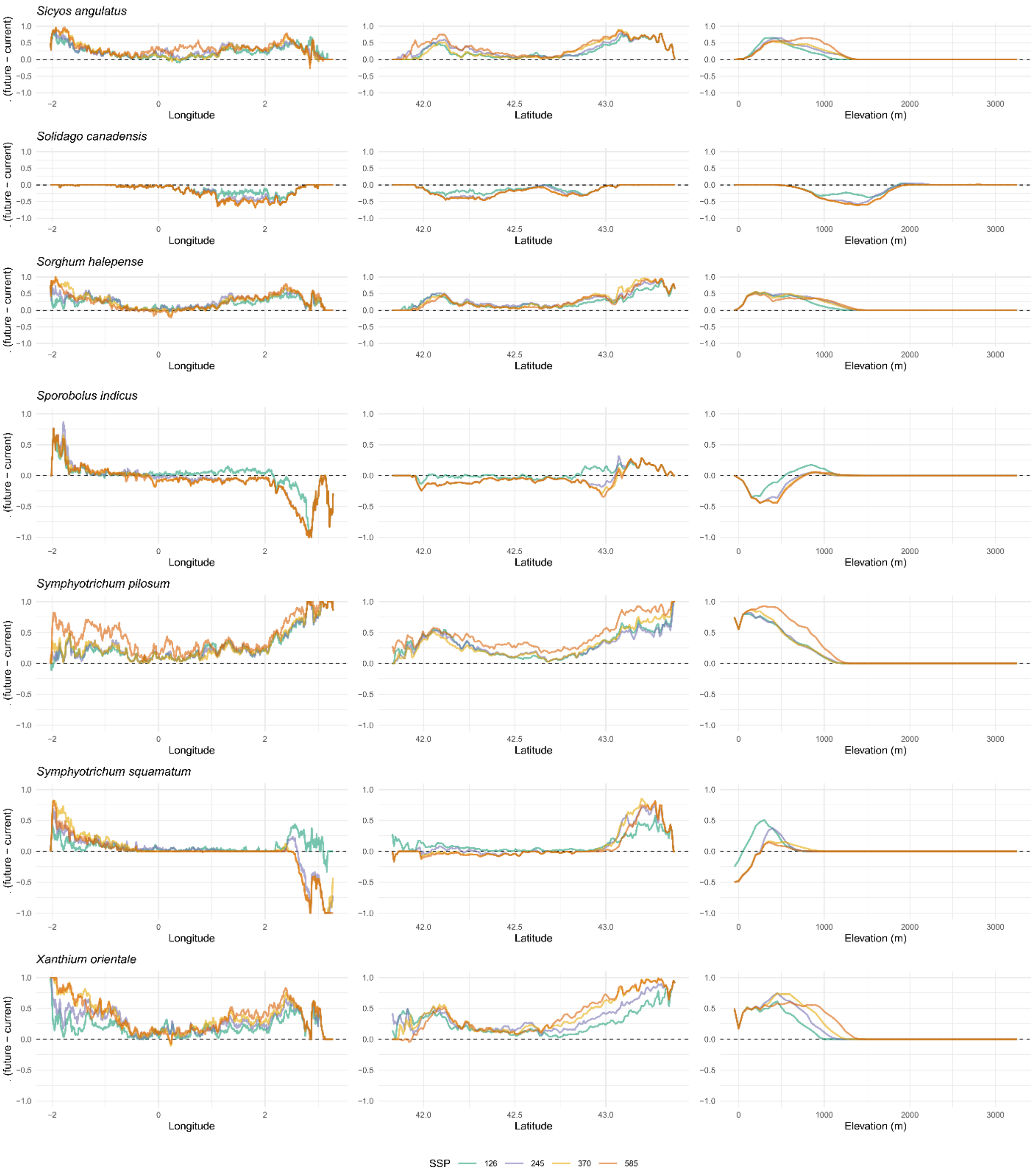
Sicyos angulatus
Solidago canadensis
Sorghum halepense
Sporobolus indicus
Symphyotrichum pilosum
Symphyotrichum squamatum
Xanthium orientale
(future − current)
Longitude
Latitude
Elevation (m)
SSP
126
245
370
585

**Appendix E: Species richness and density of invasive plant occurrence records across elevational belts in the Pyrenees**

**Figure E1: Species richness and occurrence-record density across elevational belts in the Pyrenees.**

Species richness (number of invasive plant species) and occurrence-record density (number of occurrence records per km²) are shown for four elevational belts corresponding to the main vegetation zones of the Pyrenees: foothills and lower montane (0–900 m), montane (901–1,800 m), subalpine (1,801–2,300 m), and alpine (> 2,300 m) (SAULE *ET AL.*, 2018). Metrics were computed using the full (non–spatially thinned) set of occurrence records. Bubble size is proportional to the total area (km²) of each elevational belt considered in the analysis.

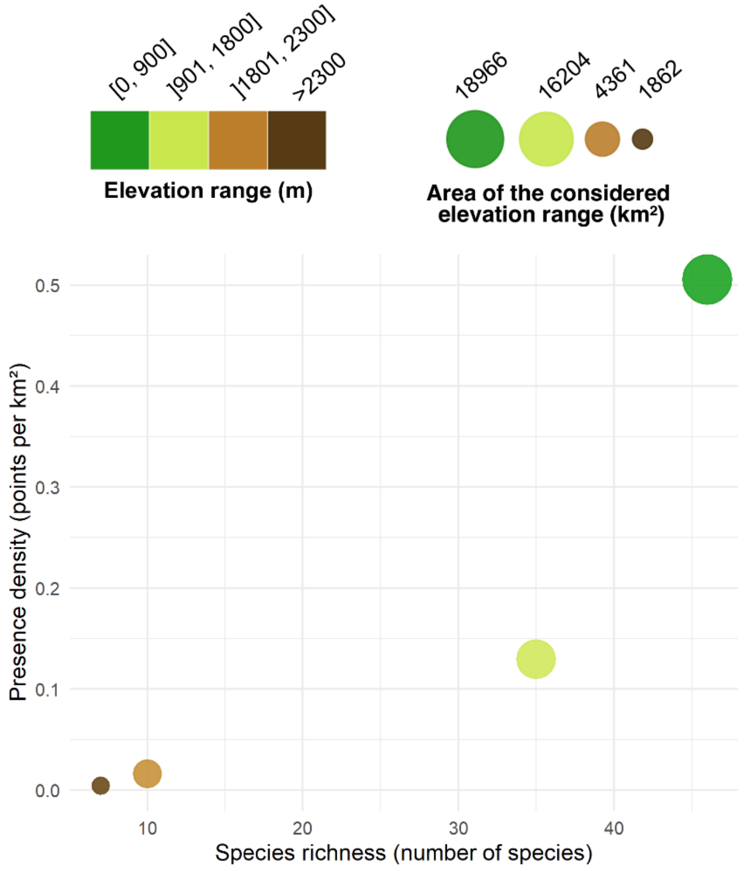

### Appendix F: Model performance evaluation

**Figure F1: Performance metrics of individual algorithms and ensemble models for invasive plant species included in this study.**

(a) Boxplot of three performance metrics, AUC ROC (discrimination), Boyce index (calibration), and sensitivity (classification), for the five modeling algorithms (GAM, GBM, GLM, MXT, RF) for all runs and species. Statistical significance is denoted as: *p < 0.05, **p <0.01, ***p < 0.001. (b) Performance of ensemble models across species, computed by averaging evaluation metrics from the validated models (Boyce index > 0.5) across the five algorithms used to build the ensemble consensus. Darker shades indicate higher performance.

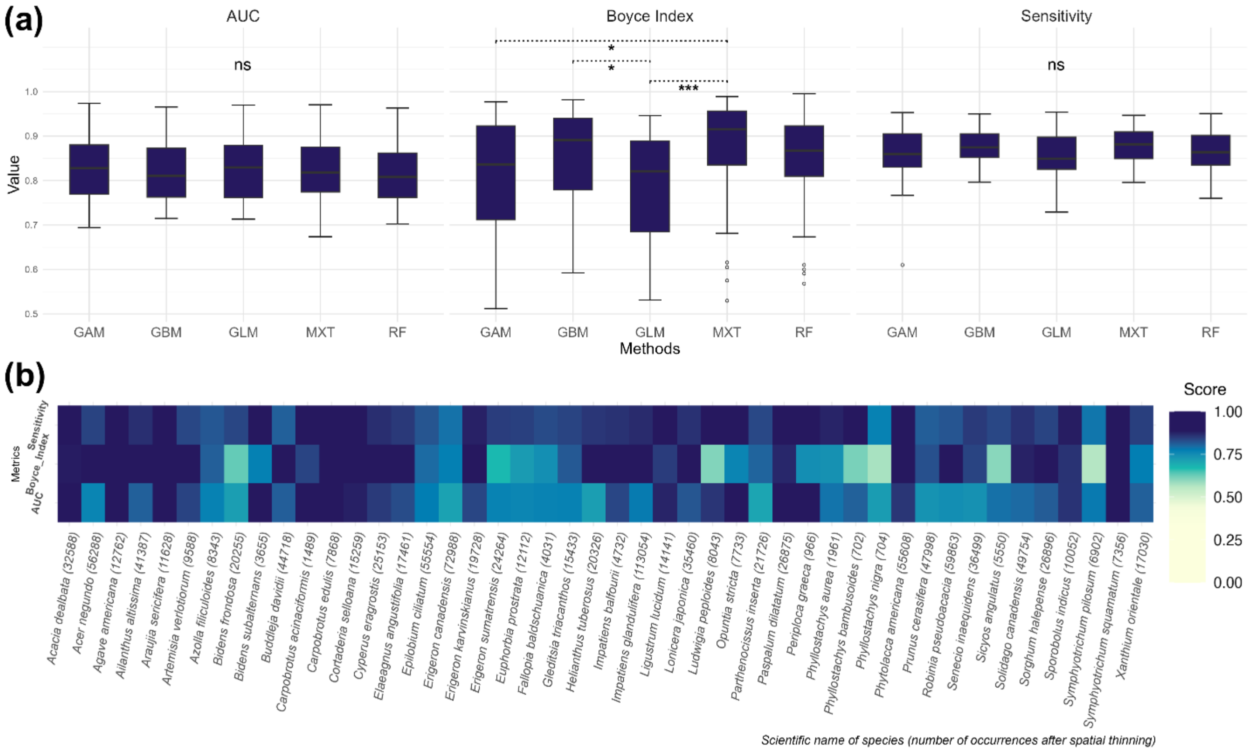
